# Substantial genetic mixing among sexual and androgenetic lineages within the clam genus *Corbicula*

**DOI:** 10.1101/590836

**Authors:** Vastrade Martin, Etoundi Emilie, Bournonville Thibaut, Colinet Mathilde, Debortoli Nicolas, Shannon M. Hedtke, Nicolas Emilien, Pigneur Lise-Marie, Virgo Julie, Flot Jean-François, Marescaux Jonathan, Van Doninck Karine

## Abstract

“Occasional” sexuality occurs when a species combines clonal reproduction and genetic mixing. This strategy is predicted to combine the advantages of both asexuality and sexuality, but its actual consequences on the genetic diversity and species longevity are poorly understood. Androgenesis, a reproductive mode in which the offspring inherits its entire nuclear genome from the father, is often reported as a strictly clonal reproductive mode. Androgenesis is the predominant reproductive mode within the hermaphroditic, invasive lineages of the mollusk genus Corbicula. Their ability to reproduce clonally through androgenesis has been determinant in their invasive success, having colonized during the 20th century American and European freshwater systems, where they became notorious invaders with a widespread, global distribution. However, in androgenetic Corbicula clams, occasional genetic mixing between distinct lineages has also been observed when the sperm of one lineage fertilizes the oocyte of another one. Because of these occasional introgressions, the genetic relationships between Corbicula species remained unclear, and the biogeographic origins of the invasive androgenetic lineages have been challenging to identify. To address these issues, we analyzed the patterns of allele sharing for several nuclear and mitochondrial molecular markers among Corbicula individuals collected across both the native and invasive range. Our results show the occurrence of an allelic pool encompassing all Corbicula freshwater species worldwide, including sexual and androgenetic ones, which highlights the substantial genetic mixing within this genus. However, the differences in allele sharing patterns between invasive lineages, and the low diversity within each lineage, suggest recent, distinct biogeographic origins of invasive Corbicula androgenetic lineages. Finally, the polyploidy, high heterozygosity, and hybrid phenotypes and genotypes found in our study probably originated from hybridization events following egg parasitism between distinct Corbicula lineages. This extensive cross-lineage mixing found in Corbicula may generate nuclear diversity in an otherwise asexually reproducing species.

## Introduction

Sexual reproduction with meiotic recombination is the most widespread reproductive mode among eukaryotes (Dimijian 2005; Barton 2009; Hartfield & Keightley 2012). Sex, through genetic reshuffling, is predicted to promote new allele combinations and to reduce genetic associations of linked loci, increasing the efficiency of selection and the purging of deleterious mutations (Lenormand *et al*., 2016; Bast *et al*. 2018). However, sex has many costs, including the two-fold cost of males (*i*.*e*., the reduced fecundity when half of the individuals does not produce offspring), the cost of meiosis (*i*.*e*., the reduced genetic contribution to the offspring), the cost of recombination (*i*.*e*., the possible loss of favorable associations between alleles), and the cost of mate searching and mating (Hartfield & Keightley 2012; Lehtonen *et al*. 2012; Liegeois *et al*. 2020). The prevalence of sex in eukaryotes is therefore considered a major paradox in evolutionary biology.

Besides sexual reproduction *sensu stricto*, including meiosis and syngamy, a diversity of asexual reproductive mechanisms exists in which oocytes appear to be produced through a modified meiosis or mitotically (Lenormand *et al*., 2016). In some organisms, asexual and sexual reproductions can alternate, referred to as occasional sexuality (or cyclical parthenogenesis in animals). In addition, there are several reproductive modes in which meiosis and fertilization still take place, but only the unreduced paternal or maternal genome is transmitted to the next generation. One such quasi-sexual reproductive mode is androgenesis, in which the offspring, after fertilization, inherits only the entire paternal nuclear genome (McKone & Halpern 2003, Schwander & Oldroyd, 2016). The maternal nuclear chromosomes are either discarded from the egg, or oocytes are non-nucleated prior to fertilization (reviewed in Schwander & Oldroyd, 2016). When androgenetic males or hermaphrodites use the eggs of other individuals to clone themselves, this reproductive mode is considered sexual parasitism or “egg parasitism” (Hedtke *et al*. 2008; Komaru *et al*. 2013, reviewed in Lehtonen *et al*. 2013), because one individual uses the female gamete of another individual to transmit his entire nuclear genome to the next generation, while the female host does not contribute with its nuclear genome. The processes that generate and maintain such a peculiar reproductive mode remain elusive. Moreover, androgenesis appears rare in nature or remains undetected, but is more prevalent in hermaphroditic species (McKone & Halpern 2003, Schwander & Oldroyd 2016). It has been observed as a main reproductive mode in only a few species, such as the conifer *Cupressus dupreziana* (Pichot *et al*. 2001), the ant *Wasmannia auropunctata* (Rey *et al*. 2013), and some stick-insect hybrid species (*e*.*g*., *Bacillus rossius-grandii*, Mantovani & Scali 2001, *Pijnackeria hispanica*, Milani *et al*. 2020). However, androgenesis is widespread in the clam genus *Corbicula*, with several species reproducing through obligate androgenesis (for a review on androgenetic species, see Schwander & Oldroyd, 2016).

The genus *Corbicula* is therefore one of the most suitable system to better understand the origin and maintenance of androgenesis and to study the genetic consequences of asexual reproduction while maintaining the sexual characteristics of meiosis and fertilization. Indeed, androgenesis can be considered as sexual from a functional and ecological perspective (eggs and sperm are required and fertilization does occur, albeit without karyogamy) but asexual from a genetic point of view (Pigneur *et al*. 2012). Moreover, within the clam genus *Corbicula*, besides hermaphroditic androgenetic species, dioecious sexual species are found, and it is the only metazoan clade known in which the hermaphroditic species reproduce through obligate androgenesis for which the underlying cytological mechanism is well described (Komaru *et al*. 2000; Ishibashi *et al*. 2002, 2003; Ishibashi & Komaru, 2006; Hotta & Komaru, 2018). Androgenetic *Corbicula* clams produce offspring after fertilization of a congeneric oocyte by a biflagellate, unreduced sperm. The oocyte continues meiosis, but the meiotic axis is parallel to the egg surface, while it is perpendicular to the cortex in the case of a canonical meiosis (Komaru *et al*. 2000, Figure 1). As a result, having both asters associated to the egg surface in androgenetic *Corbicula*, all the maternal nuclear chromosomes in meiosis I are discarded from the egg as two polar bodies, instead of one polar body in canonical meiosis (Komaru *et al*. 2000; Hotta & Komaru, 2018; Figure 1). The zygote, therefore, inherits the mitochondria from the oocyte and the nuclear genome from the sperm (reviewed in Pigneur *et al*., 2012). The unreduced, biflagellate sperm of androgenetic *Corbicula* is a presumed hallmark of their androgenesis, plausibly formed through an ameiotic process or modified meiosis in which ploidy reduction is absent (the details remain unknown). Besides the hermaphroditic androgenetic species, the sexual *Corbicula* species appear to be diploid, producing reduced, uniflagellate sperm (Glaubrecht *et al*., 2006; Obata *et al*. 2006).

**Figure 1:**
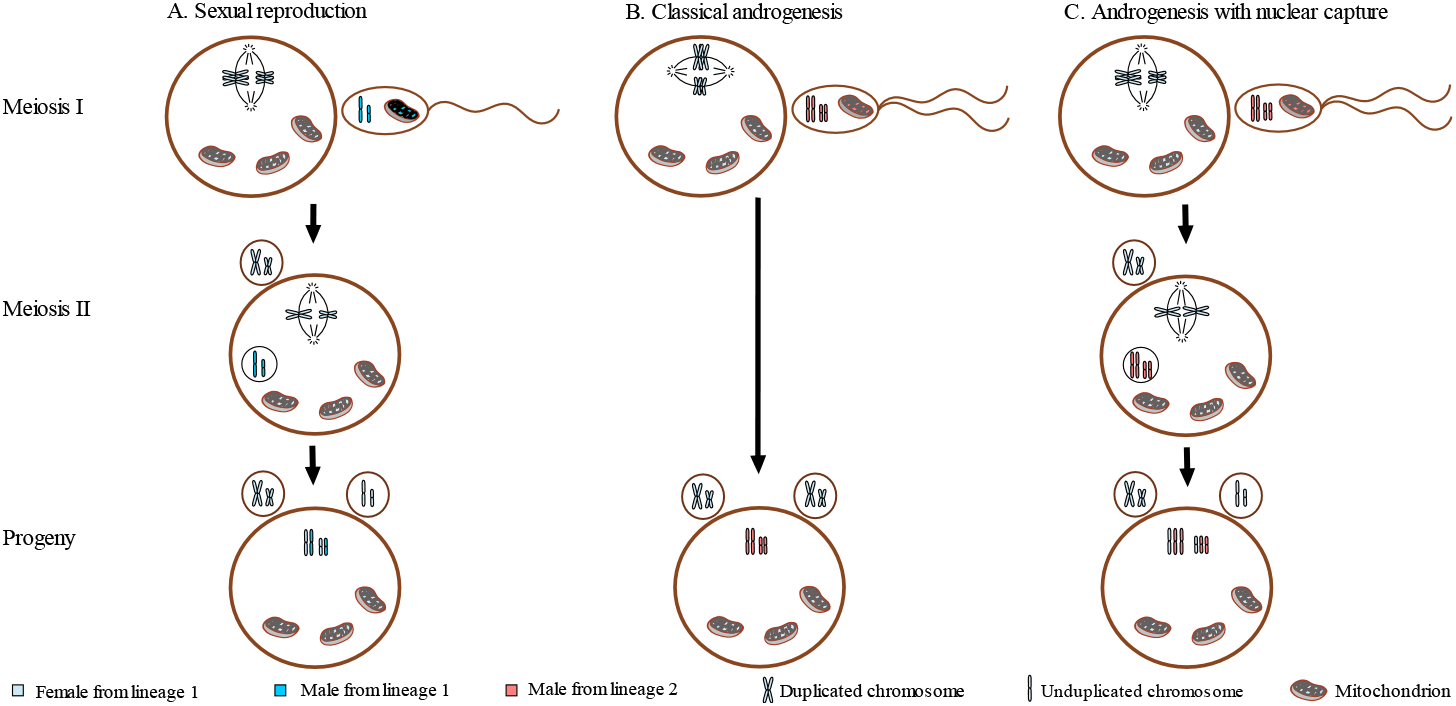
Possible outcomes of nuclear DNA through meiosis in *Corbicula* genus. Although this is variable in metazoans (Sagata 1996), fertilization occurs during meiosis I in *Corbicula* (Obata *et al.* 2006, Komaru *et al.* 2006). In all cases, the mitochondrial DNA is inherited maternally. **A**. In sexual species, a two-step meiosis leads to the production of a reduced haploid egg. Its fertilization by uniflagellate reduced sperm gives diploid progeny from both maternal (light blue) and paternal (dark blue) nuclear DNA (Obata *et al.* 2006). **B**. In androgenetic lineages, all the maternal nuclear DNA is usually extruded within polar bodies during meiosis I (meiotic axis parallel to the adjacent cortex, see Komaru *et al.* 1998), and the sperm is biflagellate and unreduced. The progeny is thus diploid from paternal nuclear DNA only. If this occurs between two distinct lineages (here represented in blue and red), this leads to a mitochondrial mismatch. **C**. In androgenetic lineages, the maternal DNA exclusion can also be incomplete, producing a triploid progeny with both maternal and paternal nuclear DNA (Komaru *et al.* 2006). If occurring between distinct lineages (blue and red), this leads to hybridization. For clarity, only two chromosome sets are represented (different size and edge thickness) and the gamete sizes are not to scale.

In *Corbicula*, androgenetically produced offspring are formed through self-fertilization or outcrossing (Kraemer *et al*., 1986). When outcrossing occurs between distinct species, the nuclear genome of one species becomes associated with the mitochondria of another one, resulting in cytonuclear mismatches (Figure 1b) (Hedtke & Hillis 2011) and increased diversity of mitochondrial sequences within a species (Lee *et al*., 2005; Hedtke *et al*., 2008; Pigneur *et al*., 2012). Moreover, in some cases, maternal nuclear chromosomes may be partially retained and both maternal and paternal nuclear chromosomes are inherited by the offspring, eventually resulting in an increased ploidy (see Komaru *et al*., 1997; Park *et al*., 2000; Qiu *et al*., 2001). Such nuclear capture or hybridization events, if they occur among different *Corbicula* species, could in addition be a cause of genetic divergence between androgenetic species (Pigneur *et al*. 2014), generating nuclear diversity in otherwise asexually reproducing species (Figure 1c) (Hedtke *et al*., 2011). This process has been recorded in *Corbicula* (Komaru *et al*., 2006) and was reported as “rare” (Hedtke *et al*. 2011), although its frequency has never been thoroughly quantified. In addition, because these introgression events appear to occur between distinct *Corbicula* species, hybrid genotypes might be the cause of observed intermediate phenotypes (Pfenninger *et al*. 2002; Pigneur *et al*. 2014) and polyphyletic gene trees that fail to resolve taxonomy in the genus *Corbicula*. The taxonomy of invasive *Corbicula* species is particularly contentious because it has been only based on morphology and mitochondrial COI barcodes with several species’ names being given including *C. fluminea, C. fluminalis, C. largillierti, C. fluviatilis*, and *C. leana*, while lacking the nuclear genotype information (*e*.*g*., Coughlan *et al*. 2019; Crespo *et al*. 2015; Gomes et al. 2020; López-Soriano *et al*. 2018; Minchin 2014; Patrick *et al*. 2017; Reyna *et al*. 2018; Voroshilova *et al*. 2021). Some of these species’ names are likely to be synonymous and therefore using this species-based nomenclature can confuse lineage identification between studies.

*Corbicula* clams originated from Africa, Asia, Australia, and the Middle East (Araujo *et al*. 1993). In some of these locations, both sexual and androgenetic species co-occur but the reproductive mode of many native lineages remains unknown. The well-described sexual dioecious species have very restricted geographic distribution: *Corbicula sandai* is endemic to Lake Biwa in Japan, *Corbicula japonica* is restricted to East Asian brackish water, and a few sexual *Corbicula* species are from Indonesia (Glaubrecht *et al*. 2006; Yamada *et al*., 2014, Bespalaya *et al*. 2020). These species were described as sexual based on sperm morphology (Konishi *et al*. 1998, Hedtke *et al*. 2011, Pigneur *et al*. 2014). The androgenetic *Corbicula* species have a more widespread distribution across Asia. While the sexual species *C. sandai* and *C. japonica* have been identified as distinct species based on reproductive, morphological and genetic parameters (Pigneur *et al*. 2014), the species delimitation of native androgenetic *Corbicula* is also uncertain. For example, Komaru *et al*. (2012) showed a sister relationship between native, androgenetic *C. leana* and sexual *C. sandai* in Lake Biwa (Japan), but in a separate study *C. leana* clustered instead with androgenetic *C. fluminea* (Komaru *et al*. 2013). We therefore refer in our previous studies and here to androgenetic *Corbicula* lineages or forms instead of species (as in Pigneur *et al*. 2014).

All of the invasive *Corbicula* lineages found in Europe, North America, and South America appear to be exclusively androgenetic (Lee *et al*., 2005; Hedtke *et al*., 2008; Pigneur *et al*., 2014; Gomes *et al*., 2016). Based on the COI barcode and confirmed by nuclear microsatellite data (Pigneur *et al*. 2014), four distinct androgenetic and invasive *Corbicula* lineages have been described: form A/R being widespread and cosmopolitan in America and Europe (COI haplotype FW5, Table S1), form B only known from America (COI haplotype FW1, Table S1), form C/S found in America and Europe (COI haplotype FW17, Table S1), and form Rlc only described in Europe (COI haplotype FW4, Table S1) (Siripattrawan *et al*. 2000; Pfenninger *et al*. 2002, Lee *et al*. 2005; Hedtke et al. 2008; Marescaux *et al*. 2010, Pigneur *et al*. 2014). Peñarrubia *et al*. (2017) considered Rlc and B as one form, because their COI haplotypes differ by only one mutation, but the nuclear data showed significant differences between forms Rlc and B (Pigneur *et al*. 2014). More recently, a new invasive form D was described in America based on mitochondrial and nuclear genetic data (Tiemann *et al*. 2017; Haponski and Foighil 2019). Within all these invasive forms virtually no genetic diversity was found over their entire invasive range at the nuclear and mitochondrial markers studied (Hedtke *et al*., 2008; Pigneur *et al*., 2014; Gomes *et al*., 2016; Haponski & Foighil, 2019). When distinct invasive forms were found in sympatry, cytonuclear mismatches and hybrids (resulting from maternal genome retention) have been detected (Lee *et al*., 2005; Hedtke *et al*., 2008; Pigneur *et al*., 2014; Peñarrubia *et al*., 2017; Tiemann *et al*., 2017; Bespalaya *et al*., 2018; Haponski & Foighil, 2019; Sano *et al*. 2020). For example, out of twelve sampled locations in Europe where form A/R occurs in sympatry with another invasive form (C/S or Rlc), cytonuclear mismatches were detected at 8 locations and hybrids at 2 locations (Pigneur *et al*. 2014; Etoundi, pers. obs.). However, a more thorough quantification would be required.

The low diversity and the invasion by only a few *Corbicula* lineages in America and Europe are probably the consequence of a recent introduction by only few hermaphroditic individuals (920’s in merica and 980’s in Europe; Mouthon, 1981, McMahon, 1892, Pigneur *et al*., 2014; Gomes *et al*., 2016; Bespalaya *et al*., 2018) and their ability to reproduce through androgenesis and to self-fertilize (Kraemer *et al*., 1986). This indeed facilitates their rapid spread because only one hermaphroditic individual is required to establish a new population. In addition, since the descendants are derived only from the unreduced sperm, there is no genetic mixing (except if different lineages occur in sympatry), no loss of heterozygosity with time (except if recombination takes place), and no inbreeding depression in the invasive populations (Schwander & Oldroyd, 2016). In contrast, androgenetic *Corbicula* from the native range have been characterized by high genetic diversity comparable to that of sexual *Corbicula* lineages (Pigneur *et al*., 2014). This high native diversity could be the consequence of genetic captures by co-occurring androgenetic lineages or multiple transitions to asexuality from sexually reproducing populations (*e*.*g*. in ostracods, Bode *et al*., 2010; in *Timema* stick insect, Schwander & Crespi, 2009; van der Kooi & Schwander, 2014), or both.

The non-monophyly of androgenetic *Corbicula* lineages could be explained either by a single origin of androgenetic *Corbicula* lineages followed by independent nuclear capture events between sympatric lineages, or by multiple, independent origins from diverse sexual lineages followed by nuclear captures (Hedtke *et al*., 2011; Pigneur *et al*. 2011a, 2012, 2014; Gomes *et al*. 2016; Peñarrubia *et al*., 2017). Hedtke *et al*. (2011) proposed, based on phylogenetic reconstructions using nuclear markers, a common origin for most invasive androgenetic *Corbicula* lineages from a *C. sandai* ancestor but with a possible second origin of androgenesis for the invasive form C/S because it grouped with *Corbicula moltkiana* from Indonesia across their phylogenies. However, in the COI phylogenies of Pigneur *et al*., form C/S clustered with *C. australis* (2011a) or with *C. fluminalis africana* (2014). These phylogenetic discordances due to nuclear and mitochondrial captures among *Corbicula* lineages have until now hampered the investigation of phylogenetic relationships between androgenetic and sexual *Corbicula* (see Hedtke & Hillis 2011; Hedtke *et al*. 2011; Pigneur *et al*., 2012, 2014).

In the present work, multiple nuclear markers combined with the COI mitochondrial barcode were used to study the genetic relationships between *Corbicula* lineages from the native and invasive distributions, including different reproductive modes. Because phylogenetic approaches are not able to resolve lineage relatedness in *Corbicula*, we investigated how individuals and lineages cluster using an allele sharing-based approach to species delimitation that can delineate species regardless of their monophyly or lack thereof (haplowebs, Flot *et al*., 2010). This is the first time this method is applied to the genus *Corbicula*, providing novel insights into the origin of their invasive alleles. The specimens used in this study were obtained from the worldwide distribution of *Corbicula*, focusing our sampling efforts on regions where genetically diverse *Corbicula* species co-occur, including sexual and/or endemic species (*C. sandai* from Lake Biwa, estuarine *C. japonica* in Japan, and *C. tobae* from Indonesia). Populations where sexual and androgenetic *Corbicula* individuals are found together (such as in Lake Biwa and South-East Asia) are a critical source of information since sympatric populations that include sexuals are putative hotspots for the origin of new asexual lineages (Simon *et al*., 2003).

The main aims of this study are to identify i) the origin of alleles found in invasive *Corbicula* lineages and ii) the genetic mixing that have occurred between distinct lineages, both from the native and invasive range. Owing to the reconstruction method of allele sharing, we predict that one genetic cluster of shared alleles, including a native population, would be observed if all invasive lineages had the same biogeographic origin, while distinct clusters including different invasive lineages would be detected in the case of multiple origins. Given the previously reported genetic relationship of *C. sandai* with androgenetic lineages, the exact nature of shared alleles between this sexual species and each invasive lineage should be identified. In addition, the occurrence of genetic captures between distinct *Corbicula* lineages (*i*.*e*. cytonuclear mismatches and hybridization events) blurs the delimitation of genetic clusters within *Corbicula* genus, unless it is rare. With the allele-sharing method, we will identify the magnitude of genetic exchange. Moreover, if introgressions are frequent but regionally restricted, a pool of shared alleles should be retrieved per geographic region. If this genetic mixing occurs worldwide, we expect to detect one large genetic cluster of shared alleles encompassing most sampled populations.

## Methods

### Specimen collection and reproductive mode characterization

A total of 359 *Corbicula* individuals from 44 distinct localities were considered in this study, collected across the geographic range of *Corbicula*, including both native and invasive areas (see Table S2). Mollusks were collected using a scoop net and preserved in 96% ethanol. The study was restricted to experimentation on bivalve mollusks, thus it did not require any ethical approval.

For samples collected in *Corbicula*’s native range, the species identity was initially based on the mitochondrial haplotype or on published species taxonomic identification in the same region where the collection took place. Because taxonomic identification of native androgenetic *Corbicula* is often uncertain, we refer to some taxa as *C. sp*. (putative species name) (Table S2-S3). The reproductive mode was inferred from the sperm morphology when possible, from the literature available for the sampled population (lineage assignment and/or reproductive morphology) or, if these characteristics were indeterminate, classified as “undetermined”.

Invasive *Corbicula* lineages encompass a wide geographic distribution both in Europe and America, and can be discriminated by eye on the basis of morphological characteristics. We refer to these invasive forms as previously described (A/R, B, Rlc, and C/S: Pigneur *et al*., 2014; note form D was not sampled in this study). Clams sampled from one invasive population (“Wa”, Netherlands) could not be identified as one of these forms because it displays an intermediate morphology. The reproductive mode of invasive individuals was determined based on spermatozoa morphology (as described in Pigneur *et al*., 2014) under the assumption that androgenetic individuals produce biflagellate sperm while sexual clams produce uniflagellate sperm (Komaru & Konishi, 1999). Owing to the very little genetic diversity observed within invasive lineages in previous studies (Lee *et al*., 2005; Hedtke *et al*., 2008; Pigneur *et al*., 2011a, 2014), we analyzed less than 10 individuals from each invasive form for further genetic analyses.

Genetic data from previous studies were included only when the raw data (chromatograms) were available, allowing a thorough check of the final sequences used in our genetic study.

### Genetic analyses

#### DNA isolation, marker amplification, and sequencing

We extracted DNA either from the adductor muscles or the foot using the DNeasy Blood and Tissue Kit (QIAGEN GmbH, Hilden, Germany) following the manufacturer’s protocol. We sequenced four different markers: (1) a 651-bp fragment of the mitochondrial cytochrome c oxidase gene (COI); (2) a 413 to 415-bp fragment of the nuclear 28S gene (28S); (3) a 551 to 660-bp fragment containing the third intron of the nuclear α-amylase gene (*amy*), and (4) a 310 to 365-bp fragment of the hypervariable putative intron of the α subunit of the adenosine triphosphate synthase (*ATPS*). PCR reactions for the four markers were carried out in a mix comprising 1-4 µl DNA, 1X Taq buffer (Promega Corp., Madison, Wisconsin, USA), 0.2 mM nucleotides, 0.3-0.5 µM of each primer, and 0.1 unit of *GoTaq G2* polymerase (Promega) in a total volume of 25 µl. The PCR conditions and primers used for each marker are described in Supplemental Information 1. PCR products were purified and sequenced on an automated sequencer at Genoscreen (Lille, France) or Beckman Coulter Genomics (ishop’s tortford, nited Kingdom).

As polyploidy has been recorded in *Corbicula* (Park *et al*., 2000; Qiu *et al*., 2001) (Figure 1), the occurrence of more than two alleles in each individual is possible. Therefore, direct sequencing data were confirmed by cloning, using the NEB^®^ PCR Cloning Kit with the Q5 High-Fidelity Taq Polymerase and following the manufacturer’s protocol. Between two and four individuals were picked in each population, based on direct sequencing results, to confirm as many alleles as possible. Between 20 and 30 clones per individual were selected randomly and purified using the QIAprep Spin Miniprep Kit (QIAGEN GmbH) in order to cover several times each represented allele. Sequences were obtained from purified plasmids using the Genewiz (Leipzig, Germany) Sanger sequencing service. A column was added to Table S2 for each marker and each individual to indicate the use of direct sequencing, cloning, or both.

#### Sequence editing, alignment, and species delimitation methods

All sequences were first assembled and cleaned using Sequencher 4.1.4 (Gene Codes Corp., Ann Arbor, Michigan, USA). Alignments were made using MAFFT version 7 (Katoh *et al*. 2017) with the iterative refinement method E-INS-I and indels treated as a 5^th^ character state in downstream analyses. Because mitochondrial and nuclear markers potentially have different inheritance patterns in *Corbicula* genus, and because the allele sharing-based approach used in this study cannot be applied to mitochondrial genes, two different clusterization methods were used: one based on the mitochondrial marker using the K/θ method and one based on the nuclear markers using the haploweb and conspecificity matrix approach.

COI sequences were manually edited using BioEdit (Hall 1999) and Sequencher, then tested for cross-contamination using AutoConTAMPR (https://github.com/jnarayan81/autoConTAMPR). No contamination was detected among these tested haplotypes. A median-joining haplotype network (haplonet) was constructed using HaplowebMaker (Spöri *et al*. 2020) based on the COI sequence alignment, including in total 251 COI *Corbicula* sequences. The K/θ method is based on a comparison between intergroup and intragroup sequence diversity (Birky *et al*. 2010; Birky 2013; Birky & Maughan, 2021) as implemented in the online tool KoT (Spöri & Flot, 2021). This method defines a new species when the gap between two clades cannot be caused by genetic drift (based on a threshold) and was shown efficient on asexual or clonal organisms (*e*.*g*.: Schön *et al*., 2012) and using haploid markers (Birky, 2013). The distinct *Corbicula* clusters (*i*.*e*., putative species) defined by this KoT method were color coded on the haplonet constructed for the COI gene and on the haplowebs constructed for each nuclear gene.

For the three nuclear genes (28S, *amy* and *ATPS*), the allele validation based on both direct and cloning sequencing data was done manually in a case-by-case process (Supplemental Information 2) for the 359 individuals with the majority (more than 90%) being sequenced in both directions. Allele sharing based on the three nuclear markers was investigated using the haploweb approach (Doyle, 1995; Flot *et al*., 2010) (Figure S1). The online tool HaplowebMaker (Spöri *et al*., 2020) was used with default parameters to build for each nuclear marker a raw haploweb, which is a median-joining haplotype network (or haplonet, as produced for the mitochondrial, haploid COI marker) on which curves are added connecting nuclear haplotypes or alleles found co-occurring in heterozygous individuals. A group of alleles linked together by heterozygotes creates an exclusive allele pool (AP) (akin to species, Doyle, 1995). The haplowebs were edited manually for visualization purpose. Allele-sharing patterns involving the four invasive forms A/R, C/S, B, and Rlc were manually numbered (using InkScape software, Bah 2011), referring to non-exclusive invasive alleles (invasive alleles that are not specific to one given invasive form) according to the Tables S2 and S3. The allele-sharing information on the three nuclear markers provided by HaplowebMaker was used to build a “conspecificity matrix” (as in Debortoli *et al*., 2016) with the CoMa tool (Spöri *et al*., 2020) with no calculation on heterospecific pairs (option 1: *i*.*e*., no value was given to the absence of sharing). In this matrix, the conspecificity score for each pair of individuals is the number of markers for which these individuals belong to the same allelic pool (ranging from 0 to 3 for the three nuclear markers taken together). After computing the matrix, rows and columns were reordered to maximize the scores along the diagonal using the hierarchical clustering method implemented in the R package ‘‘heatmap3’’ (Zhao *et al*. 2014). Fields For Recombination (FFRs) appear as blocks along the diagonal with high conspecificity scores within blocks, and low scores among them, except in case of introgressions. These introgressions between FFRs were highlighted using a letter code. The final FFR definition was based on the dendrogram (Ward clustering) built by “heatmap3”, and each R was given a different color code. This color code was used in each haploweb based on nuclear markers and on the haplonet constructed for the COI marker to highlight the FFRs defined by the conspecificity matrix using the three nuclear markers. The haplowebs provide details on the allele sharing patterns between FFRs for each marker individually.

Alleles shared between *Corbicula* lineages were also summarized and visualized in a circular plot using the command-line version of Circos (Krzywinski *et al*., 2009). Fifteen distinct groups (namely idiotypes) were manually defined in the Circos plot to highlight the allele-sharing patterns for all four markers (COI, 28S, *amy*, and *ATPS*). More specifically, the four invasive forms (A/R, C/S, B, and Rlc) were considered as four distinct idiotypes, with the invasive Wa population considered separately as it could not be formally identified as one of these four invasive lineages. The brackish water sexual lineage *C. japonica* was also considered a separate group. Other native idiotypes were defined based on the sampled geographic localities (Lake Biwa in Japan including *C. sandai*, other sites in Japan, China, Hawaii, Madagascar, Indonesia, Vietnam, and South Africa). Due to the low number of individuals sampled, we combined the few individuals from Korea, Taiwan, Thailand, and the Philippines together in a last group (or idiotype) named “ outh-East sia”. The occurrence of sexual and androgenetic lineages was represented by mono- and biflagellate spermatozoa, respectively, for each group when known. A color code was used to distinguish the invasive alleles, which fall into two groups: the exclusive invasive alleles, shared between a native lineage and only one invasive form, represented in red (form A/R), blue (form B), green (form Rlc), and orange (form C/S), and the non-exclusive invasive alleles, each found in at least two different invasive forms and some native populations, represented in purple. This same color code has been applied in the Table S2.

All alleles are summarized in Table S2, with unique names within and between the different markers: for each marker, the name of the allele starts with the name of the marker followed by the name of the individual it was retrieved from, with an additional letter (a, b, c, d) if this *Corbicula* individual is heterozygous. If the same allele is found within several individuals, the name used is arbitrarily based on one of those individuals. Alleles shared between distinct idiotypes, based on the Circos analysis, are in bold type, and the same color code is used for invasive alleles. These alleles shared between distinct invasive lineages were named IA (for Invasive Allele), are in purple and are numbered (from 1 up to 3 according to the marker). These numbers are also used in the haplowebs (Figure 2a, b, and c) to highlight these alleles.

**Figure 2:**
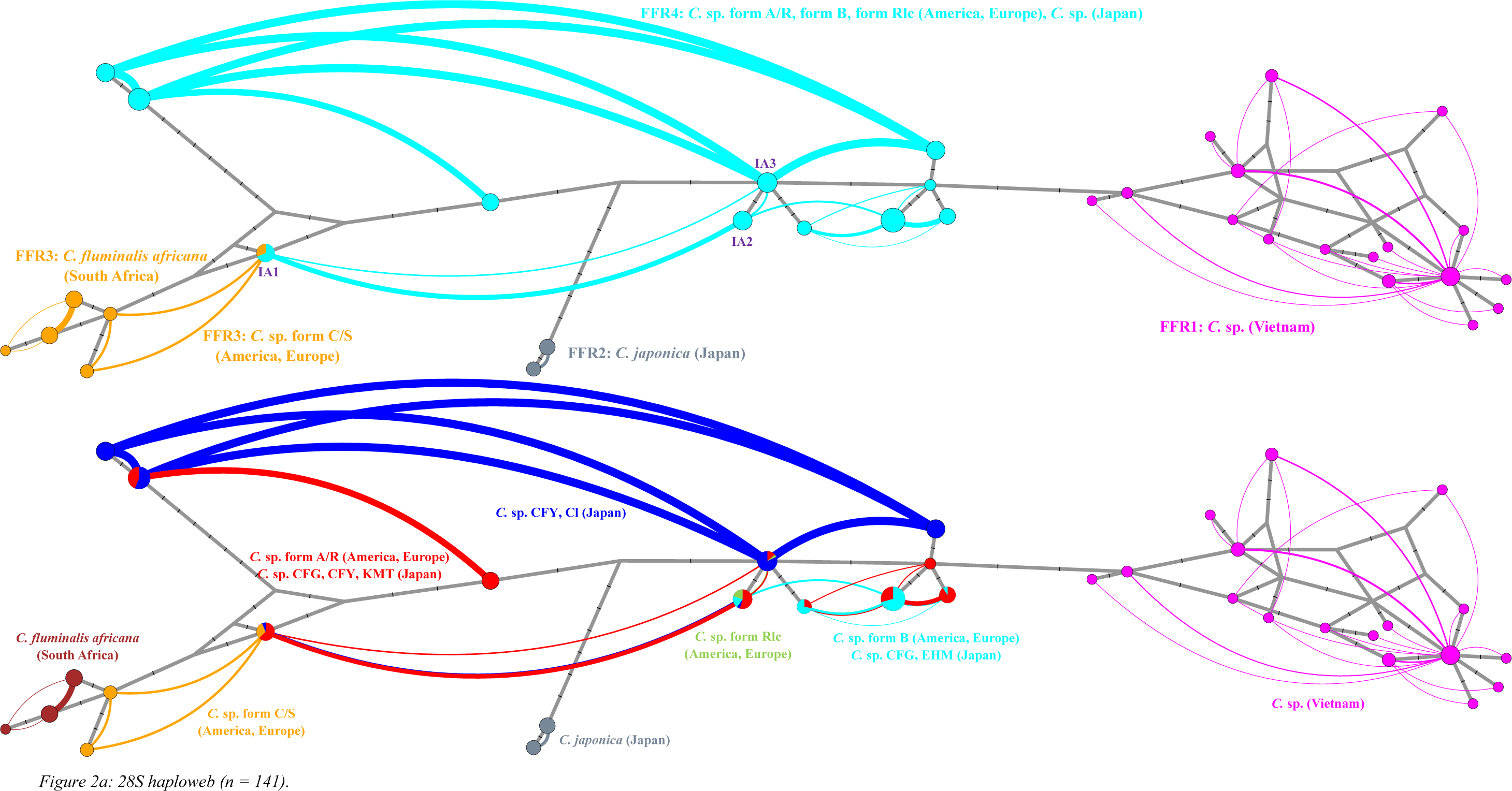
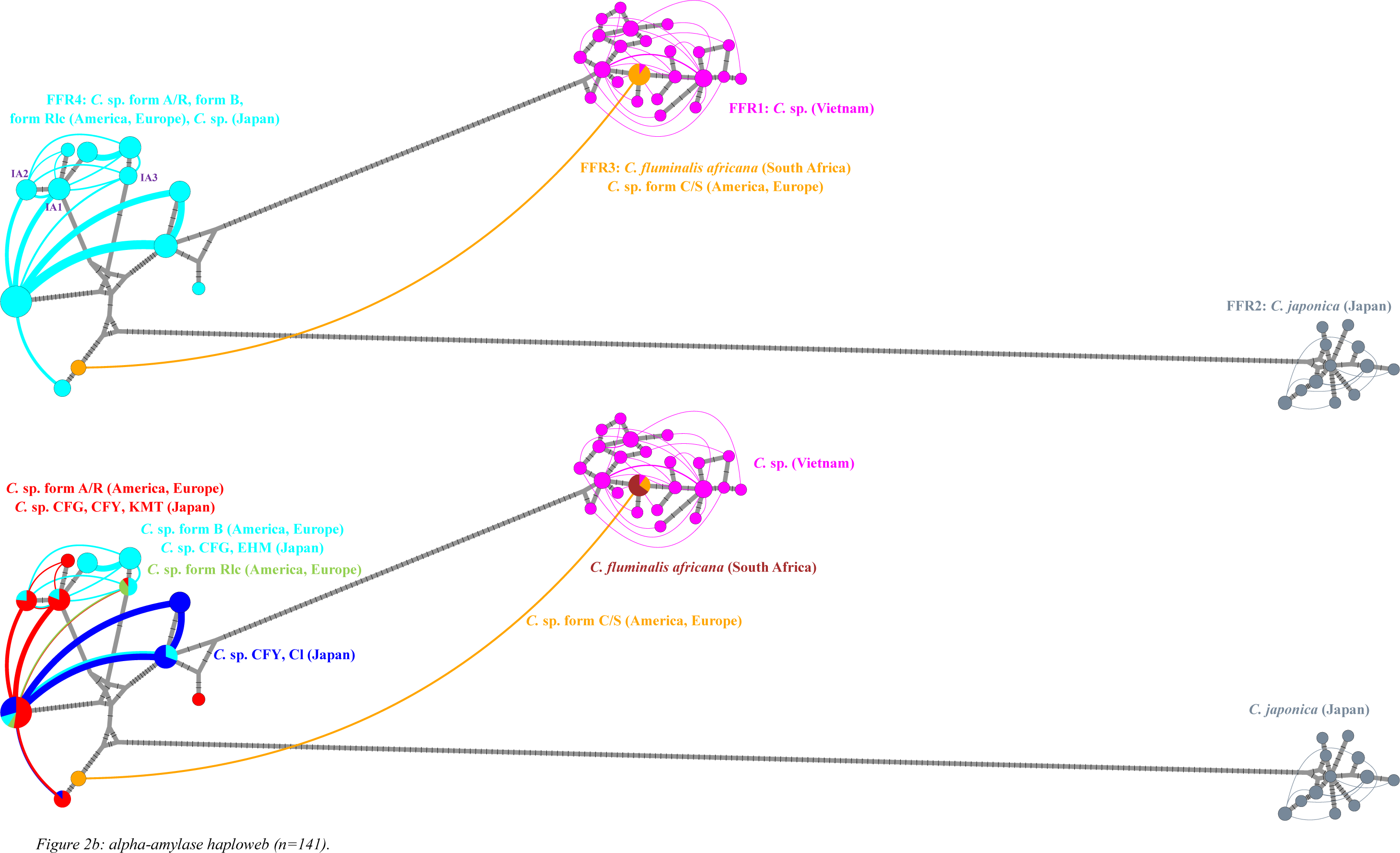
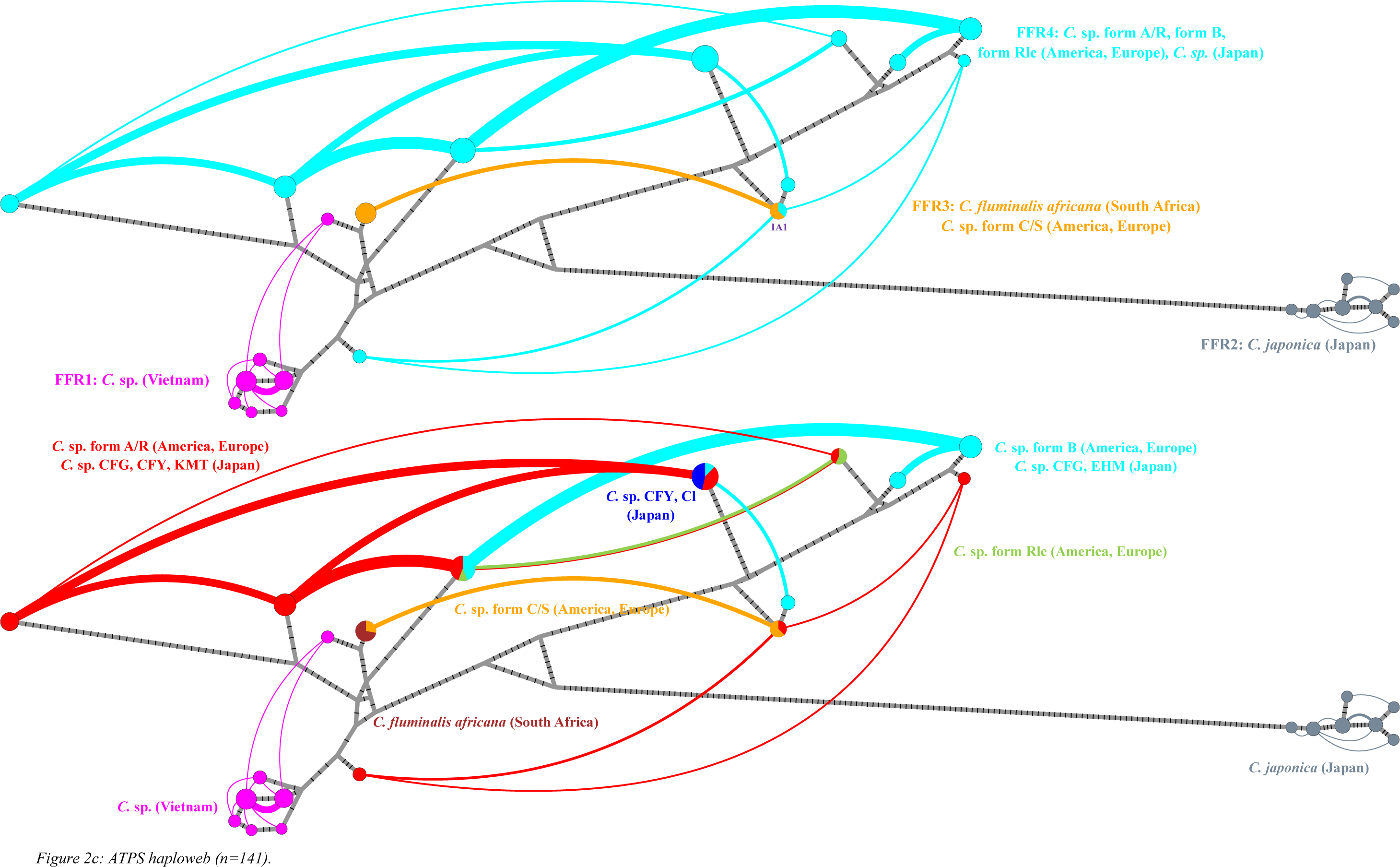
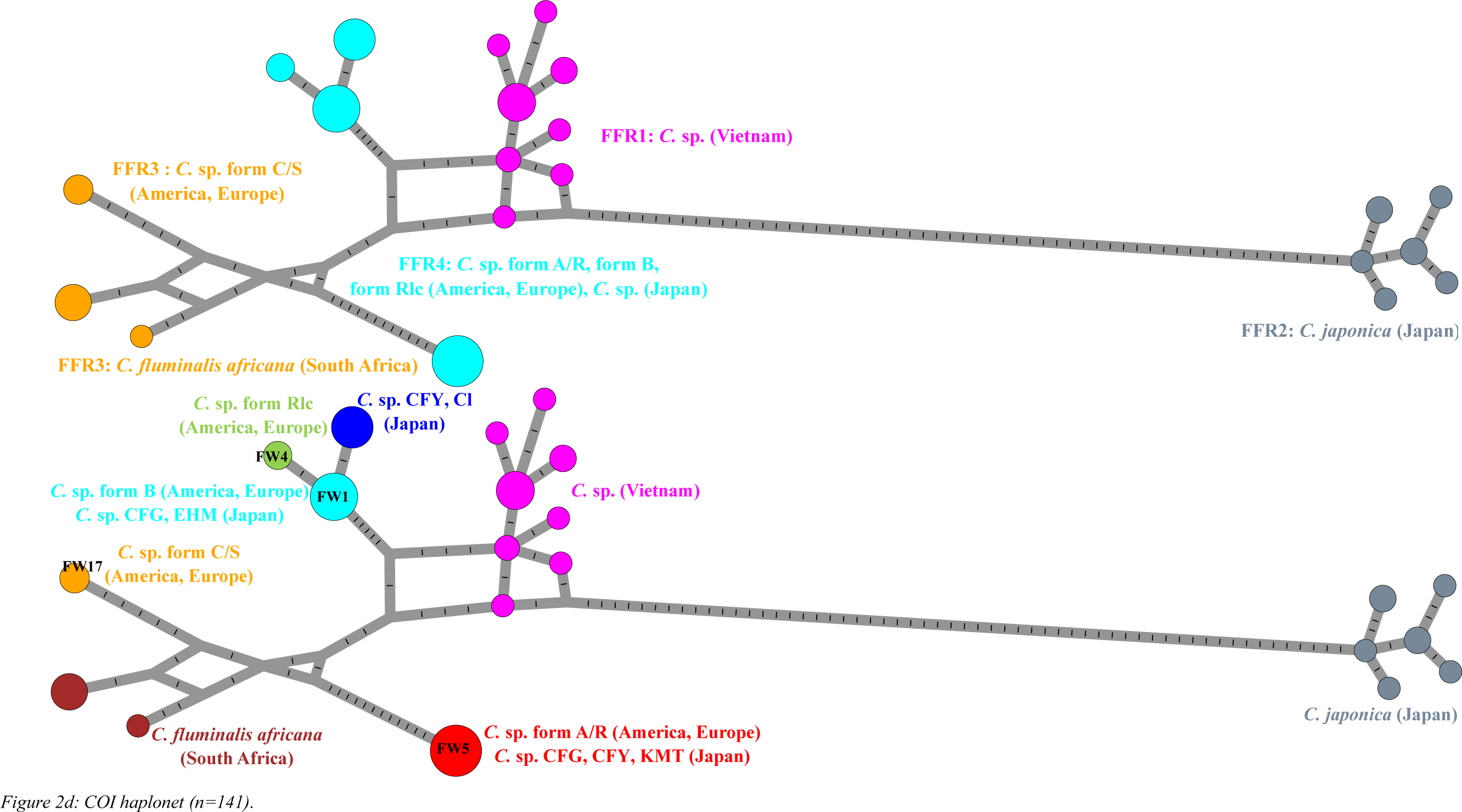
Haplotype webs (haplowebs, see Fig. S1) based on nuclear sequence alignments and haplotype network (haplonet) based on mitochondrial sequence data, all including only the 141 individuals for which all four markers are available. The circle size is proportional to the frequency of the represented alleles/haplotypes. The number of mutation steps inferred by the median-joining algorithm is displayed on the lines connecting the haplotypes. For each marker, [top] alleles on the graph are colored by their nuclear field for recombination (FFR) delimitation (Figure 3) and [bottom] alleles on the identical graph are colored by delimitation based on KoT analysis of the mitochondrial data (Table S5). For haplowebs, the curves connecting each pair of alleles co-occurring in heterozygous individuals are drawn (width proportional to the number of individuals in which the two alleles co-occur), and non-exclusive invasive alleles are numbered from 1 to maximum 3. The name and origin of each Corbicula lineage or allele follows the nomenclature of Table S3. **a)** 28S haploweb, **b)** alpha-amylase haploweb, **c)** ATPS haploweb, **d)** COI haplonet.

All validated sequences of our study were deposited on the NCBI nucleotide database (https://www.ncbi.nlm.nih.gov/)(accession numbers listed in Table S1).

#### Genetic diversity

Nucleotide diversity and haplotype diversity (π and H_d_, respectively) were calculated on the entire dataset (including all alleles even in case of polyploidy) for each marker as the percentage of variable sites between 2 sequences (*i*.*e*., uncorrected pairwise distance) using DNAsp (Librado & Rozas 2009). Gaps, however, were not considered, underestimating the diversity among nuclear alleles.

## Results

Alleles were checked independently using two methods: direct sequencing and allele inferences, and sequencing of cloned amplicons. When the called alleles gave conflicting results between those inferred from direct sequencing and cloned sequences, a manual verification was performed by comparing chromatograms (see Supp. Info. 2). As a result, the majority of alleles obtained in the present work were confirmed by several independent analyses and were identified within multiple individual clams (see Supp. Info. 2). However, some markers were not successfully amplified in an individual, or could not be confirmed with confidence, and were therefore excluded. Consequently, many individuals lack information for one or several markers (see Table S2). For this reason, two distinct datasets are considered in this study: one with all available sequences for all markers, including a total of 359 individuals but with only 251, 266, 237, and 248 individuals for COI, 28S, alpha-amylase, and ATPS markers respectively (Table S2); and a second dataset with only 141 individuals for which all four markers could be amplified and confidently validated (Table S3). These two datasets are referred to as completed and filtered datasets, respectively.

### High heterozygosity and haplotype diversity in *Corbicula*

All markers studied had a relatively high haplotype diversity (H_d_) but a low nucleotide diversity (π), indicating the presence of many low-frequency, genetically similar alleles in *Corbicula* (Table 1). This was also true for the mitochondrial COI marker: 251 sequences revealed 44 distinct haplotypes with 110 polymorphic sites. Moreover, for all three nuclear markers, most *Corbicula* individuals were heterozygous: for 28S, 84% (224/266) were heterozygous, for *amy*, 85% (201/237), while for *ATPS*, only 60% (148/248) were heterozygous (Table S2).

**Table 1:**
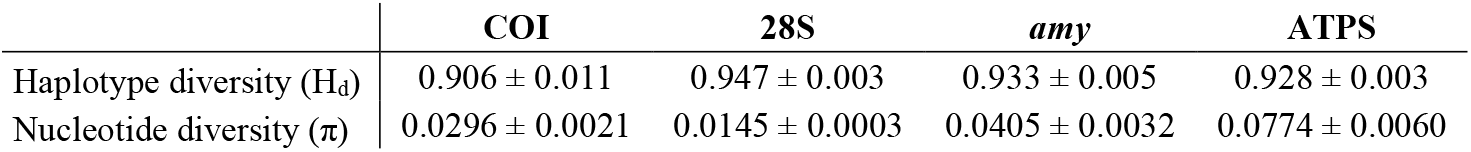
Genetic diversity indices: haplotype diversity (H_d_) and nucleotide diversity (π) for all *Corbicula* samples based on COI, 28S, alpha-amylase, and ATPS.

|  | COI | 28S | *amy* | ATPS |
| --- | --- | --- | --- | --- |
| Haplotype diversity ($H_d$) | 0.906 ± 0.011 | 0.947 ± 0.003 | 0.933 ± 0.005 | 0.928 ± 0.003 |
| Nucleotide diversity ($\pi$) | 0.0296 ± 0.0021 | 0.0145 ± 0.0003 | 0.0405 ± 0.0032 | 0.0774 ± 0.0060 |

For the three nuclear markers studied, *amy* was the most diverse, with 74 alleles retrieved with 147 polymorphic sites in 237 individuals. For *ATPS*, 40 alleles were recovered with 118 polymorphic sites in 248 individuals. 28S was the least diverse nuclear marker of this study, with only 43 polymorphic sites giving 56 alleles in 266 sequenced individuals (complete dataset, Table S2).

### Only four genetic clusters retrieved within the genus *Corbicula* using nuclear markers with two distinct biogeographic origins of the invasive lineages

Haplowebs for the three nuclear markers (using heterozygosity to link alleles, see Fig. S1) and a haplonet for the mitochondrial COI marker were produced for the filtered dataset (Fig. 2) as well as for the complete dataset (Fig. S2). Each resulting graph is presented with two color codes based on the distinct clusterization methods: (1) using the colors from the putative COI species suggested by KoT and (2) using the colors from the consensus species delimitation obtained from the nuclear markers using CoMa.

As the conspecificity matrix analysis gave consistent results between the filtered and complete datasets, only the matrix built from the filtered dataset (n = 141) is presented (Fig. 3) for clarity. Based on the CoMa clustering analysis using only the nuclear markers, a total of four genetic clusters (or FFRs) were delimited in the genus *Corbicula* (including individuals from 17 distinct populations or invasive lineages, see Table S3), suggesting a substantial allele sharing between putative species.

**Figure 3:**
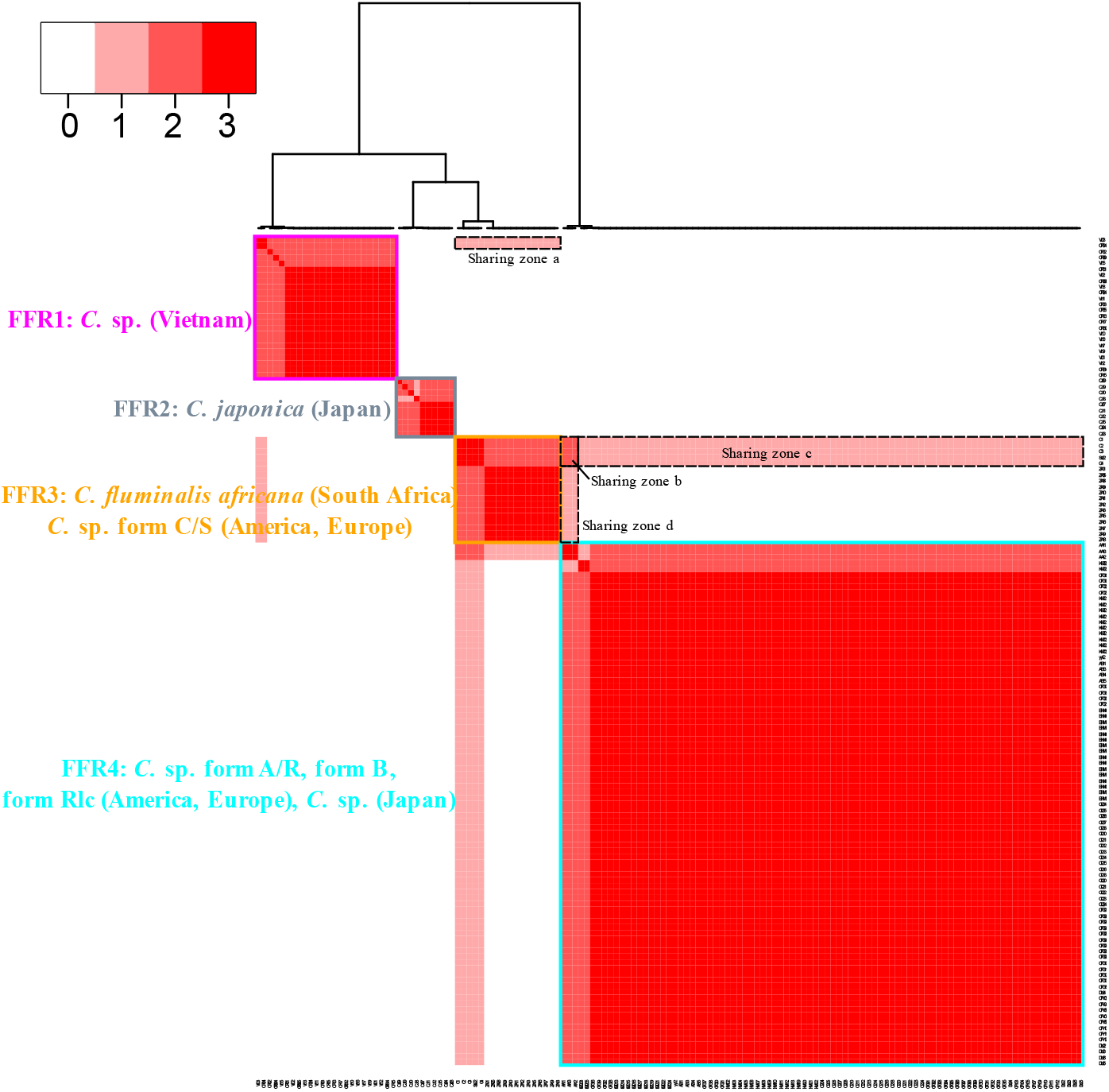
Conspecificity matrix based on alleles shared between individuals across the three nuclear markers (28S, alpha-amylase, ATPS). In this matrix, the axes correspond to the individuals (n = 141) and the red signal corresponds to the number of times individuals were found within the same allelic pool (score from 0 to 3 for all tested markers). Based on the dendrogram (Ward clustering), four FFRs were defined. All individuals are represented using the same color scheme as used for haplowebs and the haplonet (see Figure 2). Signals of allele sharing between these FFRs, corresponding to introgression events, are framed with dotted lines and labelled with letters from a to d. Zone a corresponds to alleles shared between FFR1 and 3, while zones b, c, and d all correspond to sharing between FFR3 and 4. Individual names and geographic origins are detailed in Table S3. Dendrogram and sharing zones are only represented once for clarity.

The first genetic cluster delineated by the analysis, FFR1, contained the androgenetic *Corbicula* lineages from Vietnam, and was well supported by the dendrogram (Figure 3). In the haplowebs, this cluster is clearly separated for all three nuclear markers (Fig. 2 and S2, dark pink clusters).

For all three nuclear markers, the brackish water, sexual species *C. japonica* clustered separately from other clams in both the conspecificity matrix and haplowebs, with no sharing pattern with any other cluster, and is defined as FFR2 based on the dendrogram (Fig.2, 3 and S2, grey clusters).

FFR3 contains *C. fluminalis africana* from South Africa and the invasive *C*. sp. form C/S. These two lineages formed a unique block in the conspecificity matrix, but are independent lineages based on the dendrogram (Figure 3). In the haplowebs, *C. fluminalis africana* and form C/S clustered together for the genes *amy* and *ATPS*, but not for 28S, with the 28S alleles of these two lineages differing only by 1 to 3 SNPs (Fig. 2a and S2a, orange cluster).

FFR4 was the largest FFR in all haploweb analyses and the conspecificity matrix. In the complete dataset, it included the sexual lineage *C. sandai* from Lake Biwa as well as androgenetic individuals across the entire geographic distribution of the genus *Corbicula*, notably China (28S, *ATPS*), Taiwan (*amy*), Korea (*ATPS*), Japan (28S, *amy, ATPS*) including Lake Biwa (*amy, ATPS*), Hawaii (28S, *ATPS*), the Philippines (*ATPS*), Thailand (*ATPS*), Madagascar (28S), and at least three out of the four studied invasive forms from Europe and America (forms A/R, B, and Rlc; Fig. 2 and S2, blue clusters). These species and lineages clustered together in the conspecificity matrix and therefore formed a fourth distinct group, FFR4, based on the dendrogram (Figure 3). In the filtered dataset, it included the three invasive forms and five populations from Japan (Fig. 2, Table S3).

Three of the four FFRs identified in the CoMa analysis were also supported by mitochondrial data. In both the filtered (Fig. 2d) and complete (Fig. S2d) datasets, FFR1, 2, and 3 corresponded to well-clustered sets of COI haplotypes, with a maximum of 4, 6, and 9 SNPs between two haplotypes, respectively, in the complete dataset. However, the FFR4 included highly variable COI haplotypes, with up to 43 SNPs (complete dataset) between two haplotypes from this FFR. The clusterization based on KoT analyses of the COI dataset was represented on both the corresponding mitochondrial haplonet and the nuclear haplowebs (Fig. 2 and S2). For FFR1 and FFR2, the delimitations made by the KoT analyses on the COI dataset (Table S4 and S5) are consistent with the ones from CoMa: all individuals from Vietnamese populations were in a distinct cluster (corresponding to FFR1), and all *C. japonica* individuals were another distinct cluster (corresponding to FFR2). FFR3 was however split into two genetic clusters (orange and brown on Fig. 2) based on COI, because the FW17 haplotype found in *C*. sp. form C/S differs by 7 to 9 SNPs from the sequences found in *C. fluminalis africana*. Finally, FFR4 found by the CoMa analysis is divided into four distinct lineages based on the KoT analysis of COI on the filtered dataset (Table 5) and … lineages based on the complete dataset, with all three invasive lineages (*C*. sp. Form A/R, B and Rlc) having distinct COI sequences and placed in distinct groups. The form A/R clustered with populations from Japan only (CFG, CFY, KMT, LB). In the analysis based on the filtered dataset (Table S5), the form Rlc clustered alone, the form B clustered with native populations from Japan only (EHM, CFG), and the Japanese Cl population clustered alone, while in the complete dataset (Table S4), forms B, Rlc and Japanese populations (EHM, CFG, Cl, LB) clustered together. Additional clusters delimited in Table S5 were the populations from Madagascar, China, Indonesia, with *C. sandai* clustering alone as an independent lineage.

In the end, we observed more genetic clusters within *Corbicula* defined by the COI marker than by the nuclear markers. The KoT analysis using the COI marker defined 11 and 8 putative species based on the complete (n=259, Table S4) and filtered (n=141, Table S5) datasets, respectively. Due to introgression events during androgenetic reproduction between different lineages, distinct COI haplotypes can be retrieved within a nuclear pool of shared alleles. The nuclear markers only delimited 4 genetic clusters in the present study, two of them including only geographically restricted lineages, the two others encompassing lineages from the native and invasive range that were previously described as distinct species. This suggests: i) at least two biogeographic origins of invasive lineages, and ii) a substantial genetic mixing between distinct *Corbicula* lineages.

### Introgressions occurred between three out of the four FFRs of *Corbicula*

Alleles shared between FFRs (as defined by the conspecificity matrix) can be identified and visualized using the haploweb of each nuclear marker. The sexual *C. japonica*, FFR2, does not share alleles with other clusters (Figures 2, 3 and 4). However, alleles are shared between the three other FFRs.

**Figure 4:**
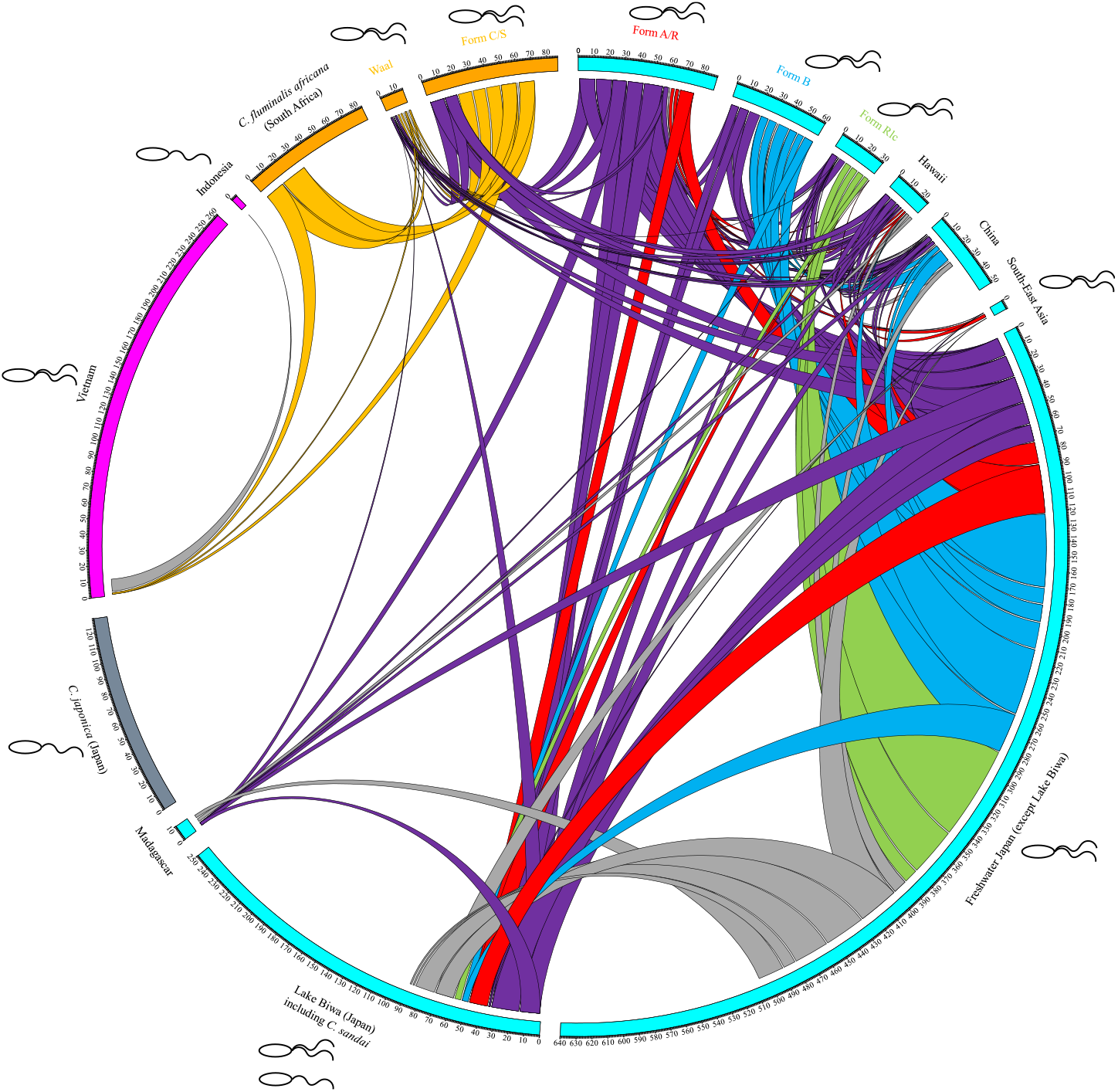
Circos plot representing allele sharing among *Corbicula* (n = 359). Groups (namely idiotypes) are represented by circle arcs with a size determined by the total number of alleles found in each group. Their colors refer to the FFR delimitation from the haplowebs (Fig. 2 and S2) and conspecificity matrix (Fig. 3) analyses. Linking lines between these groups represent shared alleles, with the thickness proportional to the number of occurrences of this allele in each population. The line color refers to alleles found in invasive lineages (Table S2), with exclusive alleles from *C.* sp. form A/R, B, C/S, and Rlc in red, blue, orange, and green, respectively, non-exclusive alleles in purple, and alleles restricted to the native range in grey. When it is known, the occurrence of sexual and androgenetic lineages is represented by mono- and biflagellate spermatozoa, respectively, for each group. For Hawaii, China, Madagascar, and South Africa idiotypes, the reproductive mode is unknown

First, shared alleles are observed between FFR1 and FFR3 for the *amy* marker (Fig. 2b, S2b and Fig. 4), because two Vietnamese individuals (FFR1: Vt03 and CR04) shared the amy-C1b allele with the invasive form C/S and with all individuals from South Africa (FFR3) (Table S2, S3). In the conspecificity matrix, this sharing pattern of one marker was observed and is indicated as zone *a* (Fig. 3).

Second, several sharing patterns were observed between FFR3 and FFR4. The invasive forms A/R and C/S shared an allele for two distinct markers, 28S (Fig. 2a and S2a) and ATPS (Fig. 2b and S2b). In the conspecificity matrix, a sharing pattern for the two markers was therefore highlighted (Fig. 3, zone *b*). For the 28S marker, the shared allele between forms A/R and C/S is 28S-IA1 (Table S2, S3). As form A/R shared one or more of his 28S alleles (28S-IA1, 28S-IA2 and 28S-IA3) with all other populations from FFR4 (Table S3), a sharing pattern of one marker was observed between form C/S from FFR3 and all individuals from FFR4, with allele 28S-IA1 being the link (Fig. 3, zone *c*). For the ATPS marker, the shared allele between forms A/R and C/S is ATPS-IA1. Since form C/S shared its second ATPS allele (ATPS-C1a) with all sequenced individuals of *C. fluminalis africana* from FFR3, a sharing pattern of one marker was observed between form A/R from FFR4 and all *C. fluminalis africana* individuals from FFR3 (Fig. 3, zone *d*).

These sharing patterns between FFRs can also be visualized using a Circos plot (Fig. 4), where the color code of the four FFRs have been applied to the circle arcs representing the groups (idiotypes). Notably, only a very small fraction (2 out of 60) of the individuals from Vietnam (FFR1) were implied in the sharing with FFR3 (including form C/S and *C. fluminalis africana*), which represented 2 out of the 262 alleles retrieved from this location (total number, Fig. 4). In FFR3, all individuals for which the *amy* marker was successfully sequenced had this shared allele with FFR1. The possible introgressions between FFR3 and FFR4 can also be characterized based on the Circos plot, implying only non-exclusive invasive alleles (in purple, Fig. 4) which are never directly found in *C. fluminalis africana*.

### Substantial allele sharing occurred between distinct *Corbicula* lineages

In addition to the previously described allele sharing with form C/S clustering in a distinct FFR, form A/R also shared alleles, for the three nuclear markers tested, with the two other invasive lineages, forms B and Rlc. In total, the four invasive forms A/R, B, C/S, and Rlc from Europe and America shared alleles with each other and with *Corbicula* individuals from 19 different locations in Asia and Africa (Table S2), including the sexual lineage *C. sandai* (Fig. 4, n = 359). Both exclusive and non-exclusive invasive alleles are shared.

In *Corbicula* form A/R, few variations between tested individuals were found within each marker at the nuclear and mitochondrial level, but 7 alleles (out of 12 for all markers) were shared with other invasive forms (non-exclusive invasive alleles, bold purple in Table S2 and Fig. 4). Three other alleles (exclusive invasive alleles) are shared only with native populations (bold red in Table S2 and Fig. 4) and two other alleles are private (red in Table S2). The invasive *Corbicula* form A/R therefore shared alleles with the three other invasive forms (C/S, B, and Rlc) and with freshwater lineages from all studied countries except Vietnam, Indonesia and South Africa (Fig. 4).

*Corbicula* form B also showed low variability between tested individuals and included only 11 distinct haplotypes (across all four markers). Among them, 3 are non-exclusive invasive alleles shared with form A/R or form Rlc (bold purple in Table S2 and Fig. 4), 7 alleles are shared with native populations only (bold blue in Table S2 and Fig. 4), and 1 is a private allele (blue in Table S2). Form B shared alleles with two invasive forms and all freshwater groups defined in this analysis, except populations from Waal (Netherlands), Madagascar, Vietnam, and South Africa.

A total of 8 haplotypes (for all 4 markers) were retrieved in *Corbicula* form Rlc with almost no variability between tested individuals. Four non-exclusive invasive alleles (bold purple in Table S2 and Fig. 4, red for COI) and 4 exclusive invasive alleles (bold green in Table S2 and Fig. 4) were found, but no private alleles. Invasive form Rlc shared alleles with forms A/R and form B but not form C/S, as well as with all native freshwater groups defined in this analysis except the Vietnamese and South African populations.

Within *Corbicula* form C/S, no variability was detected between tested individuals except for one individual from Argentina and one from the Netherlands (Table S2). A total of 12 haplotypes (for all 4 markers) were recorded in form C/S with 2 non-exclusive invasive alleles (bold purple in Table S2 and Fig. 4), 6 exclusive alleles shared with native populations (bold orange in Table S2 and Fig. 4), and 4 private sequences (orange in Table S2). This invasive form only shared alleles with form A/R and Waal (Netherlands) populations, but not with any other invasive lineage. It also shared alleles with individuals from Hawaii, China, and Japan, but never exclusive alleles, and it is the only invasive form which does not share any haplotypes with several Japanese populations (EHM, KMT, CFG), including Lake Biwa where sexual *C. sandai* is found. Form C/S is also the only invasive lineage sharing alleles with Vietnamese populations and extensively with *C. fluminalis africana* (*i*.*e*., these shared alleles are carried by all individuals from both *C. fluminalis africana* and form C/S).

Finally, within the 2 *Corbicula* individuals collected from the Waal river in the Netherlands – whose invasive form could not be identified by morphology – a total of 9 alleles (for all 3 nuclear markers) were retrieved: 3 of them were non-exclusive invasive alleles (bold purple in Table S2 and Fig. 4), and 6 were exclusive to form C/S (bold orange in Table S2 and Fig. 4). Although this population shared alleles with forms A/R, Rlc, and with native populations, its genotype is most closely related to form C/S: 8 out of its 9 shared alleles were identical to those from form C/S. For this reason, it is represented in orange in Figure 4.

In general, among the seven non-exclusive invasive alleles retrieved for the three markers (3 in 28S, 3 in *amy*, 1 in *ATPS*), one (amy-IA3) was shared between form B and form Rlc while the six others were found in form A/R shared with the three other forms (form C/S with 28S-IA1 and ATPS-IA1; form B with amy-IA1 and amy-IA2; form Rlc with 28S-IA2 and 28S-IA3). Even the FW5 COI haplotype, reported as unique to the form A/R, was found within Db25, whose morphology and nuclear markers correspond to form Rlc. As a result, we found more non-exclusive haplotypes (70/89 if all individuals are considered) than exclusive ones within form A/R (Figure 4; Table S2). The proportion of non-exclusive alleles were lower for the other invasive lineages: 21/61 in form B, 12/31 in form Rlc, and 19/87 in form C/S.

*Corbicula fluminalis africana* only shared exclusive invasive alleles (amy-C1b and ATPS-C1a) with form C/S and the Waal (Netherlands) population. One of them (amy-C1b) was also identified in two individuals (Vt03 and CR04) from Vietnam (Fig. 4, Table S2).

All native freshwater *Corbicula* lineages – except Vietnamese populations and *C. fluminalis africana*, which were included in distinct FFRs – shared alleles with each other (Fig. 4). Interestingly, this allele sharing between native *Corbicula* lineages also includes invasive alleles, notably 28S-IA3 which is responsible for the sharing observed between native populations from distinct countries such as Madagascar, China, and Japan, including *C. sandai* (Table S2). In addition, the populations from southeast Asia shared the amy-IA2 allele with Japan including Lake Biwa, and the ATPS-AA12c allele with Hawaii (Table S2).

## Discussion

### Substantial allele sharing but distinct biogeographic origins of *Corbicula* invasive alleles

Species delimitation in *Corbicula* has been fraught with conflict, in part because shell morphology alone is not a suitable parameter to define species due to its phenotypic plasticity (Glaubrecht *et al*. 2007). Moreover, species delimitation using genetic markers and phylogenetic analyses has similarly been challenged by conflicting signals between nuclear and mitochondrial markers (see Hedtke *et al*. 2008, 2011; Pigneur *et al*. 2011a, 2011b, 2014). *Corbicula* taxonomy and the specific status of most freshwater lineages are therefore still debated and there is no consensus on how to name a particular *Corbicula* species. Even the nomenclature for the invasive androgenetic lineages used here (form A/R, form B, form Rlc, and form C/S; Siripattrawan *et al*. 2000; Lee *et al*. 2005, Marescaux *et al*. 2010) could be challenged (*e*.*g*., form Rlc/B, Peñarrubia *et al*., 2017), and new forms are still described (form D, Tiemann *et al*. 2017; Haponski and Foighil 2019).

The haploweb approach used here delineated groups of shared alleles on a worldwide sampling dataset of *Corbicula* clams and found a very limited number of genetic clusters (or “fields for recombination”, FFRs) within this genus, based on three nuclear markers tested (Fig. 2, 3, S2). We observed a single allelic pool encompassing most *Corbicula* freshwater lineages worldwide (Fig. 4). These results highlight the cross-species substantial genetic mixing and introgression events in the clam genus *Corbicula* that have hampered taxonomic species delimitation.

Phylogenetic trees have suggested separate geographic origins for the invasive lineages, with form A/R originating in Japan, forms B and Rlc from the Asian mainland (China, Korea, Vietnam), and form C/S from a *C. moltkiana*-like ancestor from Indonesia, although the latter also clustered with other *Corbicula* lineages depending on the marker used (Siripattrawan *et al*. 2000, Park & Kim 2003, Lee *et al*. 2005, Hedtke *et al*. 2011, Pigneur *et al*., 2014). The allele sharing pattern observed in this study offers an alternative hypothesis.

On one hand, the invasive *C*. sp. forms A/R, B, and Rlc shared extensively their exclusive alleles (*i*.*e*., alleles found in no other invasive lineage) with native populations, mainly in Japan and China but also in Korea, Thailand, the Philippines, and Madagascar (Fig. 4, Table S2). On the other hand, none of the exclusive alleles of form C/S were retrieved in these native samples. Form C/S shared exclusive alleles with nearly all studied individuals of *C. fluminalis africana* from South Africa, and with a few specimens from Vietnam (Fig. 4, Table S2), while no exclusive alleles of the other invasive forms were found in these populations. Moreover, the androgenetic individuals from the Vietnamese populations clustered separately, with the exception of two individuals that shared the amy-C4b allele with form C/S and *C. fluminalis africana* (Fig. 2, 4, S2). Given the extent of nuclear captures taking place between androgenetic *Corbicula* lineages, as highlighted in this study, it is impossible to disentangle the origin of the peculiar reproductive mode of androgenesis in *Corbicula*. Nevertheless, since the origins of the invasive lineages appear recent in comparison, we still observed a signal of distinct biogeographic origins. The allele sharing results obtained here suggest recent and at least three distinct biogeographic sources of androgenetic lineages within the genus *Corbicula*.

First, a southeastern Asian source for the Vietnamese androgenetic *Corbicula* lineages that clustered separately from the other studied individuals (FFR1, Fig. 2, 3, S2). Their sharing of an *α-amylase* allele with one individual of the Indonesian *C. tobiae* (which has unknown reproductive mode) is intriguing. Given the low number of individuals from Indonesia in the present study, the genetic relationships between Indonesian and Vietnamese clams are still unresolved. *Corbicula* populations from this geographic area need a more thorough study, especially since sexual Indonesian *Corbicula* lineages have been reported (Glaubrecht *et al*. 2006). The sharing of one *amy* allele (amy-C4b) between Vietnamese populations, form C/S, and *C. fluminalis africana* (Fig. 2, S2, Table S2, S3) is probably the consequence of a nuclear capture event during an invasion process, although these populations have not been reported to occur in sympatry.

Second, the invasive form C/S showed a clear sharing pattern with the South African *C. fluminalis africana* individuals (whose reproductive mode could not be identified) (Fig. 3, 4). This would confirm a distinct biogeographic invasion source for *Corbicula* form C/S from Africa, the current native range of the species named *C. fluminalis* (Korniushin 2004). This putative African origin of form C/S was already suggested by Pigneur *et al*. (2014) based on COI and 10 microsatellite markers. An alternative hypothesis would be the invasion of South Africa by form C/S and the subsequent establishment of an invasive population in this region. *C. fluminalis africana* would thus originate from the European form C/S rather than the opposite. *Corbicula* genus already occurred in Africa during Pleistocene (Meijer & Preece, 2000) and was still recorded in this continent approximately 50 years before its European invasion (Pilsbry & Bequaert, 1927). Historical data on *Corbicula* distribution in Africa are thus not in favor with that hypothesis but do not exclude it. The form C/S and/or *C. fluminalis africana* could also have originated from another unsampled population from the native range (“ghost population”). ore attention should be given to this frican population in future studies, with a more thorough sampling in the native range and identification of the reproductive mode of these lineages.

Third, extensive allele sharing was observed between the Lake Biwa population in Japan, including the sexual *C. sandai*, and most freshwater *Corbicula* individuals studied here, including three out of the four invasive androgenetic lineages (forms A/R, B, and Rlc) (Fig. 3, 4). The alleles found in these three invasive forms are found in numerous native locations, including several Asian populations, Madagascar, and Hawaii, although the majority of them are found in Japan, including Lake Biwa. Because these alleles are identical across a wide geographic distribution of *Corbicula*, the shared history appears to be recent. Interestingly, within this cluster (FFR4), the population from Lake Biwa is particularly genetically diverse, with numerous private alleles, but also encompassing most alleles found in invasive lineages A/R, B, and Rlc (Table S2, S3). Our analysis therefore points to Japan as a hotspot and distinct biogeographic source of most invasive androgenetic *Corbicula* lineages, particularly Lake Biwa, where sexual *C. sandai* occurs (Houki *et al*., 2011). Repeated origins of asexual clones from a sexual relative have been observed in other taxa where sexual and asexual individuals co-occur (Simon *et al*., 2003; Bode *et al*., 2010; Neiman *et al*., 2010). Moreover, the studies of Komaru *et al*. (2012, 2013) found that androgenetic hermaphroditic individuals of *C. leana* shared 28S rDNA alleles with sexual *C. sandai*, both co-occurring in Lake Biwa. Nevertheless, we cannot argue that *C. sandai* itself is the direct ancestor of the studied androgenetic lineages as a number of morphological, cytological, and biological features (*e*.*g*. biflagellate *vs*. uniflagellate sperm, dioecy *vs*. hermaphroditism, non-brooding *vs*. brooding) distinguish it from androgenetic clams (Glaubrecht *et al*. 2006; Obata *et al*. 2006). More research in Lake Biwa populations, using genome-wide markers, is required to i) confirm the recent biogeographic origin of invasive lineages suggested by the present work and ii) understand the older origin of androgenetic reproductive mode in *Corbicula* genus.

### Genetic mixing between invasive *Corbicula* lineages

While the invasive form C/S found in Europe and South America may have a distinct biogeographic origin in Africa, the sharing of alleles between form C/S and form A/R (Fig. 4, Table S2) is probably the result of secondary nuclear captures when found in sympatry. Indeed, form C/S shared two alleles (28S-IA1 and ATPS-IA1) with the invasive form A/R. This suggests potential post-introductory nuclear genetic exchanges between forms A/R and C/S when found in sympatry (*e*.*g*. Lee *et al*., 2005; Hedtke *et al*., 2008; Pigneur *et al*., 2011, 2014).

Individuals from invasive form B shared alleles with form A/R (amy-IA1 and 2) and form Rlc (amy-IA3), the latter two also sharing two alleles with each other (28S-IA2 and 3). Allele sharing between form A/R and B (Fig. 4, Table S2) probably represents introgression events when found in sympatry, as observed by Hedtke *et al*. (2008). However, form B and form Rlc were almost never reported to occur in sympatry: form B is collected in America and form Rlc in Europe. These forms do, however, share a very similar COI haplotype, with only 1 SNP difference over 650 bp, suggesting a plausible common origin of these forms that later diverged when invading America and Europe. More recently, the FW1 COI haplotype from form B was found in Europe by Penarrubia *et al*. 2017. These authors considered forms and Rlc as a unique form (“Rlc ”), hypothesi ing that alleles shared between these two forms could be due to a close ancestry. More research in the invasive range of *Corbicula* genus, using more sampling sites and genome-wide markers and/or cytology, would provide insights on the mechanism of hybridization between distinct lineages.

Furthermore, a sampling of invasive *Corbicula* lineages in the Iberian Peninsula revealed many heterozygous individuals, with 28S alleles found in form A/R and either form Rlc or form C/S. In some cases, these heterozygotes were associated with the COI haplotype of a third lineage (Peñarrubia *et al*. 2017). Recently, in a man-made channel in Northern European Russia, all sampled *Corbicula* individuals belonged to two distinct invasive lineages apparently fixed for a mismatch between the mtDNA haplotype and morphotype. The first lineage belonged to morphotype Rlc associated with the FW5 haplotype of form A/R, while the second lineage had morphotype A/R associated with FW17 COI haplotype of form C/S (Bespalaya *et al*. 2018). Such cytonuclear mismatches, but also hybrids, have been retrieved in several locations in Europe where form A/R occurs in sympatry with another invasive form (C/S or Rlc) (Pigneur *et al*. 2014; Etoundi, pers. obs.). All these results highlight that nuclear genetic exchanges can occur between invasive *Corbicula* lineages when found in sympatry. In the present study, the two *Corbicula* individuals collected from the Waal River in Europe showed an intermediate phenotype between form A/R and form C/S (unpublished data) and contain alleles from both forms, although a majority was from form C/S, suggesting introgressions occurred after these lineages were introduced. A cytonuclear mismatch was also observed in Db25, carrying all nuclear alleles of form Rlc associated with the COI haplotype of form A/R (Fig. 2d, S2d, Table S2, S3).

Previous studies on *Corbicula* that used phylogenetics to discriminate species observed cytonuclear discordances that confounded species delimitation (*e*.*g*., Hedtke *et al*. 2011). In contrast, the approach used here identified distinct lineages that were consistent across mitochondrial and nuclear markers. CoMa (nuclear) and KoT (mitochondrial) analyses both identified the clusters FF1 and FFR2. The two other FFRs, FFR3 and FFR4, were defined as clusters by the nuclear marker-based analysis only (Figure 2a-c), but as independent lineages according to the COI-based analysis (Figure 2d) even though individuals within FFR3 are located in the same part of the mitochondrial haplotype network, and individuals within FFR4 similarly share nodes with each other. Notably, the different invasive lineages (forms A/R, B and Rlc) were delimited as distinct lineages based on the haploid mitochondrial marker COI but were included within the same FFR because of massive sharing of nuclear alleles between invasive forms and especially with native lineages. Similarly, form C/S and *C. fluminalis africana* are considered distinct lineages based on COI but are included within the same FFR. The debate that remains now is how to delineate species within a genus where frequent introgression occurs between distinct lineages. We strongly recommend the continued use of both mitochondrial and nuclear markers in *Corbicula* when studying genetic relationships and when defining taxa.

### Introgressions between invasive and native *Corbicula* lineages

The genetic diversity within the four studied androgenetic *Corbicula* lineages in the invaded range in Europe and America is very low (reviewed in Pigneur *et al*., 2012), while androgenetic *Corbicula* lineages found in the native range, particularly Asia, are both genetically and morphologically diverse (*e*.*g*. Glaubrecht *et al*., 2003; Hedtke *et al*., 2011; Pigneur *et al*., 2014; Table S2), possibly due to multiple origins from sexual species or multiple introgression events when occurring in sympatry. Interestingly, the most widespread alleles (*e*.*g*.: 28S-IA3, 28S-AB11a, amy-Db22a, ATPS-AB11a) shared between lineages and populations from FFR4 belong to the invasive forms A/R, B, and Rlc and are also shared with *Corbicula* from Japan, China, Madagascar, and Hawaii. Two hypotheses could explain this pattern: (i) these widespread alleles represent shared ancestral alleles present in the native range before the invasion of other Asian countries, America, and Europe; or (ii) these alleles are signatures of nuclear genetic exchanges in contact zones, or both.

A surprisingly higher rate of homozygosity was observed at the *ATPS* locus compared to the 28S and *amy* markers, even in populations that are genetically diverse, such as Lake Biwa. Moreover, we were not able to amplify sequences from the *C. sandai* population and a few others (Table S2), suggesting that there could be mutations in the primer binding sites that prevented amplification (*i*.*e*., null alleles), which would result in an apparent homozygosity. However, at the other two loci (*amy* and 28S), both invasive and native androgenetic *Corbicula* individuals are mostly heterozygotes (also confirmed by Hedtke et al., 2011; Pigneur *et al*., 2014; Peñarrubia *et al*. 2017). One explanation for this observation might be the “ eselson effect”, in which allelic diversity evolves over time in the absence of recombination (Judson & Normark 1996; Birky 1996). However, this divergence would probably impact all alleles in a lineage and would therefore degrade the signal of shared alleles among lineages (Hedtke et al. 2011); we observe instead that identical sequences are shared across sexual and androgenetic lineages. A more plausible explanation for this high heterozygosity is the occurrence of nuclear captures between different lineages. Polyploidy is found across androgenetic *Corbicula* populations (Okamoto & Arimoto 1986, Qiu *et al*. 2001, Ishibashi *et al*. 2003, Komaru *et al*. 2006; Hedtke *et al*. 2008), and up to four different alleles per individual were observed in some populations (Table S2). Egg parasitism by one androgenetic *Corbicula* lineage of a distinct lineage could be accompanied not only by the capture of maternal mitochondrial DNA but also of nuclear DNA, increasing the ploidy and the genetic diversity of hybrids (Figure 1) (Hedtke *et al*. 2011, Peñarrubia *et al*. 2017, Bespalaya *et al*. 2018). We hypothesize that the independent androgenetic *Corbicula* lineages confirmed here originated after hybridization from partial nuclear capture following egg parasitism. These nuclear genetic capture events between lineages, which occurs because these asexual organisms have retained sexual features such as fertilization (Figure 1), might facilitate the evolution and maintenance of genetic diversity through enhanced heterozygosity, with alleles from different genetic lineages being brought together within one individual. This hybridization can occur where distinct androgenetic species co-occur, but also in populations where sexuals and asexuals are found in sympatry (such as Lake Biwa in Japan).

The consequences of androgenetic egg parasitism for chromosome organization, and thus the extent of allele capture, are still unclear in *Corbicula*. Nuclear captures between individual clams have been observed to cause the retention of a complete haploid maternal genome, *i*.*e*. one copy of each chromosome from the mother combined with the diploid or triploid paternal genome (Figure 1b; Komaru et al. 2006). However, in the present study, results from three nuclear loci were not completely consistent with each other, as they led to the description of different allelic pools (Figures 2a, b, and c). This might be explained by non-uniform nuclear capture, with only one or some chromosomes retained from the mother and others discarded during meiosis. Such non-uniform nuclear captures would result in aneuploidy (imbalanced chromosome numbers), but research on *Corbicula* karyotypes have only reported balanced chromosome numbers (two, three, or four copies for all chromosomes, see Okamoto & Arimoto 1986; Komaru *et al*., 1997; Park *et al*., 2000; Qiu *et al*., 2001, Skuza *et al*. 2009). Another explanation for the incongruent nuclear allele pools between markers would be the occurrence of crossing-over and recombination during gamete (*i*.*e*., sperm) production. Nuclear captures were detected between distinct lineages (namely hybridization), but probably also occurs within a lineage, in which case it would be harder to detect. The frequency of this process could thus be underestimated in *Corbicula*. Further research exploring chromosome diversity and gamete production is needed to disentangle the mechanism of nuclear captures in *Corbicula*. This could also shed light on other systems, as androgenesis is associated with peculiarities such as hybridization and/or polyploidy and in some systems the occurrence of closely related sexual species, not only within the genus *Corbicula* but also in very distinct organisms (*e*.*g*., stick-insects *Clonopsis* and *Pijnackeria* genera, Milani *et al*. 2016, 2020).

## Data accessibility

Raw chromatograms including original base call and manual corrections (following the procedure described in Supplemental information 2) for the four marker datasets can be accessed on a Zenodo deposit: https://doi.org/10.5281/zenodo.6302862

For more information, please contact the corresponding authors.

## Acknowledgements

We thank B. Hespeels, M. Terwagne, L. Bankers, K. McElroy, J. Sharbrough, M. Neiman, L. Keller and T. Schwander for advice on the analyses and proofreading, as well as N. Yasuda and theM. Yamaguchi for access to Japanese samples, particularly specimens from Lake Biwa. We are also thankful to all the colleagues who provided samples (C. Ayres, Y. Bagatini, L. Beran, D. Eekelers, J. Higuti, C. Ituarte, K. de Koch, P. Laforge, T. Lee, S. Milla, K. Martens, J. Mouthon, N. Perrin, M.-A. Pierrard, S. Schmidlin, R. Sousa, N. Spann, E. Tinti, M. Yamaguchi and the Illinois Natural History Survey). This work was supported by the Belgian National Fund for Scientific Research (Fonds National pour la Recherche Scientifique - FNRS) through a PhD thesis grant (FC91712) to E.E., by a PDR FNRS research grant (CARMA, 14596412) to K.V.D., by UNamur for the teaching assistant grant to M.V. and by an Horizon 2020 research and innovation program of the European Union under the European Research Council (ERC) grant agreement 725998 (RHEA) to KVD financing postdoctoral researcher E.N. and technician M.C.

Version 4 of this preprint has been peer-reviewed and recommended by Peer Community In Evolutionary Biology (https://doi.org/10.24072/pci.evolbiol.100137).

## Conflict of interest disclosure

The authors of this preprint declare that they have no financial conflict of interest with the content of this article.

## Supplementary material

All following supplementary materials can be accessed as appendix to the main manuscript at: https://doi.org/10.1101/590836

**Supplemental information 1:** PCR conditions for all markers used in this study.

**Supplemental information 2:** details on phasing resolution method on triploid individuals.

**Figure S1:** Schematic description of the haploweb method.

**Figure S2:** Haplotype webs (haplowebs, see Fig. S1) based on nuclear sequence alignments and haplotype network (haplonet) based on mitochondrial sequence data. The circle size is proportional to the frequency of the represented alleles/haplotypes. The number of mutation steps inferred by the median-joining algorithm is displayed on the lines connecting the haplotypes. For each marker, [top] alleles on the graph are colored by their nuclear field for recombination (FFR) delimitation (Figure 3) and [bottom] alleles on the identical graph are colored by delimitation based on KoT analysis of the mitochondrial data (Table S5). For haplowebs, the curves connecting each pair of alleles co-occurring in heterozygous individuals are drawn (width proportional to the number of individuals in which the two alleles co-occur), and non-exclusive invasive alleles are numbered from 1 to maximum 3. The name and origin of each *Corbicula* lineage or allele follows the nomenclature of Table S3. **a)** 28S haploweb (n = 266), **b)** alpha-amylase haploweb (n = 237), **c)** ATPS haploweb (n = 248), **d)** COI haplonet (n = 251).

**Table S1:** GenBank accession numbers of all *Corbicula* haplotypes used in this study. The haplotype designations refer to Table S2 for further information.

**Table S2:** List of the 359 individuals collected for the present study and genetic results obtained. The color code corresponds to the four invasive lineages: exclusive alleles retrieved from *C*. sp. form A/R are in red, these from *C*. sp. form B are in blue, these from *C*. sp. form C/S are in orange and these from *C*. sp. form Rlc are in green. Non-exclusive invasive alleles are in purple. Shared allele from one idiotype to another are in bold.

**Table S3:** List of the 141 individuals for which all markers are available and genetic results obtained, using the same representation as Table S2.

**Table S4:** KoT results on COI marker for the complete dataset (n=359). A group number is assigned to each individual as in KoT output (n=251). This clusterisation was inferred for individual with no COI available but belonging to the same populations. In case of ambiguity, the COI was not inferred, these individuals are labelled in black. The 12 corresponding colors were applied on haplowebs (Figure S2).

**Table S5:** KoT results on COI marker for the filtered dataset (n=141). A group number is assigned to each individual as in KoT output. The 8 corresponding colors were applied on haplowebs (Figure 2).

## Notes

https://doi.org/10.5281/zenodo.6302862

### Competing Interest Statement

The authors have declared no competing interest.

## References

Araujo, R., Moreno, D., Ramos, M.A. 1993. The Asiatic clam Corbicula fluminea (Müller, 1974) (Bivalvia: Corbiculidae)in Europe. American Malacological Bulletin. 10(1):39–49. http://hdl.handle.net/10261/32551

Bah, T. 2011. Inkscape. Guide to a vector drawing program. 4th edition. Prentice Hall. https://inkscape.org/

Barton, N.H. 2009 Why sex and recombination? Cold spring Harbor Symposia on Quantitative Biology 74:187–195. https://doi.org/10.1101/sqb.2009.74.030

Bast, J., Parker, D.J., Dumas, Z., Jalvingh, K.M., Tran Van, P., Jaron, K.S., Figuet, E., Brandt, A., Galtier, N., Schwander, T. 2018. Consequences of asexuality in natural populations: Insights from stick insects. Mol. Biol. Evol. 35, 1668–1677. https://doi.org/10.1093/molbev/msy058

Birky, C.W. Jr., Adams, J., Gemmel, M., Perry, J. 2010. Using population genetic theory and DNA sequences for species detection and identification in asexual organisms. PLoS ONE 5(5): e10609. https://doi.org/10.1371/journal.pone.0010609

Birky, C.W.Jr. 2013. Species detection and identification in sexual organisms using population genetic theory and DNA sequences. PLoS ONE 8(1):e52544. https://doi.org/10.1371/journal.pone.0052544

Birky, C.W. Jr., Adams Maughan, H. 2021. Evolutionary genetic species detected in Prokaryotes by applying the K/θ ratio to N sequences. bioRchiv. doi: https://doi.org/0.0/2020.04.27.062828. https://doi.org/10.1101/2020.04.27.062828

Bespalaya, Y.V., Bolotov, I.N., Aksenova, O.V., Kondakov, A.V., Gofarov, M.Y., Laenko, T.M., Sokolova, S.E., Shevchenko, A.R., Travina, O.V. 2018. Aliens are moving to the Arctic frontiers: an integrative approach reveals selective expansion of androgenic hybrid Corbicula lineages towards the North of Russia. Biol. Invasions. 20:2227–2243. https://doi.org/10.1007/s10530-018-1698-z

Bespalaya, Y.V., Aksenova, O.V., Gofarov, M.Y., Kondakov Kropotin, A.V., Kononov, O.D., Bolotov, I.N. 2020. Who inhabits the world’s deepset crater lake? A taxomnomic review of Corbicula (Bivalvia: Cyrenidae) clams from Lake Toba, North Sumatra, Indonesia. J. Zool. Syst. Res. 00:1–11. https://doi.org/10.1111/jzs.12428

Bode, S.N.S., Adolfsson, S., Lamatsch, D.K., Martins, M.J.F., Schmit, O., Vandekerkhove, J., Mezquita, F., Namiotko, T., Rossetti, G., Schön, I., Butlin, R.K., Martens, K. 2010. Exceptional cryptic diversity and multiple origins of parthenogenesis in a freshwater ostracod. Molecular Phylogenetics and Evolution 54:542–552. https://doi.org/10.1016/j.ympev.2009.08.022

Coughlan, N.E., Cuthbert, R.N., Potts, S., Cunningham, E.M., Crane, K., Caffrey, J.M., Lucy, F.E., Davis, E., Dick, J.T.A. 2019. Beds are burning: eradication and control of invasive Asian clam, Corbicula fluminea, with rapid open-flame burn treatments. Management of Biological Invasions 10(3): 486–499. https://doi.org/10.3391/mbi.2019.10.3.06

Crespo, D., Dolbeth, M., Leston, S., Sousa, R., Pardal, M.A. 2015. Distribution of Corbicula fluminea (Müller, 1774) in the invaded range: a geographic approach with notes on species traits variability. Biol. Invasions 17:2087–2101. https://doi.org/10.1007/s10530-015-0862-y

Debortoli, N., Li, X., Eyres, I., Fontaneto, D., Hespeels, B., Tang, C. Q., Flot, J.-F., & Van Doninck, K. 2016. Genetic exchange among bdelloid rotifers is more likely due to horizontal gene transfer than to meiotic sex. Current Biology 26(6), 723–732. https://doi.org/10.1016/j.cub.2016.01.031

Dimijian, G.G. 2005. Evolution of Sexuality: Biology and Behavior. Baylor University Medical Center Proceedings 18:3, 244-258. https://doi.org/10.1080/08998280.2005.11928075

Dmitriev, D.A., Rakitov, R.A. (2008) Decoding of superimposed traces produced by direct sequencing of heterozygous indels. PLoS Comput. Biol. 4(7):e1000113. https://doi.org/10.1371/journal.pcbi.1000113

Doyle, J.J. 1995. The irrelevance of allele tree topologies for species delimitation, and a non-topological alternative. Systematic Botany 20:574–588. https://doi.org/10.2307/2419811

Flot, J.F., Tillier, A., Samadi, S. & Tillier, S. 2006. Phase determination from direct sequencing of length-variable DNA regions. Molecular Ecology Resources 6:627–630. https://doi.org/10.1111/j.1471-8286.2006.01355.x

Flot, J.F. 2007. Champuru 1.0: a computer software for unraveling mixtures of two DNA sequences of unequal lengths. Molecular Ecology Resources 7:974–977. https://doi.org/10.1111/j.1471-8286.2007.01857.x

Flot J.-F., Magalon, H., Cruaud, C., Couloux, A., Tillier, S. 2008. Patterns of genetic structure among Hawaiian corals of the genus Pocillopora yield clusters of individuals that are compatible with morphology. Comptes Rendus Biologies 331:239–247. https://doi.org/10.1016/j.crvi.2007.12.003

Flot, J.F. 2010. eq H E: a web tool for interconverting H E input/output files and T sequence alignments. Molecular Ecology Resources 10:162–166. https://doi.org/10.1111/j.1755-0998.2009.02732.x

Flot, J.F., Couloux, A. & Tillier, S. 2010. Haplowebs as a graphical tool for delimiting species: a revival of Doyle’s “field for recombination” approach and its application to the coral genus Pocillopora in Clipperton. BMC Evolutionary Biology 10:372. https://doi.org/10.1186/1471-2148-10-372

Glaubrecht, M., von Rintelen, T. & Korniushin, A.V. 2003. Toward a systematic revision of brooding freshwater Corbiculidae in Southeast Asia (Bilvavia, Veneroida): on shell morphology, anatomy and molecular phylogenetics of endemic taxa from islands in Indonesia. Malacologia 45:1–10.

Glaubrecht, M., Fehér, Z., von Rintelen, T. 2006. Brooding in Corbicula madagascariensis (Bivalvia, Corbiculidae) and the repeated evolution of viviparity in corbiculids. Zoologica Scripta 35:641–654. https://doi.org/10.1111/j.1463-6409.2006.00252.x

Gomes, C., Sousa, R., Mendes, T., Borges, R., Vilares, P., Vasconcelos, V., Guilhermino, L., Antunes, A. 2016 Low genetic diversity and high invasion success of Corbicula fluminea (Bivalvia, Corbiculidae) (Müller, 1774) in Portugal. PLoS ONE 11(7): e0158108. doi:10.1371/journal.pone.0158108. https://doi.org/10.1371/journal.pone.0158108

Gomes, C., Mendes, T., Borges, R., Guarneri, I., Marchi, I., Guilhermino, L., Vasconcelos, V., Riccardi, N., Antunes, A. 2020. The genetic diversity of two invasive sympatric bivalves (Corbicula fluminea and Dreissena polymorpha) from Lakes Garda and Maggiore, Northern Italy. Journal of Great Lakes Research 46: 225–229. https://doi.org/10.1016/j.jglr.2019.11.006

Hall, T.A. 1999. BioEdit: a user-friendly biological sequence alignment editor and analysis program for Windows 95/98/NT. Nucleic Acids Research 41:95–98.

Haponski, A.E. & Foighil, D.Ó. 2019. Phylogenomic analyses confirm a novel invasive North American Corbicula (Bivalvia: Cyrenidae) lineage. PeerJ 7:e7484. https://doi.org/10.7717/peerj.7484

Hartfield, M., Keightley, P.D. 2012. Current hypotheses for the evolution of sex and recombination. Integrative Zoology 7: 192–209. https://doi.org/10.1111/j.1749-4877.2012.00284.x

Hedtke, S.M., Stanger-Hall, K., Baker, R.J. & Hillis, D.M. 2008. All-male asexuality: origin and maintenance of androgenesis in the Asian clam Corbicula. Evolution 62:1119–1136. https://doi.org/10.1111/j.1558-5646.2008.00344.x

Hedtke, S.M. 2009. Origin and maintenance of androgenesis: male asexual reproduction in the clam genus Corbicula. Thesis dissertation, University of Texas at Austin (129 pages).

Hedtke, S.M., Glaubrecht, M. & Hillis, D. 2011. Rare gene capture in predominantly androgenetic species. Proc. Natl. Acad. Sci. U.S.A. 108:9520–8524. https://doi.org/10.1073/pnas.1106742108

Hedtke, S.M. & Hillis, D. 2011. The potential role of androgenesis in cytoplasmic-nuclear phylogenetic discordance. Systematics Biology 60:87–109. https://doi.org/10.1093/sysbio/syq070

Hillis, D.M. & Patton, J.C. 1982. Morphological and electrophoretic evidence for two species of Corbicula (Bivalvia: Corbiculidae) in North America. American Midland Naturalist 108:74–80. https://doi.org/10.2307/2425294

Hotta M. & Komaru A. 2018. The process of first polar body formation in eggs of the androgenetic clam Corbicula fluminea. Journal of Shellfish Research 37: 131–137. https://doi.org/10.2983/035.037.0111

Houki, S., Yamada, M., Honda, T. & Komaru, A. 2011. Origin and possible role of males in hermaphroditic androgenetic Corbicula clams. Zoological Science 28:526–531. https://doi.org/10.2108/zsj.28.526

Ishibashi, R., Komaru, A., Ookubo, K., Kiyomoto, M. 2002. The second meiosis occurs in cytochalasin D-treated eggs of Corbicula leana even though it is not observed in control androgenetic eggs because the maternal chromosomes and centrosomes are extruded at first meiosis. Dev. Biol. 244: 37–43. https://doi.org/10.1006/dbio.2002.0590

Ishibashi, R., Ookubo, K., Aoki, M., Utaki, M., Komaru, A. & Kawamura, K. 2003. Androgenetic reproduction in a freshwater diploid clam Corbicula fluminea (Bivalvia: Corbiculidae). Zool. Sci. 20:727–732. https://doi.org/10.2108/zsj.20.727

Ishibashi, R. & Komaru, A. 2006. Abortive second meiosis detected in cytochalasin-treated eggs in androgenetic diploid Corbicula fluminea. Develop. Growth Differ. 48: 277–282. https://doi.org/10.1111/j.1440-169X.2006.00862.x

Katoh, K., Rozewicki, J., Yamada, K.D. 2017. MAFFT online service: multiple sequence alignment, interactive sequence choice and visualization. Briefings in Bioinformatics, bbx108. https://doi.org/10.1093/bib/bbx108

Komaru, A., Konishi, K., Nakayama I., Kobayashi, T., Sakai, H., & Kawamura, K. 1997. Hermaphroditic freshwater clams in the genus Corbicula produce non-reductional spermatozoa with somatic DNA content. Biology Bulletin 193:320–323. https://doi.org/10.2307/1542934

Komaru, A. & Konishi, K. 1999. Non-reductional spermatozoa in three shell color types of the freshwater clam Corbicula fluminea in Taiwan. Zoological Science 16:105–108. https://doi.org/10.2108/zsj.16.105

Komaru, A., Ookubo, K. & Kiyomoto, M. 2000. All meiotic chromosomes and both centrosomes at spindle pole in the zygotes discarded as two polar bodies in clam Corbicula leana: unusual polar body formation observed by antitubulin immunofluorescence. Developmental Genes and Evolution 210:263–269. https://doi.org/10.1007/s004270050313

Komaru, A., Kumamoto, A., Kato, T., Ishibashi, R., Obata, M. & Nemoto, T. 2006. A hypothesis of ploidy elevation by formation of a female pronucleus in the androgenetic clam Corbicula fluminea in the Tone River estuary, Japan. Zoological Science 23: 529–532. https://doi.org/10.2108/zsj.23.529

Komaru, A., Houki, S., Yamada, M., Miyake, T., Obata, M., Kawamura, K. 2012. 28S rDNA haplotypes of males are distinct from those of androgenetic hermaphrodites in the clam Corbicula leana. Dev. Genes Evol. 222:181–187. https://doi.org/10.1007/s00427-012-0395-7

Komaru, A., Yamada, M., Houki, S. 2013. Relationship Between Two Androgenetic Clam Species, Corbicula leana and Corbicula fluminea, Inferred from Mitochondrial Cytochrome b and Nuclear 28S rRNA Markers. Zoological Science. 30(5):360–365. https://doi.org/10.2108/zsj.30.360

Korniushin, A.V. 2004. A revision of some Asian and African freshwater clams assigned to Corbicula fluminalis (Müller, 1774) (Mollusca: Bivalvia: Corbiculidae), with a review of anatomical characters and reproductive features based on museum collections. Hydrobiologia 529:251–270. https://doi.org/10.1007/s10750-004-9322-x

Kraemer, L.R., Swanson, C., Galloway, M. & Kraemer, R. 1986. Biological basis of behavior in Corbicula fluminea. II. Functional morphology of reproduction and development and review of evidence for self-fertilization. American Malacolical Bulletin Spec. Ed., 2:193–202.

Krzywinski, M.I., Schein, J.E., Birol, I., Connors, J., Gascoyne R, Horsman, D., et al. 2009. Circos: An information aesthetic for comparative genomics. Genome Research 19:1639–1645. https://doi.org/10.1101/gr.092759.109

Lee, T., Siripattrawan, S., Ituarte, C.F. & Ò Foighil, D. 2005. Invasion of the clonal clams: Corbicula lineages in the New World. American Malacological Bulletin. 20:113–122.

Lehtonen, J., Jennions, M.D., Kokko, H. 2012. The many costs of sex. Trends in Ecology and Evolution 27(3): 172–178. https://doi.org/10.1016/j.tree.2011.09.016

Lehtonen, J., Schmidt, D.J., Heubel, K. & Kokko, H. 2013. Evolutionary and ecological implications of sexual parasitism. Trends in Ecology and Evolution 28: 297–306. https://doi.org/10.1016/j.tree.2012.12.006

Lenormand, T., Engelstädter, J., Johnston, S.E., Wijnker, E., Haag, C.R. 2016. Evolutionary mysteries in meiosis. Phil. Trans. R. Soc. B 371: 20160001. http://dx.doi.org/10.1098/rstb.2016.0001. https://doi.org/10.1098/rstb.2016.0001

Librado, P. & Rozas, J. 2009. DnaSP v5: A software for comprehensive analysis of DNA polymorphism data. Bioinformatics 25:1451–1452. https://doi.org/10.1093/bioinformatics/btp187

Liegeois, M., Sartori, M. & Schwander, T. 2020. Extremely widespread parthenogenesis and a trade-off between alternative forms of reproduction in mayflies (Ephemeroptera). J. Hered. https://doi.org/10.1093/jhered/esaa027

López-Soriano, J., Quiñonero-Salgado, S., Cappelletti, C., Faccenda, F., Ciutti, F. 2018. Unraveling the complexity of Corbicula clams invasion in Lake Garda (Italy). Advances in Oceanography and Limnology 9(2): 97–104. https://doi.org/10.4081/aiol.2018.7857

Mantovani, B., Passamonti, M. & Scali, V. 2001. The mitochondrial cytochrome oxidase II gene in Bacillus stick insects: ancestry of hybrids, androgenesis, and phylogenetic relationships. Molecular Phylogenetics and Evolution 19:157–163. https://doi.org/10.1006/mpev.2000.0850

Marescaux, J., Pigneur, L.M. & Van Doninck, K. 2010. New records of Asian clams Corbicula spp. in French Rivers. Aquatic Invasions 5:S35–S39. https://doi.org/10.3391/ai.2010.5.S1.009

McKone, M.J., Halpern, S.L. 2003. The evolution of androgenesis. The American Naturalist 161(4): 641–656. https://doi.org/10.1086/368291

McMahon, R.F. 1982. The occurrence and spread of the introduced Asiatic freshwater clam, Corbicula fluminea (Muller), in North America: 1924-1982. Nautilus. 96:134–141.

Meijer, T. & R. C. Preece, 2000. A review of the occurrence of Corbicula in the Pleistocene of North-West Europe. Geologie en Mijnbouw (Netherlands Journal of Geo-sciences) 79(2/3): 241–255. https://doi.org/10.1017/S0016774600021739

Milani, L., Scali, V., Passamonti, M. 2016. Speciation through androgenesis in the stick insect genus Clonopsis (Insecta Phasmatodea). J. Zoolog. Syst. Evol. Res. doi: 10.1111/jzs.12087. https://doi.org/10.1111/jzs.12087

Milani, L., Scali, V., Punzi, E., Luchetti, A., Ghiselli, F. 2020. The puzzling taxonomic rank of Pijnackeria hispanica, a chimerical hybrid androgen (Insecta, Phasmida). Organisms Diversity & Evolution. 20:285–297. https://doi.org/10.1007/s13127-020-00436-1

Minchin, D. 2014. The distribution of the Asian clam Corbicula fluminea and its potential to spread in Ireland. Management of Biological Invasions 5(2): 165–177. https://doi.org/10.3391/mbi.2014.5.2.10

Mouthon, J. 1981. Sur la présence en France et au Portugal de Corbicula (Bivalvia, Corbiculidae) originaire d’Asie. Basteria. 45:109–116.

Neiman, M., Hehman, G., Miller, J.T., Logsdon J.M., Jr. & Taylor, D.R. 2010. Accelerated mutation accumulation in asexual lineages of a freshwater snail. Molecular Biology and Evolution27:954–963. https://doi.org/10.1093/molbev/msp300

Obata, M., Nishimori, K., Komaru, A. 2006. Change of centrosome attachment site causes androgenesis in the freshwater clam Corbicula fluminea: Comparison with C. sandai. Venus 65(3): 247–257.

Okamoto, A. & Arimoto, B. 1986. Chromosomes of Corbicula japonica, C. sandai and C. (Corbiculina) leana (Bivalvia: Corbiculidae). Venus Jpn J. Malacol. 45:194–202.

Park, G., Yong, T., Im, K., Chung, E. 2000. Karyotypes of three species of Corbicula (Bivalvia: Veneroida) In Korea. Journal of Shellfish Research 19(2): 979–982.

Park, J.K. & Kim, W. 2003. Two Corbicula (Corbiculidae: Bivalvia) mitochondrial lineages are widely distributed in Asian freshwater environment. Molecular Phylogenetics and Evolution 29: 529–539. https://doi.org/10.1016/S1055-7903(03)00138-6

Patrick, C.H., Waters, M.N., Golladay, S.W. 2017. The distribution and ecological role of Corbicula fluminea (Müller, 1774) in a large and shallow reservoir. BioInvasions Records 6(1): 39–48. https://doi.org/10.3391/bir.2017.6.1.07

Peñarrubia, L., Araguas, R.M., Vidal, O., Pla, C., Viñas, J., Sanz, N. 2017. Genetic characterization of the Asian clam species complex (Corbicula) invasion in the Iberian Peninsula. Hydrobiologia. https://doi.org/10.1007/s10750-016-2888-2

Pfenninger, M., Reinhardt, F., Streit, B. 2002. Evidence for cryptic hybridization between different evolutionary lineages of the invasive clam genus Corbicula (Veneroida, Bivalvia). Journal of Evolutionary Biology 15: 818–829. https://doi.org/10.1046/j.1420-9101.2002.00440.x

Pichot, C., El Maâtaoui, M., Raddi, S., Raddi, P. 2001. Surrogate mother for endangered Cupressus. Nature 412(5): 39. https://doi.org/10.1038/35083687

Pigneur, L.-M., Marescaux, J., Roland, K., Etoundi, E., Descy J.-P. & Van Doninck, K. 2011a. Phylogeny and androgenesis in the invasive Corbicula clams (Bivalvia, Corbiculidae) in Western-Europe. BMC Evolutionary Biology. 11:147. https://doi.org/10.1186/1471-2148-11-147

Pigneur, L.-M., Risterucci, A.-M., Dauchot, N., Li, X., Van Doninck, K. 2011b. Development of novel microsatellite markers to identify the different invasive lineages in the Corbicula complex and to assess androgenesis. Molecular Ecology Resources, 11: 573–577. https://doi.org/10.1111/j.1755-0998.2010.02963.x

Pigneur, L.-M., Hedtke, S.M., Etoundi, E. & Van Doninck, K. 2012. Androgenesis: a review through the study of the selfish shellfish Corbicula spp. Heredity 108:581–591. https://doi.org/10.1038/hdy.2012.3

Pigneur, L.-M., Etoundi, E., Aldridge, D.C., Marescaux, J., Yasuda, N. & Van Doninck, K. 2014. Genetic uniformity and long-distance clonal dispersal in the invasive androgenetic Corbicula clams. Molecular Ecology 23:5102–16. https://doi.org/10.1111/mec.12912

Pilsbury, H.A., & Bequaert, J. (1927) The aquatic molluscs of the Belgian Congo, with a geographical and ecological account of Congo malacology. Bull. Amer. Mus. Nat. Hist. 53: 69–602.

Qiu, A., Shi, A., Komaru, A. 2001. Yellow and brown shell color morphs of Corbicula fluminea (Bivalva: Corbiculidae) from Sichuan Province, China, are triploids and tetraploids. Journal of Shellfish Research 20(1): 323–328.

Rey, O., Facon, B., Foucaud, J., Loiseau, A., Estoup, A. 2013. Androgenesis is a maternal trait in the invasive ant Wasmannia auropunctata. Proc R Soc B 280: 20131181. https://doi.org/10.1098/rspb.2013.1181

Reyna, P., Nori, J., Ballesteros, M.L., Hued, A.C., Tatián, M. 2018. Targeting clams: insights into the invasive potential and current and future distribution of Asian clams. Environmental Conservation 1–9. https://doi.org/10.1017/S0376892918000139

Sagata, N. 1996. Meiotic metaphase arrest in animal oocytes: its mechanisms and biological significance. Trends in Cell Biology 6: 22–28. https://doi.org/10.1016/0962-8924(96)81034-8

Sano, N., Houki, S., Kodan, A., Kawamura, K., Yamada, M., Komaru, A. 2020. Genetic confirmation of “egg parasitism” in androgenetic freshwater Corbicula clams by paternity testing using microsatellite DNA markers. Plankton Benthos Res. 15(1): 58–62. https://doi.org/10.3800/pbr.15.58

Simon, J.C., Delmotte, F., Rispe, C. & Crease, T. 2003. Phylogenetic relationships between parthenogens and their sexual relatives: the possible routes to parthenogenesis in animals. Biological Journal of the Linnean Society 1: 151–163. https://doi.org/10.1046/j.1095-8312.2003.00175.x

Schön, I., Pinto, R.L., Halse, S., Smith, A.J., Martens, K., Birky, C.W.Jr. 2012. Cryptic species in putative ancient asexual Darwinulids (Crustacea, Ostracoda). PLoS ONE 7(7): e39844. https://doi.org/10.1371/journal.pone.0039844

Schwander, T., & Crespi B.J. 2009. Multiple direct transitions from sexual reproduction to apomictic parthenogenesis in Timema stick insects. Evolution 63:84–103. https://doi.org/10.1111/j.1558-5646.2008.00524.x

Schwander, T., Oldroyd, B.P. 2016. Androgenesis: where males hijack eggs to clone themselves. Phil. Trans. R. Soc. B 371:20150534. https://doi.org/10.1098/rstb.2015.0534

Sequencher® version 4.1.4 DNA sequence analysis software, Gene Codes Corporation, Ann Arbor, MI USA http://www.genecodes.com

Siripattrawan, S., Park, J.-K. & Ò Foighil, D. 2000. Two lineages of the introduced Asian freshwater clam Corbicula occur in North America. Journal of Molluscan Studies 66: 423–429. https://doi.org/10.1093/mollus/66.3.423

Skuza, L., Labecka, A.M., Domagala, J. 2009. Cytogenetic and morphological characterization of Corbicula fluminalis (O. F. Müller, 1774) (Bivalvia: Veneroida: Corbiculidae): Taxonomic status assessment of a freshwater Clam. Folia biologica (Kraków) 57: 3–4. https://doi.org/10.3409/fb57_3-4.177-185

Spöri, Y., Flot, J.F. 2020. HaplowebMaker and CoMa: Two web tools to delimit species using haplowebs and conspecificity matrices. Methods Ecol. Evol. 00:1–5. https://doi.org/10.1111/2041-210X.13454

Spöri, Y., Stoch, F., Dellicour, S., Birky, C.W.Jr., Flot, J.F. 2021. KoT: an automatic implementation of the K/theta method for species delimitation. Bioarchiv. https://doi.org/10.1101/2021.08.17.454531

Stephens, M., Smith, N. & Donnelly, P. 2001. A new statistical method for haplotype reconstruction from population data. American Journal of Human Genetics 68: 978–989. https://doi.org/10.1086/319501

Tiemann, J.S., Haponski, A.E., Douglass, D.A., Lee, T., Cummings, K.S., Davis, M.A., Foighil, D.Ó. 2017. First record of a putative novel invasive Corbicula lineage discovered in the Illinois River, Illinois, USA. BioInvasions Records. 6(2):159–166. https://doi.org/10.3391/bir.2017.6.2.12

van der Kooi, C.J. & Schwander, T. 2014. On the fate of sexual traits under sexuality. Biol. Rev. 89:805–819. https://doi.org/10.1111/brv.12078

Voroshilova, I.S., Pryanichnikova, E.G., Prokin, A.A., Sabitova, R.Z., Karabanow, D.P., Pavlov, D.D., Kurina, E.M. 2021. Morphological and genetic traits of the first invasive population of the Asiatic clam Corbicula fluminea (O.F. Müller, 1774) naturalized in the Volga river basin. Russian Journal of Biological Invasions. 12(1): 36–43. https://doi.org/10.1134/S2075111721010148

Yamada, M., Ishibashi, R., Toyoda, K., Kawamura, K. & Komaru, A. 2014. Phylogeography of the brackish water clam Corbicula japonica around the Japanese archipelago inferred from mitochondrial COII gene sequences. Zoological Science 31: 168–179. https://doi.org/10.2108/zsj.31.168

Zhao, S., Guo, Y., Sheng, Q., & Shyr, Y. 2014 Advanced heat map and clustering analysis using Heatmap3. BioMed Research International. Article ID 986048, 6p. https://doi.org/10.1155/2014/986048

