## Supplemental information 1 for "Substantial genetic mixing among sexual and androgenetic lineages within the clam genus *Corbicula*"

PCR conditions for the amplification of COI, 28S, *amy* and ATPS markers.

| Marker | Type | PCR cycle conditions | Primers (5'-3') | Fragment length (bp) (indels included) | Fragment length (bp) (indels excluded) |
|---|---|---|---|---|---|
| COI | Mitochondrial | 30 cycles: 45s at 94°C, 45s at 45°C, 45s at 72°C | LCO 1490: GGTAACAAATCATAAAGATATTGG<br>HCO 2198: TAAACTTCAGGGTGACCAAAAAATCA<br>(Folmer *et al.* 1994) | 651 | 651 |
| 28S | Nuclear | 35 cycles: 10s at 94°C, 30s at 53.5°C, 60s at 72°C | D23F: GAGAGTTCAAGAGTACGTG<br>D4RB: TGTTAGACTCCTTGGTCCGTGT<br>(Park & O' Foighil 2000) | 425 | 413-415 |
| *amy* | Nuclear | 35 cycles: 10s at 94°C, 30s at 54°C, 60s at 72°C | AmyND-f: ATGGTGCAACGATGCAAG<br>AmyND-r: TGATAACCACATCTACCAAGATCC<br>(Adapted from Hedtke *et al.* 2011) | 711 | 551-660 |
| ATPS | Nuclear | 35 cycles: 10s at 94°C, 30s at 60°C, 60s at 72°C | ATPSaSH1f: GTGCCCATYGGWAGAGGACAGAGAG<br>ATPSaSH4r: TGATGGTGTCAATAGCAATGG CAG T<br>(Hedtke *et al.* 2011) | 392 | 310-365 |


Folmer O., Black M., Hoeh W., Lutz R., Vrijenhoek R. 1994. DNA primers for amplification of mitochondrial cytochrome c oxidase subunit I from diverse metazoan invertebrates. *Molecular Marine Biology and Biotechnology*, **3**, 294-9

Hedtke S.M., Glaubrecht M., Hillis D. 2011. Rare gene capture in predominantly androgenetic species. *Proceedings of the National Academy of Sciences*, USA, **108**, 9520-8524

Park J.K., O’Foighil D. 2000. Sphaeriid and Corbiculid clams represent separate heterodont bivalve radiations into freshwater environments. *Molecular Phylogenetics and Evolution*, **14**, 75–88
