## Supplemental information 2 for "Substantial genetic mixing among sexual and androgenetic lineages within the clam genus *Corbicula*"

### Supplemental information 2: phasing on polyploid samples

Because polyploidy has been recorded in *Corbicula* (Okamoto & Arimoto 1986, Komaru *et al.*, 1997; Park *et al.*, 2000; Qiu *et al.*, 2001, Ishibashi *et al.*, 2003, Komaru *et al.*, 2006; Hedtke *et al.*, 2008), the occurrence of more than two alleles in a given individual is possible. The classical tools used to phase heterozygous alleles from direct Sanger sequencing of the gene of interest in diploid individuals (CHAMPURU, PHASE and INDELLIGENT, see below) could fail or generate incorrect alleles when more than two alleles are present. Therefore, we used cloning and subsequent sequencing of individual clones to identify alleles in polyploid samples. Here, more information is given on how these two methods were combined to identify three or even four different alleles within the same *Corbicula* individual. Notice the words "*allele*" and "*haplotype*" are used as synonyms in the whole document. Chromatogram illustrations are printed from the Sequencher 4.1.4 (Gene Codes Corp., Ann Arbor, Michigan, USA) user interface. This interface is described in **Figure 1**.

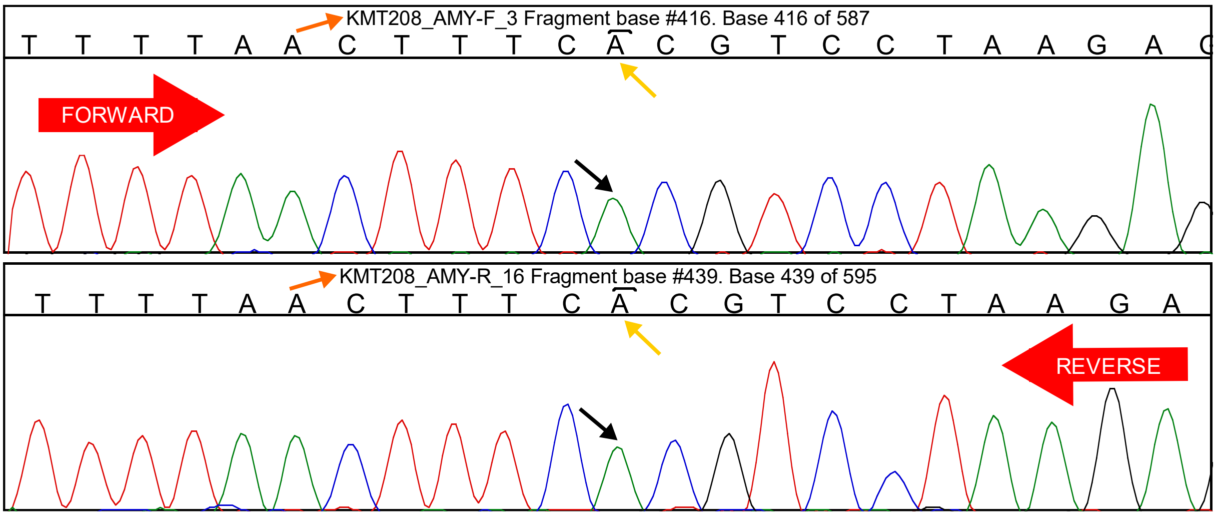


***Figure 1:*** *The Sequencher interface shows chromatograms using two windows: one (top) corresponding to the sequence obtained from the forward primer sequencing, the other (bottom) corresponding to the reverse primer sequencing of the same region (the red arrows show the sequencing orientation of both fragments). All information thus appears twice, such as the header (orange arrow) including the name of the individual, the primer used and the position in the sequence corresponding to the highlighted base (yellow arrows). The peak associated to this base is shown below (black arrow). Each base has its own colour code: A, T, C, and G are in green, red, blue and black, respectively.*

First, information from direct Sanger sequencing data was retrieved using one of these three tools:

- **PHASE** (Stephens *et al.*, 2001), used with SeqPHASE (Flot 2010) for data formatting, allows a phasing (allele reconstruction) of heterozygotes with two alleles of identical length (*i.e.*, without indels). These heterozygous individuals can display double peaks in their direct Sanger sequencing chromatograms, indicative of single nucleotide polymorphisms (SNPs) at that position between the two alleles (**Figure 2**). If only one SNP is observed, the phasing is obvious and can be done "by eye". However, if more than one SNP occurs, an analytical tool must be used to determine which character states are found on the same chromosome to reconstruct haplotypes. PHASE performs this using posterior probabilities based on the expectation that the same haplotypes are found in various combinations. However, it assumes that only two alleles must be reconstructed. If a polyploid has more than two alleles that are the same length, the chromatogram could look identical to a diploid, assuming no more than two character states exist at a given position. Indeed, the occurrence of more than two character states at the same position would return a triple peak in direct Sanger sequencing chromatogram, thus triploidy is obvious is that case.

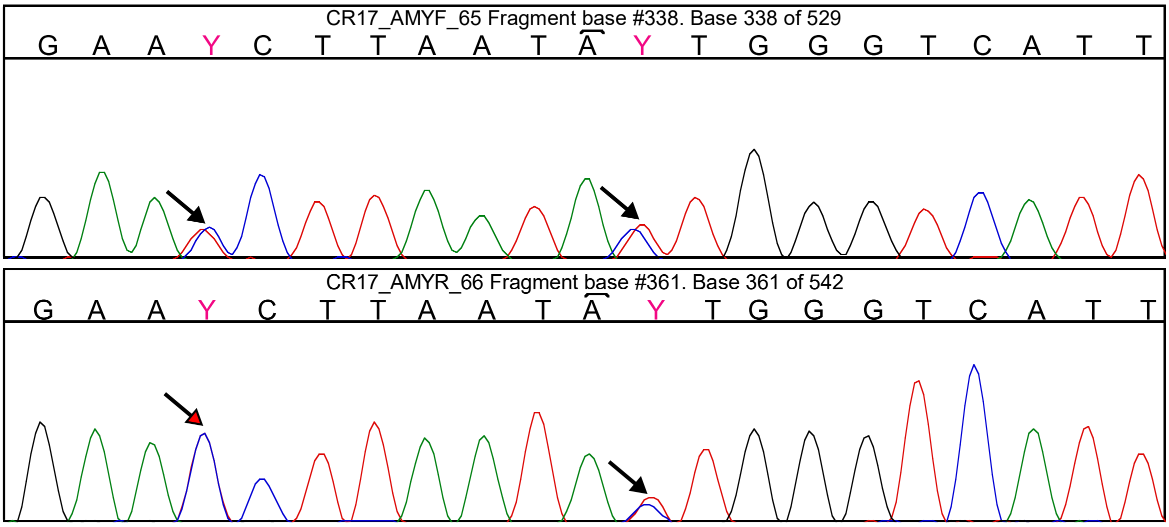


***Figure 2:*** *Heterozygote with two visible double peaks (black arrows). Notice the first double peak in hardly visible on the reverse chromatogram (red-headed arrow), as there is a nearly perfect overlap. Sequencher allows the researcher to hide one base to remove any ambiguity. At these two positions, the base was edited manually (bases in bold pink) to add the ambiguity based on the IUPAC code (here a Y letter stands for both a C and a T).*

- **CHAMPURU** 1.0 (Flot 2007) and 1.1 (https://eeg-ebe.github.io/Champuru/input.html) allows direct phasing of length-variant heterozygotes (Flot *et al.* 2006). The direct Sanger sequencing chromatograms of such individuals display numerous double peaks occurring from a certain point (typically after the indel) until the end of both forward and reverse chromatograms (**Figure 3**). Based on the forward and reverse sequences, in which all double peaks are correctly reported using the IUPAC one-letter code, CHAMPURU can reconstruct two sequences. However, if more than two alleles are represented in the direct sequencing data, this can prevent a correct allele reconstruction or, even worse, lead to the reconstruction of erroneous sequences. CHAMPURU is also unable to deal with triple peaks.

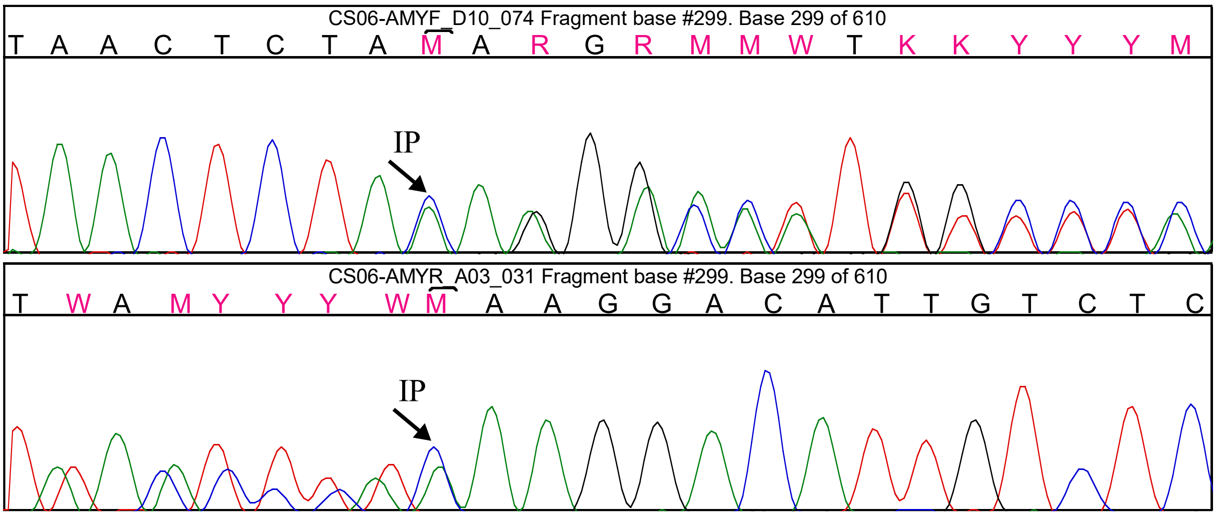


***Figure 3:*** *Length-variant heterozygote. The "inflexion point" – where the double peaks start to appear in both forward and reverse chromatograms in their respective orientations ("IP", black arrows) – shows the indel between the two alleles.*

- **IN*D*ELLIGENT** v. 1.2 (Dmitriev & Rakitov 2008) allows phasing alleles when indels are at different positions but are the same size in each allele. As a result, these alleles are non-length variants but the chromatograms will display numerous double peaks from a certain point to another (**Figure 4**). As these double peaks would be identical in both forward and reverse sequences (which is not the case with a length-variant heterozygous), IN*D*ELLIGENT reconstructs two alleles only based on one sequence, in which the double peaks are reported using the IUPAC one-letter code. This tool will simply add indels on both alleles to explain the double peaks. Once again, it can not deal with the occurrence of more than two alleles.

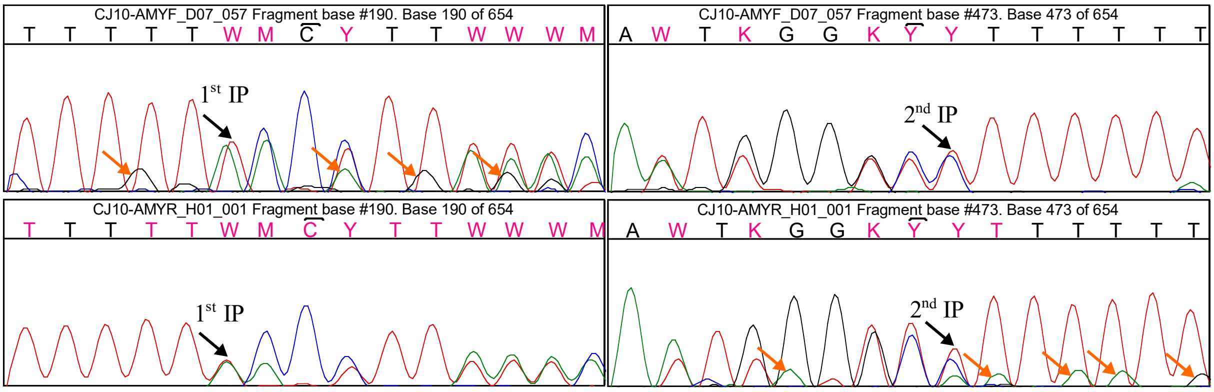


***Figure 4:*** *Heterozygote with compensating indels in both alleles. In this case, there are two inflexion points (IP, black arrows), an indel from the first allele generating double peaks then another indel of equal size from the second allele restoring the frame. Notice some minor peaks are visible here (orange arrows), probably due to a contaminant as this population is strictly sexual. Usually, contaminants are easy to distinguish, as they are only found in one sequence (forward or reverse).*

These tools alone are not enough to confidently build the alleles of a population where triploidy or even tetraploidy occurs. Only two alleles will be reconstructed for each individual, and these tools cannot handle triple peaks, which were frequently observed in androgenetic *Corbicula* populations (not quantified). Therefore, cloning was used to verify the alleles retrieved by direct sequencing phasing methods and to help resolve the missing alleles of polyploid individuals. Only alleles that were consistent with data from both direct sequencing and cloning were validated.

Practically, most of the encountered phasing resolutions fell into one of these cases:

- **CASE 1: homozygosity**. Forward and reverse chromatograms show the same sequence with no double peak, only one allele is validated, no tool nor cloning required (**Figure 1**);
- **CASE 2:** heterozygosity (2 alleles), **no length variant, one SNP**. The forward and reverse chromatograms display only one double peak (same as **Figure 2** but implying only one position), the two alleles can be reconstructed “by eyes”, no tool nor cloning are required. Occurrence of a single triple peak could also be solved by eyes (3 alleles) but this case was never encountered in the present work;
- **CASE 3:** heterozygosity (2 alleles or more), **no length variant, more than one SNP**. The forward and reverse chromatograms display several double peaks without inflexion point (**Figure 2**), which cannot be solved by eye. These cases can be solved using PHASE (Bayesian inference of the phasing based on other sequences from the dataset). Alternatively (*e.g.:* when alleles constructed by PHASE are poorly supported), these alleles can be retrieved and corrected by cloning. This confirmation was considered if at least one of the retrieved alleles was not confirmed in the dataset, as it could result from a mix between two alleles in case of triploidy. Occurrence of one or several triple peak(s) would also be possible (3 alleles) but this case was never encountered in the present work;
- **CASE 4:** heterozygosity (2 alleles), **no length variant, compensating indels**. Both the forward and reverse chromatograms display numerous double peaks with two inflexion points, meaning no double peak is observed at the beginning nor at the end of the sequence (**Figure 4**). This case can be solved using IN*D*ELLIGENT. Alternatively (*e.g.:* when the alleles are not completely

resolved), theses alleles can be retrieved and corrected by cloning. This confirmation was considered if at least one of the retrieved alleles was not confirmed in the direct sequencing dataset.

- **CASE 5:** heterozygosity (2 alleles), **length variant.** The forward and reverse chromatograms contain numerous double peaks from a certain point (marking the location of the indel) until the end of both chromatograms (**Figure 3**). In these cases, all double peaks are annotated using the correct ambiguous letter (IUPAC code), and the two corresponding alleles can be reconstructed by CHAMPURU. Cloning may be used to confirm these retrieved alleles, mainly if the retrieved alleles are not both confirmed alleles in the dataset, as this method alone can generate erroneous alleles in case of triploidy (see CASE 6);
- **CASE 6a:** heterozygosity (3 alleles), **one length variant and two alleles of same size, no triple peak.** This case will usually display chromatograms similar to the previous one (**Figure 3**), but most of the time, CHAMPURU will not be able to explain all the double peaks with only 2 alleles if ambiguity code annotation corresponds to the chromatogram. These cases can be (at least partially) solved by CHAMPURU and PHASE, using a two-step process: first solving the length variant allele using CHAMPURU, which would generate a second allele containing at least one ambiguity, then solving the two remaining alleles by eye (case 2) or with PHASE (case 3). Confirmation of these alleles by cloning was considered in this case. Notice in rare cases (not quantified), CHAMPURU can reconstruct two alleles, but at least one of them would be a chimeric allele (actually resulting from two mixed alleles) and a singleton in the dataset (encountered in 28S marker for CFY06 individual, for example). In that case, the acquisition of cloning data is mandatory to retrieve reliable alleles.
- **CASE 6b:** heterozygosity (3 alleles), **one length variant and two alleles of same size, one or several triple peak(s).** This case is the same as the previous one but is easier to assign to a triploid situation, as at least one triple peak will be observed (**Figure 5**). It can be solved the same way as case 6a with cloning being recommended to confirm.

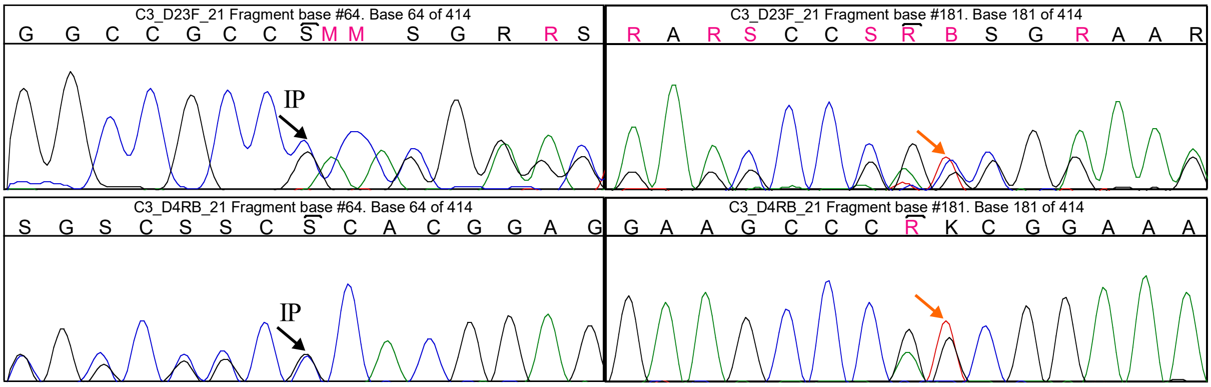


***Figure 5:*** *occurrence of three alleles with one length variant. An inflexion point is encountered (IP, black arrows) as in Figure 2, but a triple peak is observable (orange arrows), here a B in front of a K in one offset and a C in the other offset (not shown), each offset corresponding to one or more allele(s), the two offsets explaining the three peaks (a B standing for C, T and G, while the K in the shown offset stands for the T and G, the C in the other offset is consistent).*

- **CASE 7:** heterozygosity (3 alleles), **several length variants**. The chromatograms include two inflexion points but different from case 4, with occurrence of double peaks after the first inflexion point then triple peaks after the second one (**Figure 6**). This case is really difficult to solve using CHAMPURU only. A correspondence must be made between direct sequencing chromatograms and cloning, in order to validate a set of alleles that explain all double or triple peaks displayed by direct sequencing data. It could, however, be solved by a two-step process as in case 6, using CHAMPURU and confirmed alleles from the dataset. As in all cases, validation of such alleles should only be made when all alleles are confirmed in the dataset and consistent in the individual (sampling location, population, etc.). Also, all alleles must be aligned with forward and reverse to make sure these alleles explain all double peaks from direct sequencing data. Cloning is again recommended here.

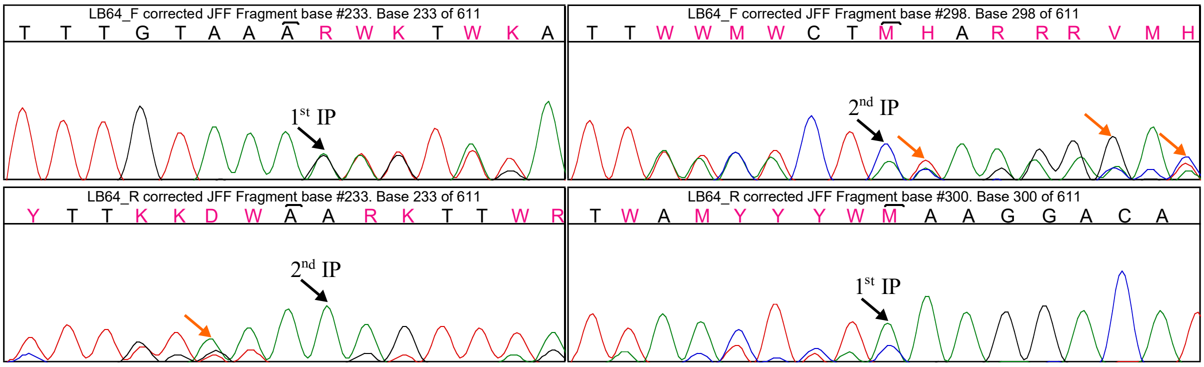


*Figure 6: Heterozygote with three alleles of different sizes. Two inflexion points are found (black arrows), corresponding to the two indels between the three alleles, with occurrence of triple peaks (orange arrows) from the second inflexion point on both chromatograms.*

Other cases are possible in theory, but are not detailed here as they were not encountered in the present work. For example, a combination of an IN*D*ELLIGENT situation (case 4) with a third allele of different size could be imagined, and would probably require cloning to be solved.

**To ensure consistency between the two methods (direct sequencing and cloning), only sequences that are explained by both direct sequencing and cloning data should be validated.** Also, some singularities deserve to be mentioned, as they could lead to important mistakes in case of wrong interpretation:

- The cases 5 to 7 (length variant heterozygotes) must be distinguished from **aspecific amplification**. Small contaminants could indeed lead to chromatograms displaying similar double peaks over a certain part of the chromatograms (example: **Figure 3**), but these contaminants would usually start at the beginning of forward and reverse chromatograms, with the last peak of the contaminant being generally of greater size and usually an A (**Figure 7**, see also Flot *et al.* 2008), while double peaks appear later in the chromatograms of length variant heterozygous and continue until the end (**Figure 2**).

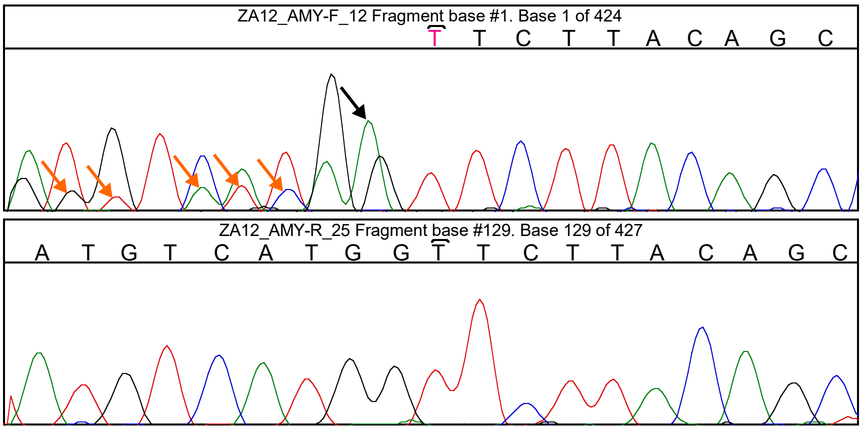


*Figure 7: aspecific amplification at the beginning of the forward chromatogram (orange arrows). A similar contamination was observed in the reverse, which could suggest a CHAMPURU situation (case 5 or 6) if the two chromatograms were inverted. However, double peaks are not expected at the very beginning of a chromatogram, and contaminant usually ends with a higher A peak (black arrow, see Flot* et al. *2008).*

- **Incongruences between forward and reverse chromatograms** must be considered carefully. A practical example is the occurrence of a double peak in one chromatogram but not the other (**Figure 8**). This could be explained by an allele dropout (one allele is amplified by only one of the two primers; in this case, this double peak must be considered) or by contamination (an aspecific product being amplified at a very low level, from which only the last peak is visible; in this case, this double peak must not be considered). This can be solved by cloning or alternatively using the information of the other sequences from closely related individuals to confirm the SNP or not.

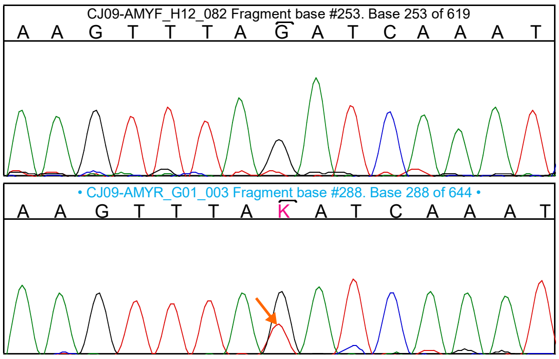


***Figure 8**: inconsistency between F and R chromatograms. At this position, the double peaks are clear on reverse sequence (orange arrow) but nothing is observed in forward sequence, except a slight background.*

- **Poor-quality regions** are also often encountered in direct sequencing data. Notably, the very beginning of these chromatograms is often of poor quality. As a consequence, the beginning and the end of the chromatograms are usually readable for only one chromatogram out of the two (respectively the reverse and the forward). This case can be solved by cloning, as the poor-quality region fall into the vector sequence giving an insert sequence of good quality from the beginning to the end. Alternatively, a hypothetic sequence can be constructed directly in the chromatogram, *e.g.* using a sequence from cloning or from a closely related individual for which this region is clean, then this hypothesis has to be tested using CHAMPURU and the information from the other chromatograms (Figure 9).

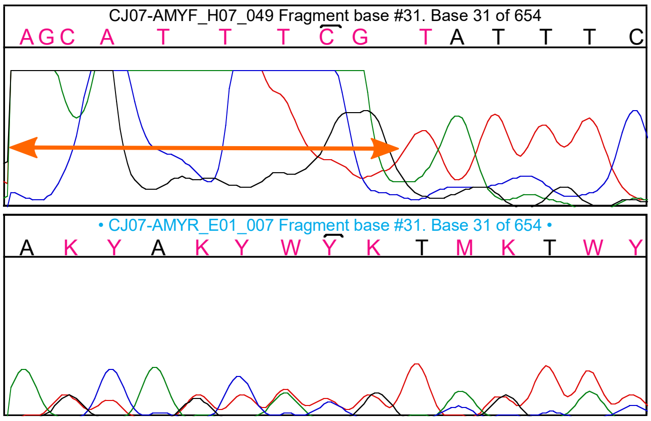


*Figure 9: poor-quality region (orange double arrows) correction. Here, the sequence was deduced from the information of the reverse chromatogram.*

In conclusion, it appears that the two methods we applied gave good results if combined: direct sequencing combined with phasing tools designed for diploids in addition to cloning sequencing (prone to contamination and recombined sequences) can retrieve confidently alleles from polyploid organisms. We strongly recommend combining them, and hope the previous details will help future studies.
