## Supplementary material for "Substantial genetic mixing among sexual and androgenetic lineages within the clam genus *Corbicula*": Table S1

GenBank accession numbers of all *Corbicula* haplotypes used in this study. The haplotype designations refer to Table S2 for further information.

| Haplotypes | Accession number | Haplotypes | Accession number |
|---|---|---|---|
| 28S-AA11a | OM905104 | ATPS-EHM102a | OM959680 |
| 28S-AA12a | OM905105 | ATPS-EHM103a | OM959681 |
| 28S-AA13a | OM905106 | ATPS-EHM104a | OM959682 |
| 28S-AA14a | OM905107 | ATPS-EHM105a | OM959683 |
| 28S-C1b | OM905108 | ATPS-EHM106 | OM959684 |
| 28S-C2b | OM905109 | ATPS-EHM107a | OM959685 |
| 28S-C3b | OM905110 | ATPS-EHM108a | OM959686 |
| 28S-C4b | OM905111 | ATPS-EHM109a | OM959687 |
| 28S-CFY01b | OM905112 | ATPS-EHM110a | OM959688 |
| 28S-CFY02b | OM905113 | ATPS-EHM111a | OM959689 |
| 28S-CFY03b | OM905114 | ATPS-EHM112a | OM959690 |
| 28S-CFY04b | OM905115 | ATPS-EHM113a | OM959691 |
| 28S-CFY05b | OM905116 | ATPS-EHM114a | OM959692 |
| 28S-CFY07b | OM905117 | ATPS-EHM115 | OM959693 |
| 28S-CFY08b | OM905118 | ATPS-EHM116a | OM959694 |
| 28S-CFY09b | OM905119 | ATPS-EHM119a | OM959695 |
| 28S-CFY10b | OM905120 | ATPS-EHM120a | OM959696 |
| 28S-CFY11b | OM905121 | ATPS-EHM121a | OM959697 |
| 28S-CFY12b | OM905122 | ATPS-EHM122a | OM959698 |
| 28S-Cl201b | OM905123 | ATPS-CFG14b | OM959699 |
| 28S-Cl202b | OM905124 | ATPS-CFG16b | OM959700 |
| 28S-Hw2c | OM905125 | ATPS-CFG17b | OM959701 |
| 28S-Hw3c | OM905126 | ATPS-CFG18b | OM959702 |
| 28S-Hw4d | OM905127 | ATPS-CFG19b | OM959703 |
| 28S-Hw5d | OM905128 | ATPS-CFG20b | OM959704 |
| 28S-Hw6d | OM905129 | ATPS-CFG21b | OM959705 |
| 28S-Muj02c | OM905130 | ATPS-CFG22b | OM959706 |
| 28S-Sa22b | OM905131 | ATPS-CFG23b | OM959707 |
| 28S-Sa23b | OM905132 | ATPS-LB36 | OM959708 |
| 28S-Sa24b | OM905133 | ATPS-LB77 | OM959709 |
| 28S-Sa25b | OM905134 | ATPS-Db22b | OM959710 |
| 28S-Sa26b | OM905135 | ATPS-Db23b | OM959711 |
| 28S-Su1a | OM905136 | ATPS-Db24b | OM959712 |
| 28S-Su2a | OM905137 | ATPS-Db25b | OM959713 |
| 28S-Su3a | OM905138 | ATPS-Db26b | OM959714 |
| 28S-Su4a | OM905139 | ATPS-KMT201a | OM959715 |
| 28S-Su5a | OM905140 | ATPS-KMT202a | OM959716 |
| 28S-U3b | OM905141 | ATPS-KMT203a | OM959717 |
| 28S-Wa1a | OM905142 | ATPS-KMT204a | OM959718 |
| 28S-Wa2a | OM905143 | ATPS-KMT205a | OM959719 |
| 28S-ccc7c | OM905144 | ATPS-KMT206a | OM959720 |
| 28S-ggg2b | OM905145 | ATPS-KMT207a | OM959721 |

| | | | |
|---|---|---|---|
| 28S-Cl201c | OM905146 | ATPS-KMT208a | OM959722 |
| 28S-Cl202c | OM905147 | ATPS-KMT209a | OM959723 |
| 28S-C1c | OM905148 | ATPS-KMT210a | OM959724 |
| 28S-C2c | OM905149 | ATPS-LB74b | OM959725 |
| 28S-C3c | OM905150 | ATPS-AA11b | OM959726 |
| 28S-C4c | OM905151 | ATPS-AA12b | OM959727 |
| 28S-Sa22c | OM905152 | ATPS-AA13b | OM959728 |
| 28S-Sa23c | OM905153 | ATPS-C1b | OM959729 |
| 28S-Sa24c | OM905154 | ATPS-C2b | OM959730 |
| 28S-Sa25c | OM905155 | ATPS-C3b | OM959731 |
| 28S-Sa26c | OM905156 | ATPS-C4b | OM959732 |
| 28S-Wa1b | OM905157 | ATPS-FU4c | OM959733 |
| 28S-Wa2b | OM905158 | ATPS-FU5c | OM959734 |
| 28S-ZA01a | OM905159 | ATPS-FU6c | OM959735 |
| 28S-ZA03a | OM905160 | ATPS-Go1 | OM959736 |
| 28S-ZA04a | OM905161 | ATPS-Hw2c | OM959737 |
| 28S-ZA05a | OM905162 | ATPS-Hw4c | OM959738 |
| 28S-ZA06a | OM905163 | ATPS-Sa22b | OM959739 |
| 28S-ZA07a | OM905164 | ATPS-Sa72b | OM959740 |
| 28S-ZA08a | OM905165 | ATPS-Sa73b | OM959741 |
| 28S-ZA09a | OM905166 | ATPS-Sa74b | OM959742 |
| 28S-ZA10a | OM905167 | ATPS-Sa75b | OM959743 |
| 28S-ZA11a | OM905168 | ATPS-Wa1b | OM959744 |
| 28S-ZA12a | OM905169 | ATPS-Wa2b | OM959745 |
| 28S-ZA13a | OM905170 | ATPS-U3a | OM959746 |
| 28S-ZA14a | OM905171 | ATPS-AB11b | OM959747 |
| 28S-ZA15a | OM905172 | ATPS-AB12b | OM959748 |
| 28S-ZA16a | OM905173 | ATPS-AB13b | OM959749 |
| 28S-ZA17a | OM905174 | ATPS-AB14b | OM959750 |
| 28S-ZA18a | OM905175 | ATPS-AB15b | OM959751 |
| 28S-ZA19a | OM905176 | ATPS-BA5a | OM959752 |
| 28S-ZA01c | OM905177 | ATPS-FU4b | OM959753 |
| 28S-ZA03b | OM905178 | ATPS-FU5b | OM959754 |
| 28S-ZA04b | OM905179 | ATPS-FU6b | OM959755 |
| 28S-ZA05b | OM905180 | ATPS-AB11a | OM959756 |
| 28S-ZA06b | OM905181 | ATPS-AB12a | OM959757 |
| 28S-ZA07b | OM905182 | ATPS-AB13a | OM959758 |
| 28S-ZA08b | OM905183 | ATPS-AB14a | OM959759 |
| 28S-ZA09b | OM905184 | ATPS-AB15a | OM959760 |
| 28S-ZA10b | OM905185 | ATPS-CFY01 | OM959761 |
| 28S-ZA11b | OM905186 | ATPS-CFY02 | OM959762 |
| 28S-ZA12b | OM905187 | ATPS-CFY03 | OM959763 |
| 28S-ZA13b | OM905188 | ATPS-CFY04 | OM959764 |
| 28S-ZA14b | OM905189 | ATPS-CFY05 | OM959765 |
| 28S-ZA15b | OM905190 | ATPS-CFY07 | OM959766 |
| 28S-ZA16b | OM905191 | ATPS-CFY08 | OM959767 |
| 28S-ZA17b | OM905192 | ATPS-CFY09 | OM959768 |
| 28S-ZA18b | OM905193 | ATPS-CFY10 | OM959769 |

| | | | |
|---|---|---|---|
| 28S-ZA19b | OM905194 | ATPS-CFY11 | OM959770 |
| 28S-ZA01b | OM905195 | ATPS-CFY12 | OM959771 |
| 28S-C1a | OM905196 | ATPS-Cl204 | OM959772 |
| 28S-C2a | OM905197 | ATPS-Cl205 | OM959773 |
| 28S-C3a | OM905198 | ATPS-Cl206 | OM959774 |
| 28S-C4a | OM905199 | ATPS-Cl207 | OM959775 |
| 28S-Sa22a | OM905200 | ATPS-Cl208 | OM959776 |
| 28S-Sa23a | OM905201 | ATPS-Cl209 | OM959777 |
| 28S-Sa24a | OM905202 | ATPS-Cl210 | OM959778 |
| 28S-Sa25a | OM905203 | ATPS-Cl211 | OM959779 |
| 28S-Sa26a | OM905204 | ATPS-Cl212 | OM959780 |
| 28S-U3c | OM905205 | ATPS-Cl213 | OM959781 |
| 28S-Wa1c | OM905206 | ATPS-Cl214 | OM959782 |
| 28S-Wa2c | OM905207 | ATPS-Cl215 | OM959783 |
| 28S-U3a | OM905208 | ATPS-Cl216 | OM959784 |
| 28S-AA11b | OM905209 | ATPS-Cl217 | OM959785 |
| 28S-AA12b | OM905210 | ATPS-Cl218 | OM959786 |
| 28S-AA13b | OM905211 | ATPS-Cl220 | OM959787 |
| 28S-AA14b | OM905212 | ATPS-Cl221 | OM959788 |
| 28S-CFG16b | OM905213 | ATPS-Cl222 | OM959789 |
| 28S-CFG19b | OM905214 | ATPS-Cl223 | OM959790 |
| 28S-CFG22b | OM905215 | ATPS-Cl224 | OM959791 |
| 28S-CFG23b | OM905216 | ATPS-Hw4d | OM959792 |
| 28S-CFY01a | OM905217 | ATPS-KMT211c | OM959793 |
| 28S-CFY02a | OM905218 | ATPS-KMT212c | OM959794 |
| 28S-CFY03a | OM905219 | ATPS-KMT213c | OM959795 |
| 28S-CFY04a | OM905220 | ATPS-KMT214c | OM959796 |
| 28S-CFY05a | OM905221 | ATPS-KMT215c | OM959797 |
| 28S-CFY07a | OM905222 | ATPS-KMT216c | OM959798 |
| 28S-CFY08a | OM905223 | ATPS-KMT217c | OM959799 |
| 28S-CFY09a | OM905224 | ATPS-KMT218c | OM959800 |
| 28S-CFY10a | OM905225 | ATPS-KMT219c | OM959801 |
| 28S-CFY11a | OM905226 | ATPS-KMT220c | OM959802 |
| 28S-CFY12a | OM905227 | ATPS-KMT221c | OM959803 |
| 28S-Db22 | OM905228 | ATPS-KMT222c | OM959804 |
| 28S-Db23 | OM905229 | ATPS-KMT223c | OM959805 |
| 28S-Db24a | OM905230 | ATPS-KMT224c | OM959806 |
| 28S-Db25 | OM905231 | ATPS-yy12 | OM959807 |
| 28S-Db26 | OM905232 | ATPS-266691b | OM959808 |
| 28S-Hw2b | OM905233 | ATPS-CFG01b | OM959809 |
| 28S-Hw3b | OM905234 | ATPS-CFG02b | OM959810 |
| 28S-Hw4c | OM905235 | ATPS-CFG03b | OM959811 |
| 28S-Hw5c | OM905236 | ATPS-CFG04b | OM959812 |
| 28S-Hw6c | OM905237 | ATPS-CFG05b | OM959813 |
| 28S-Muj02b | OM905238 | ATPS-CFG06b | OM959814 |
| 28S-Su1b | OM905239 | ATPS-CFG07b | OM959815 |
| 28S-Su2b | OM905240 | ATPS-CFG08b | OM959816 |
| 28S-Su3b | OM905241 | ATPS-CFG09b | OM959817 |

| | | | |
|---|---|---|---|
| 28S-Su4b | OM905242 | ATPS-CFG10b | OM959818 |
| 28S-Su5b | OM905243 | ATPS-CFG11b | OM959819 |
| 28S-ccc7b | OM905244 | ATPS-CFG12b | OM959820 |
| 28S-ggg2c | OM905245 | ATPS-CFG13b | OM959821 |
| 28S-Hw4b | OM905246 | ATPS-CFG15b | OM959822 |
| 28S-Hw5b | OM905247 | ATPS-CFY06b | OM959823 |
| 28S-Hw6b | OM905248 | ATPS-KMT211b | OM959824 |
| 28S-AA11c | OM905249 | ATPS-KMT212b | OM959825 |
| 28S-AA12c | OM905250 | ATPS-KMT213b | OM959826 |
| 28S-AA13c | OM905251 | ATPS-KMT215b | OM959827 |
| 28S-AA14c | OM905252 | ATPS-KMT216b | OM959828 |
| 28S-BA5b | OM905253 | ATPS-KMT217b | OM959829 |
| 28S-BA6b | OM905254 | ATPS-KMT218b | OM959830 |
| 28S-BA7b | OM905255 | ATPS-KMT219b | OM959831 |
| 28S-BA8b | OM905256 | ATPS-KMT220b | OM959832 |
| 28S-CS04a | OM905257 | ATPS-KMT221b | OM959833 |
| 28S-CS04c | OM905258 | ATPS-KMT222b | OM959834 |
| 28S-CS06c | OM905259 | ATPS-KMT223b | OM959835 |
| 28S-CS09a | OM905260 | ATPS-KMT224b | OM959836 |
| 28S-CS09c | OM905261 | ATPS-LB08 | OM959837 |
| 28S-CS10a | OM905262 | ATPS-LB13 | OM959838 |
| 28S-CS12c | OM905263 | ATPS-LB16 | OM959839 |
| 28S-CS13c | OM905264 | ATPS-LB19 | OM959840 |
| 28S-CS14c | OM905265 | ATPS-LB31 | OM959841 |
| 28S-CS15c | OM905266 | ATPS-LB40 | OM959842 |
| 28S-CS19c | OM905267 | ATPS-LB43 | OM959843 |
| 28S-CS21c | OM905268 | ATPS-LB44 | OM959844 |
| 28S-CS22c | OM905269 | ATPS-LB47 | OM959845 |
| 28S-CS23b | OM905270 | ATPS-LB66 | OM959846 |
| 28S-CS25b | OM905271 | ATPS-LB68 | OM959847 |
| 28S-CS26a | OM905272 | ATPS-LB76 | OM959848 |
| 28S-CS26c | OM905273 | ATPS-LB91 | OM959849 |
| 28S-Cl204a | OM905274 | ATPS-BA5b | OM959850 |
| 28S-Cl205a | OM905275 | ATPS-BA8h | OM959851 |
| 28S-Cl206a | OM905276 | ATPS-FU4a | OM959852 |
| 28S-Cl207a | OM905277 | ATPS-FU5a | OM959853 |
| 28S-Cl208a | OM905278 | ATPS-FU6a | OM959854 |
| 28S-Cl210a | OM905279 | ATPS-LB04b | OM959855 |
| 28S-Cl211a | OM905280 | ATPS-LB20 | OM959856 |
| 28S-Cl212a | OM905281 | ATPS-LB48 | OM959857 |
| 28S-Cl213a | OM905282 | ATPS-LB59 | OM959858 |
| 28S-Cl214a | OM905283 | ATPS-LB64 | OM959859 |
| 28S-Cl215a | OM905284 | ATPS-LB67 | OM959860 |
| 28S-Cl216a | OM905285 | ATPS-CFG01a | OM959861 |
| 28S-Cl218a | OM905286 | ATPS-CFG02a | OM959862 |
| 28S-Cl220a | OM905287 | ATPS-CFG03a | OM959863 |
| 28S-Cl221a | OM905288 | ATPS-CFG04a | OM959864 |
| 28S-Cl222a | OM905289 | ATPS-CFG05a | OM959865 |

| | | | |
|---|---|---|---|
| 28S-Cl223a | OM905290 | ATPS-CFG06a | OM959866 |
| 28S-Cl224a | OM905291 | ATPS-CFG07a | OM959867 |
| 28S-Db24b | OM905292 | ATPS-CFG08a | OM959868 |
| 28S-Hw2a | OM905293 | ATPS-CFG09a | OM959869 |
| 28S-Hw3a | OM905294 | ATPS-CFG10a | OM959870 |
| 28S-Mad2c | OM905295 | ATPS-CFG11a | OM959871 |
| 28S-Mad3c | OM905296 | ATPS-CFG12a | OM959872 |
| 28S-Mad6c | OM905297 | ATPS-CFG13a | OM959873 |
| 28S-Su1c | OM905298 | ATPS-CFG15a | OM959874 |
| 28S-Su2c | OM905299 | ATPS-CFY06a | OM959875 |
| 28S-Su3c | OM905300 | ATPS-Db22a | OM959876 |
| 28S-Su4c | OM905301 | ATPS-Db23a | OM959877 |
| 28S-Su5c | OM905302 | ATPS-Db24a | OM959878 |
| 28S-Wa1d | OM905303 | ATPS-Db25a | OM959879 |
| 28S-ccc7a | OM905304 | ATPS-Db26a | OM959880 |
| 28S-CS04b | OM905305 | ATPS-EHM101b | OM959881 |
| 28S-CS06a | OM905306 | ATPS-EHM102b | OM959882 |
| 28S-CS10c | OM905307 | ATPS-EHM103b | OM959883 |
| 28S-CS11h | OM905308 | ATPS-EHM104b | OM959884 |
| 28S-CS12a | OM905309 | ATPS-EHM105b | OM959885 |
| 28S-CS13a | OM905310 | ATPS-EHM107b | OM959886 |
| 28S-CS14a | OM905311 | ATPS-EHM108b | OM959887 |
| 28S-CS15a | OM905312 | ATPS-EHM109b | OM959888 |
| 28S-CS18 | OM905313 | ATPS-EHM110b | OM959889 |
| 28S-CS19a | OM905314 | ATPS-EHM111b | OM959890 |
| 28S-CS21a | OM905315 | ATPS-EHM112b | OM959891 |
| 28S-CS22a | OM905316 | ATPS-EHM113b | OM959892 |
| 28S-CS24a | OM905317 | ATPS-EHM114b | OM959893 |
| 28S-CS26b | OM905318 | ATPS-EHM116b | OM959894 |
| 28S-CS27 | OM905319 | ATPS-EHM119b | OM959895 |
| 28S-CS31a | OM905320 | ATPS-EHM120b | OM959896 |
| 28S-CS32 | OM905321 | ATPS-EHM121b | OM959897 |
| 28S-BA2b | OM905322 | ATPS-EHM122b | OM959898 |
| 28S-BA3b | OM905323 | ATPS-LB04a | OM959899 |
| 28S-BA4a | OM905324 | ATPS-LB22b | OM959900 |
| 28S-BA5c | OM905325 | ATPS-LB58 | OM959901 |
| 28S-Hw4a | OM905326 | ATPS-AA11a | OM959902 |
| 28S-Hw5a | OM905327 | ATPS-AA12a | OM959903 |
| 28S-Hw6a | OM905328 | ATPS-AA13a | OM959904 |
| 28S-KMT201a | OM905329 | ATPS-Hw2a | OM959905 |
| 28S-KMT202b | OM905330 | ATPS-Hw4a | OM959906 |
| 28S-KMT203b | OM905331 | ATPS-CR01a | OM959907 |
| 28S-KMT204b | OM905332 | ATPS-CR04b | OM959908 |
| 28S-KMT205b | OM905333 | ATPS-CR05b | OM959909 |
| 28S-KMT206b | OM905334 | ATPS-CR09b | OM959910 |
| 28S-KMT207b | OM905335 | ATPS-CR13 | OM959911 |
| 28S-KMT208b | OM905336 | ATPS-CR14a | OM959912 |
| 28S-KMT209b | OM905337 | ATPS-CR17a | OM959913 |

| | | | |
|---|---|---|---|
| 28S-KMT210b | OM905338 | ATPS-CR18a | OM959914 |
| 28S-BA5a | OM905339 | ATPS-CR21b | OM959915 |
| 28S-BA6a | OM905340 | ATPS-CR23b | OM959916 |
| 28S-BA7a | OM905341 | ATPS-CR24a | OM959917 |
| 28S-BA8a | OM905342 | ATPS-CRB01a | OM959918 |
| 28S-FU4c | OM905343 | ATPS-CRB04a | OM959919 |
| 28S-FU5c | OM905344 | ATPS-CRB06a | OM959920 |
| 28S-FU6c | OM905345 | ATPS-CRB10a | OM959921 |
| 28S-KMT211b | OM905346 | ATPS-CRB12b | OM959922 |
| 28S-KMT212b | OM905347 | ATPS-CRB14 | OM959923 |
| 28S-KMT213b | OM905348 | ATPS-CRB18a | OM959924 |
| 28S-KMT214b | OM905349 | ATPS-CRB20 | OM959925 |
| 28S-KMT215b | OM905350 | ATPS-Vt01a | OM959926 |
| 28S-KMT216b | OM905351 | ATPS-Vt02 | OM959927 |
| 28S-KMT217b | OM905352 | ATPS-Vt03a | OM959928 |
| 28S-KMT218b | OM905353 | ATPS-Vt10b | OM959929 |
| 28S-KMT219b | OM905354 | ATPS-Vt12 | OM959930 |
| 28S-KMT221b | OM905355 | ATPS-Vt13a | OM959931 |
| 28S-KMT222b | OM905356 | ATPS-Vt14a | OM959932 |
| 28S-KMT224b | OM905357 | ATPS-Vt15a | OM959933 |
| 28S-Muj08d | OM905358 | ATPS-Vt16a | OM959934 |
| 28S-Muj10d | OM905359 | ATPS-Vt19 | OM959935 |
| 28S-yy12c | OM905360 | ATPS-CRB01b | OM959936 |
| 28S-BA8c | OM905361 | ATPS-CRB04b | OM959937 |
| 28S-CS23a | OM905362 | ATPS-CRB13b | OM959938 |
| 28S-CS25a | OM905363 | ATPS-CRB18b | OM959939 |
| 28S-CS06b | OM905364 | ATPS-Vt10a | OM959940 |
| 28S-CS10b | OM905365 | ATPS-Vt17a | OM959941 |
| 28S-CS12b | OM905366 | ATPS-CR01b | OM959942 |
| 28S-CS13b | OM905367 | ATPS-CR02 | OM959943 |
| 28S-CS14b | OM905368 | ATPS-CR03 | OM959944 |
| 28S-CS15b | OM905369 | ATPS-CR04a | OM959945 |
| 28S-CS19b | OM905370 | ATPS-CR05a | OM959946 |
| 28S-CS21b | OM905371 | ATPS-CR09a | OM959947 |
| 28S-CS22b | OM905372 | ATPS-CR14b | OM959948 |
| 28S-CS23c | OM905373 | ATPS-CR15a | OM959949 |
| 28S-CS25c | OM905374 | ATPS-CR15b | OM959950 |
| 28S-AB11a | OM905375 | ATPS-CR17b | OM959951 |
| 28S-AB13a | OM905376 | ATPS-CR18b | OM959952 |
| 28S-AB14a | OM905377 | ATPS-CR20 | OM959953 |
| 28S-AB15a | OM905378 | ATPS-CR21a | OM959954 |
| 28S-AB16a | OM905379 | ATPS-CR23a | OM959955 |
| 28S-BA2a | OM905380 | ATPS-CR24b | OM959956 |
| 28S-BA3a | OM905381 | ATPS-CR25 | OM959957 |
| 28S-CFG14 | OM905382 | ATPS-CR26a | OM959958 |
| 28S-CFG16a | OM905383 | ATPS-CRB05b | OM959959 |
| 28S-CFG17 | OM905384 | ATPS-CRB06b | OM959960 |
| 28S-CFG18 | OM905385 | ATPS-CRB10b | OM959961 |

| | | | |
|---|---|---|---|
| 28S-CFG19a | OM905386 | ATPS-CRB13a | OM959962 |
| 28S-CFG20 | OM905387 | ATPS-CRB16a | OM959963 |
| 28S-CFG21 | OM905388 | ATPS-CRB16b | OM959964 |
| 28S-CFG22a | OM905389 | ATPS-CRB19b | OM959965 |
| 28S-CFG23a | OM905390 | ATPS-Vt01b | OM959966 |
| 28S-EHM102 | OM905391 | ATPS-Vt03b | OM959967 |
| 28S-EHM103 | OM905392 | ATPS-Vt07 | OM959968 |
| 28S-EHM104 | OM905393 | ATPS-Vt11 | OM959969 |
| 28S-EHM105 | OM905394 | ATPS-Vt13b | OM959970 |
| 28S-EHM106 | OM905395 | ATPS-Vt14b | OM959971 |
| 28S-EHM107 | OM905396 | ATPS-Vt16b | OM959972 |
| 28S-EHM108 | OM905397 | ATPS-Vt17b | OM959973 |
| 28S-EHM109 | OM905398 | ATPS-Vt15b | OM959974 |
| 28S-EHM110 | OM905399 | ATPS-LB22a | OM959975 |
| 28S-EHM111 | OM905400 | ATPS-LB74a | OM959976 |
| 28S-EHM112 | OM905401 | ATPS-C1a | OM959977 |
| 28S-EHM113 | OM905402 | ATPS-C2a | OM959978 |
| 28S-EHM119 | OM905403 | ATPS-C3a | OM959979 |
| 28S-EHM120 | OM905404 | ATPS-C4a | OM959980 |
| 28S-EHM121 | OM905405 | ATPS-Sa22a | OM959981 |
| 28S-EHM122 | OM905406 | ATPS-Sa72a | OM959982 |
| 28S-EHM123 | OM905407 | ATPS-Sa73a | OM959983 |
| 28S-EHM124 | OM905408 | ATPS-Sa74a | OM959984 |
| 28S-KMT202a | OM905409 | ATPS-Sa75a | OM959985 |
| 28S-KMT203a | OM905410 | ATPS-U3b | OM959986 |
| 28S-KMT204a | OM905411 | ATPS-Wa1a | OM959987 |
| 28S-KMT205a | OM905412 | ATPS-Wa2a | OM959988 |
| 28S-KMT206a | OM905413 | ATPS-ZA01 | OM959989 |
| 28S-KMT207a | OM905414 | ATPS-ZA02 | OM959990 |
| 28S-KMT208a | OM905415 | ATPS-ZA04 | OM959991 |
| 28S-KMT209a | OM905416 | ATPS-ZA05 | OM959992 |
| 28S-KMT210a | OM905417 | ATPS-ZA06 | OM959993 |
| 28S-KMT211a | OM905418 | ATPS-ZA08 | OM959994 |
| 28S-KMT212a | OM905419 | ATPS-ZA09 | OM959995 |
| 28S-KMT213a | OM905420 | ATPS-ZA10 | OM959996 |
| 28S-KMT214a | OM905421 | ATPS-ZA11 | OM959997 |
| 28S-KMT215a | OM905422 | ATPS-ZA12 | OM959998 |
| 28S-KMT216a | OM905423 | ATPS-ZA13 | OM959999 |
| 28S-KMT217a | OM905424 | ATPS-ZA14 | OM960000 |
| 28S-KMT218a | OM905425 | ATPS-ZA15 | OM960001 |
| 28S-KMT219a | OM905426 | ATPS-ZA16 | OM960002 |
| 28S-KMT221a | OM905427 | ATPS-ZA17 | OM960003 |
| 28S-KMT222a | OM905428 | ATPS-ZA18 | OM960004 |
| 28S-KMT224a | OM905429 | ATPS-ZA19 | OM960005 |
| 28S-Mad2b | OM905430 | ATPS-ZA20 | OM960006 |
| 28S-Mad3b | OM905431 | ATPS-fff2 | OM960007 |
| 28S-Mad6b | OM905432 | ATPS-CR26b | OM960008 |
| 28S-Muj08a | OM905433 | ATPS-CRB05a | OM960009 |

| | | | |
|---|---|---|---|
| 28S-Muj10a | OM905434 | ATPS-CRB12a | OM960010 |
| 28S-yy12a | OM905435 | ATPS-CRB19a | OM960011 |
| 28S-Cl201a | OM905436 | ATPS-LB12 | OM960012 |
| 28S-Cl202a | OM905437 | ATPS-LB62 | OM960013 |
| 28S-CR20a | OM905438 | ATPS-KMT201b | OM960014 |
| 28S-Cl204d | OM905439 | ATPS-KMT202b | OM960015 |
| 28S-Cl205d | OM905440 | ATPS-KMT203b | OM960016 |
| 28S-Cl206d | OM905441 | ATPS-KMT204b | OM960017 |
| 28S-Cl207d | OM905442 | ATPS-KMT205b | OM960018 |
| 28S-Cl208d | OM905443 | ATPS-KMT206b | OM960019 |
| 28S-Cl210d | OM905444 | ATPS-KMT207b | OM960020 |
| 28S-Cl211d | OM905445 | ATPS-KMT208b | OM960021 |
| 28S-Cl212d | OM905446 | ATPS-KMT209b | OM960022 |
| 28S-Cl213d | OM905447 | ATPS-KMT210b | OM960023 |
| 28S-Cl214d | OM905448 | ATPS-KMT211a | OM960024 |
| 28S-Cl215d | OM905449 | ATPS-KMT212a | OM960025 |
| 28S-Cl216d | OM905450 | ATPS-KMT213a | OM960026 |
| 28S-Cl218d | OM905451 | ATPS-KMT214a | OM960027 |
| 28S-Cl220d | OM905452 | ATPS-KMT215a | OM960028 |
| 28S-Cl221d | OM905453 | ATPS-KMT216a | OM960029 |
| 28S-Cl222d | OM905454 | ATPS-KMT217a | OM960030 |
| 28S-Cl223d | OM905455 | ATPS-KMT218a | OM960031 |
| 28S-Cl224d | OM905456 | ATPS-KMT219a | OM960032 |
| 28S-CR01a | OM905457 | ATPS-KMT220a | OM960033 |
| 28S-CR02a | OM905458 | ATPS-KMT221a | OM960034 |
| 28S-CRB14h | OM905459 | ATPS-KMT222a | OM960035 |
| 28S-Vt02a | OM905460 | ATPS-KMT223a | OM960036 |
| 28S-Vt20a | OM905461 | ATPS-KMT224a | OM960037 |
| 28S-CR04a | OM905462 | ATPS-LB69 | OM960038 |
| 28S-CR06h | OM905463 | ATPS-LB72 | OM960039 |
| 28S-CR12a | OM905464 | ATPS-CJ02a | OM960040 |
| 28S-CR16a | OM905465 | ATPS-CJ02b | OM960041 |
| 28S-CR18a | OM905466 | ATPS-CJ03a | OM960042 |
| 28S-CR22a | OM905467 | ATPS-CJ04a | OM960043 |
| 28S-CR24b | OM905468 | ATPS-CJ04b | OM960044 |
| 28S-CRB05a | OM905469 | ATPS-CJ06a | OM960045 |
| 28S-CRB07a | OM905470 | ATPS-CJ06b | OM960046 |
| 28S-Vt01a | OM905471 | ATPS-CJ09a | OM960047 |
| 28S-Vt02b | OM905472 | ATPS-CJ09b | OM960048 |
| 28S-Vt12 | OM905473 | ATPS-CJ10 | OM960049 |
| 28S-Vt13a | OM905474 | ATPS-CJ11a | OM960050 |
| 28S-Vt14a | OM905475 | ATPS-CJ11b | OM960051 |
| 28S-Vt01b | OM905476 | ATPS-CJ13 | OM960052 |
| 28S-CR25a | OM905477 | ATPS-CJ15b | OM960053 |
| 28S-CRB02a | OM905478 | ATPS-CJ19 | OM960054 |
| 28S-CRB03a | OM905479 | ATPS-CJ22a | OM960055 |
| 28S-CRB09a | OM905480 | ATPS-Jp03a | OM960056 |
| 28S-CRB10a | OM905481 | ATPS-Jp03b | OM960057 |

| | | | |
|---|---|---|---|
| 28S-Vt03a | OM905482 | ATPS-Jp04 | OM960058 |
| 28S-Vt08a | OM905483 | ATPS-Jp06b | OM960059 |
| 28S-Vt15a | OM905484 | ATPS-Jp08a | OM960060 |
| 28S-Vt05a | OM905485 | ATPS-Jp08b | OM960061 |
| 28S-Vt11a | OM905486 | ATPS-Jp10a | OM960062 |
| 28S-CFG01b | OM905487 | ATPS-Jp10b | OM960063 |
| 28S-CFG02b | OM905488 | ATPS-Jp06a | OM960064 |
| 28S-CFG03b | OM905489 | ATPS-CJ14b | OM960065 |
| 28S-CFG04b | OM905490 | ATPS-CJ18b | OM960066 |
| 28S-CFG05b | OM905491 | ATPS-CJ18a | OM960067 |
| 28S-CFG06b | OM905492 | ATPS-CJ03b | OM960068 |
| 28S-CFG07b | OM905493 | ATPS-CJ07a | OM960069 |
| 28S-CFG08b | OM905494 | ATPS-CJ12a | OM960070 |
| 28S-CFG09b | OM905495 | ATPS-CJ15a | OM960071 |
| 28S-CFG10b | OM905496 | ATPS-CJ20 | OM960072 |
| 28S-CFG11b | OM905497 | ATPS-CJ22b | OM960073 |
| 28S-CFG12b | OM905498 | ATPS-CJ07b | OM960074 |
| 28S-CFG13b | OM905499 | ATPS-CJ12b | OM960075 |
| 28S-CFG15b | OM905500 | ATPS-CJ14a | OM960076 |
| 28S-CFY06b | OM905501 | ATPS-Ka3a | OM960077 |
| 28S-Muj02a | OM905502 | ATPS-Ka3b | OM960078 |
| 28S-ggg2a | OM905503 | amy-103027 | OM960079 |
| 28S-CR01b | OM905504 | amy-CR01b | OM960080 |
| 28S-CR02b | OM905505 | amy-CR03a | OM960081 |
| 28S-CR03 | OM905506 | amy-CR15b | OM960082 |
| 28S-CR04b | OM905507 | amy-CR17a | OM960083 |
| 28S-CR05b | OM905508 | amy-CRB09a | OM960084 |
| 28S-CR09b | OM905509 | amy-CRB17 | OM960085 |
| 28S-CR12b | OM905510 | amy-Vt01a | OM960086 |
| 28S-CR13a | OM905511 | amy-Vt12a | OM960087 |
| 28S-CR13b | OM905512 | amy-Vt19b | OM960088 |
| 28S-CR16b | OM905513 | amy-CR04a | OM960089 |
| 28S-CR17b | OM905514 | amy-CR05a | OM960090 |
| 28S-CR18b | OM905515 | amy-CR22b | OM960091 |
| 28S-CR19b | OM905516 | amy-Vt13b | OM960092 |
| 28S-CR20b | OM905517 | amy-CR02b | OM960093 |
| 28S-CR22b | OM905518 | amy-CR05b | OM960094 |
| 28S-CR23b | OM905519 | amy-CR13b | OM960095 |
| 28S-CR24a | OM905520 | amy-CRB14 | OM960096 |
| 28S-CR25b | OM905521 | amy-Vt10b | OM960097 |
| 28S-CR26b | OM905522 | amy-Vt11b | OM960098 |
| 28S-CRB01b | OM905523 | amy-Vt16b | OM960099 |
| 28S-CRB02b | OM905524 | amy-Vt10a | OM960100 |
| 28S-CRB03b | OM905525 | amy-Vt16a | OM960101 |
| 28S-CRB05b | OM905526 | amy-CR24b | OM960102 |
| 28S-CRB06b | OM905527 | amy-CRB12b | OM960103 |
| 28S-CRB07b | OM905528 | amy-Vt15b | OM960104 |
| 28S-CRB09b | OM905529 | amy-CRB05b | OM960105 |

| | | | |
|---|---|---|---|
| 28S-CRB10b | OM905530 | amy-Vt01b | OM960106 |
| 28S-CRB12b | OM905531 | amy-C1b | OM960107 |
| 28S-CRB13b | OM905532 | amy-C2b | OM960108 |
| 28S-CRB15 | OM905533 | amy-C3b | OM960109 |
| 28S-CRB18b | OM905534 | amy-C4b | OM960110 |
| 28S-CRB20b | OM905535 | amy-CR04b | OM960111 |
| 28S-CRB21b | OM905536 | amy-Sa22b | OM960112 |
| 28S-Vt02c | OM905537 | amy-Sa23b | OM960113 |
| 28S-Vt03b | OM905538 | amy-Sa24b | OM960114 |
| 28S-Vt05b | OM905539 | amy-Sa25b | OM960115 |
| 28S-Vt06b | OM905540 | amy-Sa26b | OM960116 |
| 28S-Vt08b | OM905541 | amy-U4a | OM960117 |
| 28S-Vt09b | OM905542 | amy-Vt03b | OM960118 |
| 28S-Vt10a | OM905543 | amy-Wa1 | OM960119 |
| 28S-Vt11b | OM905544 | amy-Wa2b | OM960120 |
| 28S-Vt13b | OM905545 | amy-ZA01 | OM960121 |
| 28S-Vt14b | OM905546 | amy-ZA05 | OM960122 |
| 28S-Vt15b | OM905547 | amy-ZA06 | OM960123 |
| 28S-Vt16b | OM905548 | amy-ZA07 | OM960124 |
| 28S-Vt17b | OM905549 | amy-ZA09 | OM960125 |
| 28S-Vt19b | OM905550 | amy-ZA10 | OM960126 |
| 28S-CR09a | OM905551 | amy-ZA11 | OM960127 |
| 28S-CR15a | OM905552 | amy-ZA12 | OM960128 |
| 28S-CR17a | OM905553 | amy-ZA13 | OM960129 |
| 28S-CR24c | OM905554 | amy-ZA15 | OM960130 |
| 28S-CR26a | OM905555 | amy-ZA16 | OM960131 |
| 28S-CRB01a | OM905556 | amy-ZA17 | OM960132 |
| 28S-CRB04a | OM905557 | amy-ZA18 | OM960133 |
| 28S-CRB12a | OM905558 | amy-ZA19 | OM960134 |
| 28S-CRB18a | OM905559 | amy-fff2a | OM960135 |
| 28S-Vt06a | OM905560 | amy-CR15a | OM960136 |
| 28S-Vt09a | OM905561 | amy-CR02a | OM960137 |
| 28S-Vt19a | OM905562 | amy-CR03b | OM960138 |
| 28S-Vt20b | OM905563 | amy-CR13a | OM960139 |
| 28S-CR15b | OM905564 | amy-CR17b | OM960140 |
| 28S-CRB04b | OM905565 | amy-CR22a | OM960141 |
| 28S-CR05a | OM905566 | amy-CRB05a | OM960142 |
| 28S-CRB13a | OM905567 | amy-CRB09b | OM960143 |
| 28S-CR19a | OM905568 | amy-CRB12a | OM960144 |
| 28S-CRB20a | OM905569 | amy-Vt02a | OM960145 |
| 28S-CRB21a | OM905570 | amy-Vt13a | OM960146 |
| 28S-Vt06c | OM905571 | amy-Vt17a | OM960147 |
| 28S-Vt16a | OM905572 | amy-Vt19a | OM960148 |
| 28S-CRB06a | OM905573 | amy-Vt20b | OM960149 |
| 28S-CR23a | OM905574 | amy-CRB04b | OM960150 |
| 28S-Vt10b | OM905575 | amy-Vt15a | OM960151 |
| 28S-Vt17a | OM905576 | amy-CR24a | OM960152 |
| 28S-CR02c | OM905577 | amy-Vt12b | OM960153 |

| | | | |
|---|---|---|---|
| 28S-CS04d | OM905578 | amy-Vt20a | OM960154 |
| 28S-CS06d | OM905579 | amy-Vt03a | OM960155 |
| 28S-CS09b | OM905580 | amy-CRB04a | OM960156 |
| 28S-CS12d | OM905581 | amy-Vt11a | OM960157 |
| 28S-CS13d | OM905582 | amy-Vt17b | OM960158 |
| 28S-CS14d | OM905583 | amy-Vt02b | OM960159 |
| 28S-CS15d | OM905584 | amy-CR01a | OM960160 |
| 28S-CS19d | OM905585 | amy-170096 | OM960161 |
| 28S-CS21d | OM905586 | amy-AA10b | OM960162 |
| 28S-CS24b | OM905587 | amy-AA11b | OM960163 |
| 28S-CS31b | OM905588 | amy-AA12b | OM960164 |
| 28S-FU4a | OM905589 | amy-AA13b | OM960165 |
| 28S-FU5a | OM905590 | amy-AA14b | OM960166 |
| 28S-FU6a | OM905591 | amy-AB11b | OM960167 |
| 28S-Muj08b | OM905592 | amy-AB12b | OM960168 |
| 28S-Muj10b | OM905593 | amy-AB13b | OM960169 |
| 28S-FU4b | OM905594 | amy-AB14b | OM960170 |
| 28S-FU5b | OM905595 | amy-AB15b | OM960171 |
| 28S-FU6b | OM905596 | amy-AB16b | OM960172 |
| 28S-AB11b | OM905597 | amy-KMT211a | OM960173 |
| 28S-AB13b | OM905598 | amy-KMT212a | OM960174 |
| 28S-AB14b | OM905599 | amy-KMT213a | OM960175 |
| 28S-AB15b | OM905600 | amy-KMT214a | OM960176 |
| 28S-AB16b | OM905601 | amy-KMT215a | OM960177 |
| 28S-CFY06c | OM905602 | amy-KMT216a | OM960178 |
| 28S-KMT201b | OM905603 | amy-KMT217a | OM960179 |
| 28S-KMT202c | OM905604 | amy-KMT218a | OM960180 |
| 28S-KMT203c | OM905605 | amy-KMT219a | OM960181 |
| 28S-KMT204c | OM905606 | amy-KMT221a | OM960182 |
| 28S-KMT205c | OM905607 | amy-KMT222a | OM960183 |
| 28S-KMT206c | OM905608 | amy-KMT224a | OM960184 |
| 28S-KMT207c | OM905609 | amy-LB11b | OM960185 |
| 28S-KMT208c | OM905610 | amy-LB64b | OM960186 |
| 28S-KMT209c | OM905611 | amy-Su1b | OM960187 |
| 28S-KMT210c | OM905612 | amy-Su2b | OM960188 |
| 28S-yy12b | OM905613 | amy-Su3b | OM960189 |
| 28S-BA4b | OM905614 | amy-Su4b | OM960190 |
| 28S-CFG01a | OM905615 | amy-Su5 | OM960191 |
| 28S-CFG02a | OM905616 | amy-ggg2a | OM960192 |
| 28S-CFG03a | OM905617 | amy-qq1c | OM960193 |
| 28S-CFG04a | OM905618 | amy-rr1b | OM960194 |
| 28S-CFG05a | OM905619 | amy-xx11b | OM960195 |
| 28S-CFG06a | OM905620 | amy-LB71b | OM960196 |
| 28S-CFG07a | OM905621 | amy-qq1a | OM960197 |
| 28S-CFG08a | OM905622 | amy-AA10a | OM960198 |
| 28S-CFG09a | OM905623 | amy-AA11a | OM960199 |
| 28S-CFG10a | OM905624 | amy-AA12a | OM960200 |
| 28S-CFG11a | OM905625 | amy-AA13a | OM960201 |

| | | | |
|---|---|---|---|
| 28S-CFG12a | OM905626 | amy-AA14a | OM960202 |
| 28S-CFG13a | OM905627 | amy-AB11a | OM960203 |
| 28S-CFG15a | OM905628 | amy-AB12a | OM960204 |
| 28S-CFY06a | OM905629 | amy-AB13a | OM960205 |
| 28S-Cl204b | OM905630 | amy-AB14a | OM960206 |
| 28S-Cl205b | OM905631 | amy-AB15a | OM960207 |
| 28S-Cl206b | OM905632 | amy-AB16a | OM960208 |
| 28S-Cl207b | OM905633 | amy-CFG01a | OM960209 |
| 28S-Cl208b | OM905634 | amy-CFG02a | OM960210 |
| 28S-Cl210b | OM905635 | amy-CFG03a | OM960211 |
| 28S-Cl211b | OM905636 | amy-CFG04a | OM960212 |
| 28S-Cl212b | OM905637 | amy-CFG05a | OM960213 |
| 28S-Cl213b | OM905638 | amy-CFG06a | OM960214 |
| 28S-Cl214b | OM905639 | amy-CFG07a | OM960215 |
| 28S-Cl215b | OM905640 | amy-CFG08a | OM960216 |
| 28S-Cl216b | OM905641 | amy-CFG09a | OM960217 |
| 28S-Cl218b | OM905642 | amy-CFG10a | OM960218 |
| 28S-Cl220b | OM905643 | amy-CFG11a | OM960219 |
| 28S-Cl221b | OM905644 | amy-CFG12a | OM960220 |
| 28S-Cl222b | OM905645 | amy-CFG13a | OM960221 |
| 28S-Cl223b | OM905646 | amy-CFG15a | OM960222 |
| 28S-Cl224b | OM905647 | amy-CFY06a | OM960223 |
| 28S-Muj08c | OM905648 | amy-CS19 | OM960224 |
| 28S-Muj10c | OM905649 | amy-CS22a | OM960225 |
| 28S-Cl204c | OM905650 | amy-CS26a | OM960226 |
| 28S-Cl205c | OM905651 | amy-LB02a | OM960227 |
| 28S-Cl206c | OM905652 | amy-LB06a | OM960228 |
| 28S-Cl207c | OM905653 | amy-LB13a | OM960229 |
| 28S-Cl208c | OM905654 | amy-LB16a | OM960230 |
| 28S-Cl210c | OM905655 | amy-LB17c | OM960231 |
| 28S-Cl211c | OM905656 | amy-LB19a | OM960232 |
| 28S-Cl212c | OM905657 | amy-LB31 | OM960233 |
| 28S-Cl213c | OM905658 | amy-LB40a | OM960234 |
| 28S-Cl214c | OM905659 | amy-LB43a | OM960235 |
| 28S-Cl215c | OM905660 | amy-LB44a | OM960236 |
| 28S-Cl216c | OM905661 | amy-LB62a | OM960237 |
| 28S-Cl218c | OM905662 | amy-LB66a | OM960238 |
| 28S-Cl220c | OM905663 | amy-LB69a | OM960239 |
| 28S-Cl221c | OM905664 | amy-LB71a | OM960240 |
| 28S-Cl222c | OM905665 | amy-LB76a | OM960241 |
| 28S-Cl223c | OM905666 | amy-Su1a | OM960242 |
| 28S-Cl224c | OM905667 | amy-Su2a | OM960243 |
| 28S-Mad2a | OM905668 | amy-Su3a | OM960244 |
| 28S-Mad3a | OM905669 | amy-Su4a | OM960245 |
| 28S-Mad6a | OM905670 | amy-qq1b | OM960246 |
| 28S-CJ01a | OM905671 | amy-xx11a | OM960247 |
| 28S-CJ05a | OM905672 | amy-CS04a | OM960248 |
| 28S-CJ06a | OM905673 | amy-CS13b | OM960249 |

| | | | |
|---|---|---|---|
| 28S-CJ07a | OM905674 | amy-CS14a | OM960250 |
| 28S-CJ08a | OM905675 | amy-CS15a | OM960251 |
| 28S-CJ09a | OM905676 | amy-CS22b | OM960252 |
| 28S-CJ10a | OM905677 | amy-CS23a | OM960253 |
| 28S-CJ11a | OM905678 | amy-CS26b | OM960254 |
| 28S-CJ12a | OM905679 | amy-CS31b | OM960255 |
| 28S-CJ13a | OM905680 | amy-CS32 | OM960256 |
| 28S-CJ14a | OM905681 | amy-LB13b | OM960257 |
| 28S-CJ15a | OM905682 | amy-LB16b | OM960258 |
| 28S-CJ16a | OM905683 | amy-LB40b | OM960259 |
| 28S-CJ19a | OM905684 | amy-LB76b | OM960260 |
| 28S-CJ20a | OM905685 | amy-CS04b | OM960261 |
| 28S-CJ21a | OM905686 | amy-CS25a | OM960262 |
| 28S-Jp01a | OM905687 | amy-CS06a | OM960263 |
| 28S-Jp02a | OM905688 | amy-CS09a | OM960264 |
| 28S-CJ17a | OM905689 | amy-CS10a | OM960265 |
| 28S-CJ01b | OM905690 | amy-CS12a | OM960266 |
| 28S-CJ02 | OM905691 | amy-CS21a | OM960267 |
| 28S-CJ03 | OM905692 | amy-CS24a | OM960268 |
| 28S-CJ04 | OM905693 | amy-CS27a | OM960269 |
| 28S-CJ05b | OM905694 | amy-CS14b | OM960270 |
| 28S-CJ06b | OM905695 | amy-CS15b | OM960271 |
| 28S-CJ07b | OM905696 | amy-CS23b | OM960272 |
| 28S-CJ08b | OM905697 | amy-CS31a | OM960273 |
| 28S-CJ09b | OM905698 | amy-LB02b | OM960274 |
| 28S-CJ10b | OM905699 | amy-LB43b | OM960275 |
| 28S-CJ11b | OM905700 | amy-EHM102a | OM960276 |
| 28S-CJ12b | OM905701 | amy-EHM103a | OM960277 |
| 28S-CJ13b | OM905702 | amy-EHM104a | OM960278 |
| 28S-CJ14b | OM905703 | amy-EHM105a | OM960279 |
| 28S-CJ15b | OM905704 | amy-EHM106a | OM960280 |
| 28S-CJ16b | OM905705 | amy-EHM107a | OM960281 |
| 28S-CJ17b | OM905706 | amy-EHM108a | OM960282 |
| 28S-CJ18 | OM905707 | amy-EHM109a | OM960283 |
| 28S-CJ19b | OM905708 | amy-EHM110a | OM960284 |
| 28S-CJ20b | OM905709 | amy-EHM111a | OM960285 |
| 28S-CJ21b | OM905710 | amy-EHM112a | OM960286 |
| 28S-CJ22 | OM905711 | amy-EHM113a | OM960287 |
| 28S-Jp01b | OM905712 | amy-EHM119a | OM960288 |
| 28S-Jp02b | OM905713 | amy-EHM120a | OM960289 |
| 28S-Jp10 | OM905714 | amy-EHM121a | OM960290 |
| COI-170096 | OM912053 | amy-EHM122a | OM960291 |
| COI-190768 | OM912054 | amy-EHM123a | OM960292 |
| COI-190852 | OM912055 | amy-EHM124a | OM960293 |
| COI-191042 | OM912056 | amy-CS13a | OM960294 |
| COI-AA10 | OM912057 | amy-AA10c | OM960295 |
| COI-AA11 | OM912058 | amy-AA11c | OM960296 |
| COI-AA12 | OM912059 | amy-AA12c | OM960297 |

| | | | |
|---|---|---|---|
| COI-AA13 | OM912060 | amy-AA13c | OM960298 |
| COI-AA14 | OM912061 | amy-AA14c | OM960299 |
| COI-AB11 | OM912062 | amy-Su1c | OM960300 |
| COI-AB12 | OM912063 | amy-Su2c | OM960301 |
| COI-AB13 | OM912064 | amy-Su3c | OM960302 |
| COI-AB14 | OM912065 | amy-Su4c | OM960303 |
| COI-AB15 | OM912066 | amy-ggg2b | OM960304 |
| COI-AB16 | OM912067 | amy-xx11c | OM960305 |
| COI-BA1 | OM912068 | amy-AB12c | OM960306 |
| COI-BA6 | OM912069 | amy-AB13c | OM960307 |
| COI-C1 | OM912070 | amy-AB14c | OM960308 |
| COI-C2 | OM912071 | amy-AB15c | OM960309 |
| COI-C3 | OM912072 | amy-AB16c | OM960310 |
| COI-C4 | OM912073 | amy-Db22b | OM960311 |
| COI-CFG01 | OM912074 | amy-Db23b | OM960312 |
| COI-CFG02 | OM912075 | amy-Db24b | OM960313 |
| COI-CFG03 | OM912076 | amy-Db25b | OM960314 |
| COI-CFG04 | OM912077 | amy-Db26b | OM960315 |
| COI-CFG05 | OM912078 | amy-yy11a | OM960316 |
| COI-CFG06 | OM912079 | amy-yy12a | OM960317 |
| COI-CFG07 | OM912080 | amy-LB44b | OM960318 |
| COI-CFG08 | OM912081 | amy-AB12d | OM960319 |
| COI-CFG09 | OM912082 | amy-AB13d | OM960320 |
| COI-CFG10 | OM912083 | amy-AB14d | OM960321 |
| COI-CFG11 | OM912084 | amy-AB15d | OM960322 |
| COI-CFG12 | OM912085 | amy-AB16d | OM960323 |
| COI-CFG13 | OM912086 | amy-EHM102b | OM960324 |
| COI-CFG15 | OM912087 | amy-EHM103b | OM960325 |
| COI-CFG16 | OM912088 | amy-EHM104b | OM960326 |
| COI-CFG17 | OM912089 | amy-EHM105b | OM960327 |
| COI-CFG18 | OM912090 | amy-EHM106b | OM960328 |
| COI-CFG19 | OM912091 | amy-EHM107b | OM960329 |
| COI-CFG20 | OM912092 | amy-EHM108b | OM960330 |
| COI-CFG21 | OM912093 | amy-EHM109b | OM960331 |
| COI-CFG22 | OM912094 | amy-EHM110b | OM960332 |
| COI-CFG23 | OM912095 | amy-EHM111b | OM960333 |
| COI-CFY03 | OM912096 | amy-EHM112b | OM960334 |
| COI-CFY04 | OM912097 | amy-EHM113b | OM960335 |
| COI-CFY05 | OM912098 | amy-EHM119b | OM960336 |
| COI-CFY07 | OM912099 | amy-EHM120b | OM960337 |
| COI-CFY09 | OM912100 | amy-EHM121b | OM960338 |
| COI-CFY10 | OM912101 | amy-EHM122b | OM960339 |
| COI-CFY11 | OM912102 | amy-EHM123b | OM960340 |
| COI-CFY12 | OM912103 | amy-EHM124b | OM960341 |
| COI-CJ01 | OM912104 | amy-yy11b | OM960342 |
| COI-CJ03 | OM912105 | amy-yy12b | OM960343 |
| COI-CJ04 | OM912106 | amy-LB66b | OM960344 |
| COI-CJ05 | OM912107 | amy-rr1a | OM960345 |

| | | | |
|---|---|---|---|
| COI-CJ07 | OM912108 | amy-LB62b | OM960346 |
| COI-CJ09 | OM912109 | amy-LB19b | OM960347 |
| COI-CJ10 | OM912110 | amy-LB20b | OM960348 |
| COI-CJ11 | OM912111 | amy-CFG01b | OM960349 |
| COI-CJ12 | OM912112 | amy-CFG02b | OM960350 |
| COI-CJ15 | OM912113 | amy-CFG03b | OM960351 |
| COI-CJ18 | OM912114 | amy-CFG04b | OM960352 |
| COI-CJ19 | OM912115 | amy-CFG05b | OM960353 |
| COI-CJ21 | OM912116 | amy-CFG06b | OM960354 |
| COI-Cl204 | OM912117 | amy-CFG07b | OM960355 |
| COI-Cl205 | OM912118 | amy-CFG08b | OM960356 |
| COI-Cl206 | OM912119 | amy-CFG09b | OM960357 |
| COI-Cl207 | OM912120 | amy-CFG10b | OM960358 |
| COI-Cl208 | OM912121 | amy-CFG11b | OM960359 |
| COI-Cl210 | OM912122 | amy-CFG12b | OM960360 |
| COI-Cl211 | OM912123 | amy-CFG13b | OM960361 |
| COI-Cl212 | OM912124 | amy-CFG14b | OM960362 |
| COI-Cl213 | OM912125 | amy-CFG15b | OM960363 |
| COI-Cl214 | OM912126 | amy-CFG16b | OM960364 |
| COI-Cl215 | OM912127 | amy-CFG17b | OM960365 |
| COI-Cl216 | OM912128 | amy-CFG18b | OM960366 |
| COI-Cl218 | OM912129 | amy-CFG19b | OM960367 |
| COI-Cl220 | OM912130 | amy-CFG20b | OM960368 |
| COI-Cl221 | OM912131 | amy-CFG22b | OM960369 |
| COI-Cl222 | OM912132 | amy-CFG23b | OM960370 |
| COI-Cl223 | OM912133 | amy-CFY01b | OM960371 |
| COI-Cl224 | OM912134 | amy-CFY02b | OM960372 |
| COI-CR01 | OM912135 | amy-CFY03b | OM960373 |
| COI-CR02 | OM912136 | amy-CFY04b | OM960374 |
| COI-CR03 | OM912137 | amy-CFY05b | OM960375 |
| COI-CR04 | OM912138 | amy-CFY06b | OM960376 |
| COI-CR05 | OM912139 | amy-CFY07b | OM960377 |
| COI-CR13 | OM912140 | amy-CFY08b | OM960378 |
| COI-CR15 | OM912141 | amy-CFY09b | OM960379 |
| COI-CR17 | OM912142 | amy-CFY10b | OM960380 |
| COI-CR22 | OM912143 | amy-CFY11b | OM960381 |
| COI-CR24 | OM912144 | amy-CFY12b | OM960382 |
| COI-CR26 | OM912145 | amy-Cl204b | OM960383 |
| COI-CRB01 | OM912146 | amy-Cl205b | OM960384 |
| COI-CRB04 | OM912147 | amy-Cl206b | OM960385 |
| COI-CRB05 | OM912148 | amy-Cl207b | OM960386 |
| COI-CRB09 | OM912149 | amy-Cl208b | OM960387 |
| COI-CRB12 | OM912150 | amy-Cl210b | OM960388 |
| COI-CRB14 | OM912151 | amy-Cl211b | OM960389 |
| COI-CRB15 | OM912152 | amy-Cl213b | OM960390 |
| COI-CRB16 | OM912153 | amy-Cl214b | OM960391 |
| COI-CS04 | OM912154 | amy-Cl215b | OM960392 |
| COI-CS06 | OM912155 | amy-Cl216b | OM960393 |

| | | | |
|---|---|---|---|
| COI-CS09 | OM912156 | amy-Cl218b | OM960394 |
| COI-CS10 | OM912157 | amy-Cl220b | OM960395 |
| COI-CS12 | OM912158 | amy-Cl221b | OM960396 |
| COI-CS13 | OM912159 | amy-Cl222b | OM960397 |
| COI-CS14 | OM912160 | amy-Cl223b | OM960398 |
| COI-CS15 | OM912161 | amy-Cl224b | OM960399 |
| COI-CS19 | OM912162 | amy-Db22a | OM960400 |
| COI-CS21 | OM912163 | amy-Db23a | OM960401 |
| COI-CS22 | OM912164 | amy-Db24a | OM960402 |
| COI-CS23 | OM912165 | amy-Db25a | OM960403 |
| COI-CS24 | OM912166 | amy-Db26a | OM960404 |
| COI-CS25 | OM912167 | amy-KMT211b | OM960405 |
| COI-CS26 | OM912168 | amy-KMT212b | OM960406 |
| COI-CS27 | OM912169 | amy-KMT213b | OM960407 |
| COI-CS31 | OM912170 | amy-KMT214b | OM960408 |
| COI-CS32 | OM912171 | amy-KMT215b | OM960409 |
| COI-Db22 | OM912172 | amy-KMT216b | OM960410 |
| COI-Db23 | OM912173 | amy-KMT217b | OM960411 |
| COI-Db24 | OM912174 | amy-KMT218b | OM960412 |
| COI-Db25 | OM912175 | amy-KMT219b | OM960413 |
| COI-Db26 | OM912176 | amy-KMT221b | OM960414 |
| COI-EHM102 | OM912177 | amy-KMT222b | OM960415 |
| COI-EHM103 | OM912178 | amy-KMT224b | OM960416 |
| COI-EHM104 | OM912179 | amy-LB17b | OM960417 |
| COI-EHM105 | OM912180 | amy-C1a | OM960418 |
| COI-EHM106 | OM912181 | amy-C2a | OM960419 |
| COI-EHM107 | OM912182 | amy-C3a | OM960420 |
| COI-EHM108 | OM912183 | amy-C4a | OM960421 |
| COI-EHM109 | OM912184 | amy-Sa22a | OM960422 |
| COI-EHM110 | OM912185 | amy-Sa23a | OM960423 |
| COI-EHM111 | OM912186 | amy-Sa24a | OM960424 |
| COI-EHM112 | OM912187 | amy-Sa25a | OM960425 |
| COI-EHM113 | OM912188 | amy-Sa26a | OM960426 |
| COI-EHM119 | OM912189 | amy-Wa2a | OM960427 |
| COI-EHM120 | OM912190 | amy-fff2b | OM960428 |
| COI-EHM121 | OM912191 | amy-LB98c | OM960429 |
| COI-EHM122 | OM912192 | amy-CS06b | OM960430 |
| COI-EHM123 | OM912193 | amy-CS09b | OM960431 |
| COI-EHM124 | OM912194 | amy-CS10b | OM960432 |
| COI-FU1 | OM912195 | amy-CS12b | OM960433 |
| COI-Jp01 | OM912196 | amy-CS21b | OM960434 |
| COI-Jp02 | OM912197 | amy-CS24b | OM960435 |
| COI-Jp03 | OM912198 | amy-CS25b | OM960436 |
| COI-Jp10 | OM912199 | amy-CS27b | OM960437 |
| COI-Ka1 | OM912200 | amy-LB68 | OM960438 |
| COI-Ka2 | OM912201 | amy-LB72b | OM960439 |
| COI-Ka3 | OM912202 | amy-LB04a | OM960440 |
| COI-Ka5 | OM912203 | amy-LB04b | OM960441 |

| | | | |
|---|---|---|---|
| COI-KMT205 | OM912204 | amy-LB59a | OM960442 |
| COI-KMT208 | OM912205 | amy-LB59b | OM960443 |
| COI-KMT213 | OM912206 | amy-LB64a | OM960444 |
| COI-KMT214 | OM912207 | amy-LB64c | OM960445 |
| COI-KMT215 | OM912208 | amy-LB98a | OM960446 |
| COI-KMT216 | OM912209 | amy-LB98b | OM960447 |
| COI-KMT217 | OM912210 | amy-U4b | OM960448 |
| COI-KMT218 | OM912211 | amy-LB14 | OM960449 |
| COI-KMT219 | OM912212 | amy-LB18b | OM960450 |
| COI-KMT221 | OM912213 | amy-LB22b | OM960451 |
| COI-KMT222 | OM912214 | amy-LB42b | OM960452 |
| COI-KMT224 | OM912215 | amy-Wa2c | OM960453 |
| COI-LB01 | OM912216 | amy-fff2c | OM960454 |
| COI-LB02 | OM912217 | amy-CFY01a | OM960455 |
| COI-LB04 | OM912218 | amy-CFY02a | OM960456 |
| COI-LB05 | OM912219 | amy-CFY03a | OM960457 |
| COI-LB08 | OM912220 | amy-CFY04a | OM960458 |
| COI-LB11 | OM912221 | amy-CFY05a | OM960459 |
| COI-LB12 | OM912222 | amy-CFY07a | OM960460 |
| COI-LB13 | OM912223 | amy-CFY08a | OM960461 |
| COI-LB14 | OM912224 | amy-CFY09a | OM960462 |
| COI-LB16 | OM912225 | amy-CFY10a | OM960463 |
| COI-LB17 | OM912226 | amy-CFY11a | OM960464 |
| COI-LB18 | OM912227 | amy-CFY12a | OM960465 |
| COI-LB19 | OM912228 | amy-CFG14a | OM960466 |
| COI-LB20 | OM912229 | amy-CFG16a | OM960467 |
| COI-LB22 | OM912230 | amy-CFG17a | OM960468 |
| COI-LB31 | OM912231 | amy-CFG18a | OM960469 |
| COI-LB36 | OM912232 | amy-CFG19a | OM960470 |
| COI-LB40 | OM912233 | amy-CFG20a | OM960471 |
| COI-LB42 | OM912234 | amy-CFG21 | OM960472 |
| COI-LB43 | OM912235 | amy-CFG22a | OM960473 |
| COI-LB44 | OM912236 | amy-CFG23a | OM960474 |
| COI-LB45 | OM912237 | amy-Cl201 | OM960475 |
| COI-LB47 | OM912238 | amy-Cl204a | OM960476 |
| COI-LB48 | OM912239 | amy-Cl204c | OM960477 |
| COI-LB49 | OM912240 | amy-Cl205a | OM960478 |
| COI-LB58 | OM912241 | amy-Cl205c | OM960479 |
| COI-LB59 | OM912242 | amy-Cl206a | OM960480 |
| COI-LB62 | OM912243 | amy-Cl206c | OM960481 |
| COI-LB64 | OM912244 | amy-Cl207a | OM960482 |
| COI-LB66 | OM912245 | amy-Cl207c | OM960483 |
| COI-LB67 | OM912246 | amy-Cl208a | OM960484 |
| COI-LB68 | OM912247 | amy-Cl208c | OM960485 |
| COI-LB69 | OM912248 | amy-Cl210a | OM960486 |
| COI-LB71 | OM912249 | amy-Cl210c | OM960487 |
| COI-LB72 | OM912250 | amy-Cl211a | OM960488 |
| COI-LB74 | OM912251 | amy-Cl211c | OM960489 |

| | | | |
|---|---|---|---|
| COI-LB75 | OM912252 | amy-Cl212a | OM960490 |
| COI-LB76 | OM912253 | amy-Cl212b | OM960491 |
| COI-LB77 | OM912254 | amy-Cl213a | OM960492 |
| COI-LB80 | OM912255 | amy-Cl213c | OM960493 |
| COI-LB81 | OM912256 | amy-Cl214a | OM960494 |
| COI-LB91 | OM912257 | amy-Cl214c | OM960495 |
| COI-LB92 | OM912258 | amy-Cl215a | OM960496 |
| COI-LB98 | OM912259 | amy-Cl215c | OM960497 |
| COI-Mad2 | OM912260 | amy-Cl216a | OM960498 |
| COI-Sa22 | OM912261 | amy-Cl216c | OM960499 |
| COI-Sa23 | OM912262 | amy-Cl218a | OM960500 |
| COI-Sa24 | OM912263 | amy-Cl218c | OM960501 |
| COI-Sa25 | OM912264 | amy-Cl220a | OM960502 |
| COI-Sa26 | OM912265 | amy-Cl220c | OM960503 |
| COI-Sa72 | OM912266 | amy-Cl221a | OM960504 |
| COI-Sa73 | OM912267 | amy-Cl221c | OM960505 |
| COI-Sa74 | OM912268 | amy-Cl222a | OM960506 |
| COI-Sa75 | OM912269 | amy-Cl222c | OM960507 |
| COI-Su1 | OM912270 | amy-Cl223a | OM960508 |
| COI-Su2 | OM912271 | amy-Cl223c | OM960509 |
| COI-Su3 | OM912272 | amy-Cl224a | OM960510 |
| COI-Su4 | OM912273 | amy-Cl224c | OM960511 |
| COI-Su5 | OM912274 | amy-LB05 | OM960512 |
| COI-Vt01 | OM912275 | amy-LB11a | OM960513 |
| COI-Vt02 | OM912276 | amy-LB12b | OM960514 |
| COI-Vt03 | OM912277 | amy-LB17a | OM960515 |
| COI-Vt07 | OM912278 | amy-LB18a | OM960516 |
| COI-Vt10 | OM912279 | amy-LB20a | OM960517 |
| COI-Vt11 | OM912280 | amy-LB22a | OM960518 |
| COI-Vt12 | OM912281 | amy-LB36 | OM960519 |
| COI-Vt13 | OM912282 | amy-LB42a | OM960520 |
| COI-Vt15 | OM912283 | amy-LB49 | OM960521 |
| COI-Vt16 | OM912284 | amy-LB58 | OM960522 |
| COI-Vt17 | OM912285 | amy-LB67 | OM960523 |
| COI-Vt19 | OM912286 | amy-LB72a | OM960524 |
| COI-Vt20 | OM912287 | amy-LB81 | OM960525 |
| COI-yy12 | OM912288 | amy-LB01a | OM960526 |
| COI-ZA01 | OM912289 | amy-LB12a | OM960527 |
| COI-ZA03 | OM912290 | amy-LB01b | OM960528 |
| COI-ZA05 | OM912291 | amy-LB06b | OM960529 |
| COI-ZA06 | OM912292 | amy-LB69b | OM960530 |
| COI-ZA07 | OM912293 | amy-KMT205 | OM960531 |
| COI-ZA09 | OM912294 | amy-KMT208 | OM960532 |
| COI-ZA10 | OM912295 | amy-CJ01a | OM960533 |
| COI-ZA11 | OM912296 | amy-CJ03a | OM960534 |
| COI-ZA12 | OM912297 | amy-CJ04a | OM960535 |
| COI-ZA13 | OM912298 | amy-CJ11a | OM960536 |
| COI-ZA15 | OM912299 | amy-CJ12b | OM960537 |

| | | | |
|---|---|---|---|
| COI-ZA16 | OM912300 | amy-CJ01b | OM960538 |
| COI-ZA17 | OM912301 | amy-CJ05a | OM960539 |
| COI-ZA18 | OM912302 | amy-CJ10a | OM960540 |
| COI-ZA19 | OM912303 | amy-CJ12a | OM960541 |
| ATPS-103026 | OM959662 | amy-CJ19 | OM960542 |
| ATPS-266690 | OM959663 | amy-CJ03b | OM960543 |
| ATPS-266691a | OM959664 | amy-CJ05b | OM960544 |
| ATPS-AA12c | OM959665 | amy-CJ10b | OM960545 |
| ATPS-AA13c | OM959666 | amy-CJ15b | OM960546 |
| ATPS-Hw2b | OM959667 | amy-CJ18b | OM960547 |
| ATPS-Hw4b | OM959668 | amy-CJ21b | OM960548 |
| ATPS-xx11 | OM959669 | amy-CJ07a | OM960549 |
| ATPS-CFG14a | OM959670 | amy-CJ11b | OM960550 |
| ATPS-CFG16a | OM959671 | amy-CJ04b | OM960551 |
| ATPS-CFG17a | OM959672 | amy-CJ15a | OM960552 |
| ATPS-CFG18a | OM959673 | amy-CJ16a | OM960553 |
| ATPS-CFG19a | OM959674 | amy-CJ18a | OM960554 |
| ATPS-CFG20a | OM959675 | amy-CJ21a | OM960555 |
| ATPS-CFG21a | OM959676 | amy-CJ09a | OM960556 |
| ATPS-CFG22a | OM959677 | amy-CJ16b | OM960557 |
| ATPS-CFG23a | OM959678 | amy-CJ09b | OM960558 |
| ATPS-EHM101a | OM959679 | amy-CJ07b | OM960559 |
