## Supplementary material for "Substantial genetic mixing among sexual and androgenetic lineages within the clam genus *Corbicula*": Table S2

Table S2: list of the 359 analyzed individuals. The allele color code corresponds to the four invasive lineages: alleles retrieved from *C.* sp. form A/R, B, C/S and Rlc are respectively in red, blue, orange and green. Invasive alleles (IA), defined here as nuclear alleles found in at least two different invasive lineages, are shown in purple. Alleles shared between idiotypes (referring to Figure 4) are highlighted in bold, whereas individuals from Ref. 2 are followed by asterisks (*). For each marker, the methods used to retrieve each allele is indicated (d = direct sequencing, c = cloning, l = lite[illegible]

| Idiotype | Taxon | Location | Population code | Collection number | COI method | COI haplotypes | 28S methods | 28S haplotypes | α-amylase | α-amylase haplotypes | ATPS-α methods | ATPS-α haplotypes | Reproductive mode | Evidence of (a)sexuality |
|---|---|---|---|---|---|---|---|---|---|---|---|---|---|---|
| Form_AR | *C.* sp. form A/R (*C. fluminea* , white form) | San Gabriel River, USA | AA | AA10 | l | COI-FW5 | | | d | amy-IA1 amy-IA2 amy-AA10c | | | Androgenetic | Sperm morphology (1) |
| | | | | AA11 | l | COI-FW5 | d | 28S-IA1 28S-IA2 28S-IA3 | d+c | amy-IA1 amy-IA2 amy-AA10c | c | ATPS-AA11a ATPS-IA1 | | |
| | | | | AA12 | l | COI-FW5 | d+c | 28S-IA1 28S-IA2 28S-IA3 | d+c | amy-IA1 amy-IA2 amy-AA10c | c | ATPS-AA11a ATPS-IA1 ATPS-AA12c | | |
| | | | | AA13 | l | COI-FW5 | d | 28S-IA1 28S-IA2 28S-IA3 | d | amy-IA1 amy-IA2 amy-AA10c | c | ATPS-AA11a ATPS-IA1 ATPS-AA12c | | |
| | | | | AA14 | l | COI-FW5 | d | 28S-IA1 28S-IA2 28S-IA3 | d | amy-IA1 amy-IA2 amy-AA10c | | | | |
| | | Neuchâtel Lake, Switzerland | Su | Su1 | d | COI-FW5 | d | 28S-IA1 28S-IA2 28S-IA3 | d | amy-IA1 amy-IA2 amy-AA10c | | | Androgenetic | Sperm morphology (1) |
| | | | | Su2 | d | COI-FW5 | d | 28S-IA1 28S-IA2 28S-IA3 | d | amy-IA1 amy-IA2 amy-AA10c | | | | |
| | | | | Su3 | d | COI-FW5 | d | 28S-IA1 28S-IA2 28S-IA3 | d | amy-IA1 amy-IA2 amy-AA10c | | | | |
| | | | | Su4 | d | COI-FW5 | d | 28S-IA1 28S-IA2 28S-IA3 | d | amy-IA1 amy-IA2 amy-AA10c | | | | |
| | | | | Su5 | d | COI-FW5 | d | 28S-IA1 28S-IA2 28S-IA3 | d | amy-IA2 | | | | |
| | | Meuse River, Godinne, Belgium | Go | Go1 | | | | | | | c | ATPS-IA1 | Androgenetic | Lineage identification, histology |
| | | Pecos River, Texas, USA | ccc | ccc7* | | | c | 28S-IA3 28S-IA2 28S-IA1 | | | | | Androgenetic | Reproductive morphology, lack of genetic diversity (2) |
| | | USA | xx | xx11* | | | | | d | amy-IA1 amy-IA2 amy-AA10c | c | ATPS-AA12c | Androgenetic | Reproductive morphology, lack of genetic diversity (2) |
| | | Colorado River, Texas, USA | qq | qq1* | | | | | c | amy-qq1a amy-IA1 amy-IA2 | | | Androgenetic | Reproductive morphology, lack of genetic diversity (2) |
| | | The Netherlands | ggg | ggg2* | | | c | 28S-ggg2a 28S-IA1 28S-IA2 | c | amy-IA2 amy-AA10c | | | Androgenetic | Reproductive morphology, lack of genetic diversity (2) |
| Form_B | *C.* sp. form B (purple form) | San Gabriel River, USA | AB | AB11 | l | COI-FW1 | d | 28S-AB11a 28S-AB11b | d | amy-IA1 amy-IA2 amy-IA3 amy-AB12d | c | ATPS-AB11a ATPS-AB11b | Androgenetic | Sperm morphology (1) |
| | | | | AB12 | l | COI-FW1 | | | c | amy-IA1 amy-IA2 amy-IA3 amy-AB12d | d | ATPS-AB11a ATPS-AB11b | | |
| | | | | AB13 | l | COI-FW1 | d+c | 28S-AB11a 28S-AB11b | d+c | amy-IA1 amy-IA2 amy-IA3 amy-AB12d | c | ATPS-AB11a ATPS-AB11b | | |
| | | | | AB14 | l | COI-FW1 | d | 28S-AB11a 28S-AB11b | d | amy-IA1 amy-IA2 amy-IA3 amy-AB12d | d | ATPS-AB11a ATPS-AB11b | | |
| | | | | AB15 | l | COI-FW1 | d | 28S-AB11a 28S-AB11b | d | amy-IA1 amy-IA2 amy-IA3 amy-AB12d | c | ATPS-AB11a ATPS-AB11b | | |
| | | | | AB16 | l | COI-FW1 | d | 28S-AB11a 28S-AB11b | d | amy-IA1 amy-IA2 amy-IA3 amy-AB12d | | | | |
| | | USA | yy | yy11* | | | | | d | amy-IA3 amy-AB12d | | | Androgenetic | Reproductive morphology, lack of genetic diversity (2) |
| | | | | yy12* | d | COI-FW1 | c | 28S-AB11a 28S-AB11b 28S-yy12c | c | amy-IA3 amy-AB12d | c | ATPS-AB11a | | |
| | | Colorado River, Texas, USA | rr | rr1* | | | | | c | amy-rr1a amy-IA2 | | | Androgenetic | Reproductive morphology, lack of genetic diversity (2) |
| Form_CS | *C.* sp. form C/S (*C. fluminalis*, *C. largillierti* ) | Argentina | C | C1 | l | COI-FW17 | d+c | 28S-C1a 28S-IA1 28S-C1c | d | amy-C1a amy-C1b | d+c | ATPS-C1a ATPS-IA1 | Androgenetic | Lack of genetic diversity (1) |
| | | | | C2 | l | COI-FW17 | d | 28S-C1a 28S-IA1 28S-C1c | d | amy-C1a amy-C1b | d | ATPS-C1a ATPS-IA1 | | |
| | | | | C3 | l | COI-FW17 | d | 28S-C1a 28S-IA1 28S-C1c | d | amy-C1a amy-C1b | d | ATPS-C1a ATPS-IA1 | | |
| | | | | C4 | l | COI-FW17 | d | 28S-C1a 28S-IA1 28S-C1c | d+c | amy-C1a amy-C1b | d+c | ATPS-C1a ATPS-IA1 | | |
| | | Saône River, France | Sa | Sa22 | l | COI-FW17 | d | 28S-C1a 28S-IA1 28S-C1c | d | amy-C1a amy-C1b | d+c | ATPS-C1a ATPS-IA1 | | |
| | | | | Sa23 | l | COI-FW17 | d | 28S-C1a 28S-IA1 28S-C1c | d | amy-C1a amy-C1b | | | | |
| | | | | Sa24 | l | COI-FW17 | d | 28S-C1a 28S-IA1 28S-C1c | d | amy-C1a amy-C1b | | | | |
| | | | | Sa25 | l | COI-FW17 | d | 28S-C1a 28S-IA1 28S-C1c | d | amy-C1a amy-C1b | | | | |
| | | | | Sa26 | l | COI-FW17 | d | 28S-C1a 28S-IA1 28S-C1c | d | amy-C1a amy-C1b | | | | |
| | | | | Sa72 | l | COI-FW17 | | | | | d | ATPS-C1a ATPS-IA1 | | |
| | | | | Sa73 | l | COI-FW17 | | | | | d | ATPS-C1a ATPS-IA1 | | |
| | | | | Sa74 | l | COI-FW17 | | | | | d | ATPS-C1a ATPS-IA1 | | |
| | | | | Sa75 | l | COI-FW17 | | | | | d | ATPS-C1a ATPS-IA1 | | |
| | | Argentina | U | U3* | | | c | 28S-U3a 28S-IA1 28S-C1a | | | c | ATPS-U3a ATPS-C1a | Androgenetic | Sperm morphology (2) |
| | | | | U4* | | | | | c | amy-C1b amy-U4b | | | | |
| | | The Netherlands | S | ff2* | | | | | c | amy-C1b amy-C1a amy-fff2c | c | ATPS-C1a | Androgenetic | Lineage identification (2) |
| Form_Rlc | *C.* sp. form Rlc | Doubs River, France | Rlc | Db22 | d | COI-FW4 | d | 28S-IA2 | d | amy-Db22a amy-IA3 | d | ATPS-Db22a ATPS-Db22b | Androgenetic | Sperm morphology (1) |
| | | | | Db23 | d | COI-FW4 | d | 28S-IA2 | d | amy-Db22a amy-IA3 | d | ATPS-Db22a ATPS-Db22b | | |
| | | | | Db24 | d | COI-FW4 | d | 28S-IA2 28S-IA3 | d+c | amy-Db22a amy-IA3 | d | ATPS-Db22a ATPS-Db22b | | |
| | | | | Db25 | d | COI-FW5 | d+c | 28S-IA2 | d+c | amy-Db22a amy-IA3 | d+c | ATPS-Db22a ATPS-Db22b | | |
| | | | | Db26 | d | COI-FW4 | d | 28S-IA2 | d | amy-Db22a amy-IA3 | d | ATPS-Db22a ATPS-Db22b | | |
| Waal | *C.* sp. | Waal River, The Netherlands | Wa | Wa1 | | | c | 28S-IA1 28S-C1c 28S-C1a 28S-IA3 | c | amy-C1b | c | ATPS-C1a ATPS-IA1 | Androgenetic | Lack of genetic diversity |
| | | | | Wa2 | | | c | 28S-IA1 28S-C1c 28S-C1a | c | amy-C1a amy-C1b amy-fff2c | c | ATPS-C1a ATPS-IA1 | | |
| Hawaii | *C.* sp. | Hawaii | Hw | Hw2 | | | d+c | 28S-IA3 28S-IA2 28S-IA1 | | | c | ATPS-AA11a ATPS-AA12c ATPS-IA1 | ? | |
| | | | | Hw3 | | | d | 28S-IA3 28S-IA2 28S-IA1 | | | | | | |
| | | | | Hw4 | | | d+c | **28S-KMT205a** **28S-Hw4b** 28S-IA2 28S-IA1 | | | c | ATPS-AA11a ATPS-AA12c ATPS-IA1 ATPS-AB11a | | |
| | | | | Hw5 | | | d | **28S-KMT205a** **28S-Hw4b** 28S-IA2 28S-IA1 | | | | | | |
| | | | | Hw6 | | | d | **28S-KMT205a** **28S-Hw4b** 28S-IA2 28S-IA1 | | | | | | |
| China | *C.* sp. (*C. largillierti* ) | Baxin, Yunnan, China | BA | BA1 | d | COI-BA1 | | | | | | | ? | |
| | | | | BA2 | | | d | 28S-AB11a **28S-KMT205a** | | | | | | |
| | | | | BA3 | | | d | 28S-AB11a **28S-KMT205a** | | | | | | |
| | | | | BA4 | | | d | **28S-KMT205a** 28S-BA4b | | | | | | |
| | | | | BA5 | | | d+c | 28S-yy12c 28S-IA3 **28S-KMT205a** | | | d+c | ATPS-AB11b ATPS-FU4a | | |
| | | | | BA6 | d | COI-BA1 | d | 28S-yy12c 28S-IA3 | | | | | | |
| | | | | BA7 | | | d | 28S-yy12c 28S-IA3 | | | | | | |
| | | | | BA8 | | | d+c | 28S-yy12c 28S-IA3 28S-BA8c | | | c | ATPS-FU4a | | |
| | | Mujiang River, Yunnan, China | Muj | Muj02 | | | d+c | 28S-ggg2a 28S-IA2 28S-IA1 | | | | | ? | |
| | | | | Muj08 | | | d+c | 28S-AB11a 28S-FU4a 28S-Muj08c 28S-yy12c | | | | | | |
| | | | | Muj10 | | | d+c | 28S-AB11a 28S-FU4a 28S-Muj08c 28S-yy12c | | | | | | |
| | *C.* sp. | Fuxian Hu Lake, Yunnan, China | FU | FU1 | d | COI-BA1 | | | | | | | ? | |
| | | | | FU4 | | | d+c | 28S-FU4a 28S-FU4b 28S-yy12c | | | c | ATPS-FU4a ATPS-FU4b ATPS-IA1 | | |
| | | | | FU5 | | | d | 28S-FU4a 28S-FU4b 28S-yy12c | | | d | ATPS-FU4a ATPS-FU4b ATPS-IA1 | | |
| | | | | FU6 | | | d+c | 28S-FU4a 28S-FU4b 28S-yy12c | | | d+c | ATPS-FU4a ATPS-FU4b ATPS-IA1 | | |
| C_japonica | *C. japonica* | Kyoto, Japan | Jp | Jp01 | d | COI-Jp1 | d | 28S-CJ01a 28S-CJ01b | | | | | Sexual | Lineage identification (1), cytological evidence (3) |
| | | | | Jp02 | d | COI-Jp2 | d | 28S-CJ01a 28S-CJ01b | | | | | | |
| | | | | Jp03 | d | COI-Jp3 | | | | | d | ATPS-CJ02a ATPS-CJ02b | | |
| | | | | Jp04 | | | | | | | d | ATPS-CJ02b | | |
| | | | | Jp06 | | | | | | | d | ATPS-Jp06a ATPS-CJ02b | | |
| | | | | Jp08 | | | | | | | d | ATPS-CJ02a ATPS-CJ02b | | |
| | | | | Jp10 | d | COI-Jp10 | d | 28S-CJ01b | | | d | ATPS-CJ02a ATPS-CJ02b | | |
| | | Kamo River, Kyoto, Japan | KA | Ka1 | d | COI-Ka1 | | | | | | | | |
| | | | | Ka2 | d | COI-Ka2 | | | | | | | | |
| | | | | Ka3 | d | COI-Ka2 | | | | | d | ATPS-Ka3a ATPS-Ka3b | | |
| | | | | Ka5 | d | COI-Ka5 | | | | | | | | |
| | | Japan | CJ | CJ01 | d | COI-CJ1 | d | 28S-CJ01a 28S-CJ01b | d | amy-CJ01a amy-CJ01b | | | | |
| | | | | CJ02 | | | d | 28S-CJ01b | | | d | ATPS-CJ02a ATPS-CJ02b | | |
| | | | | CJ03 | d | COI-Jp3 | d | 28S-CJ01b | d | amy-CJ03a amy-CJ03b | d | ATPS-CJ02a ATPS-CJ03b | | |
| | | | | CJ04 | d | COI-Jp3 | d | 28S-CJ01b | d | amy-CJ01a amy-CJ04b | d | ATPS-CJ02a ATPS-CJ02b | | |
| | | | | CJ05 | d | COI-Jp3 | d | 28S-CJ01a 28S-CJ01b | d | amy-CJ05a amy-CJ03b | | | | |
| | | | | CJ06 | | | d | 28S-CJ01a 28S-CJ01b | | | d | ATPS-CJ02a ATPS-CJ02b | | |
| | | | | CJ07 | d | COI-CJ7 | d | 28S-CJ01a 28S-CJ01b | d | amy-CJ07a amy-CJ07b | d | ATPS-CJ03b ATPS-CJ07b | | |
| | | | | CJ08 | | | d | 28S-CJ01a 28S-CJ01b | | | | | | |
| | | | | CJ09 | d | COI-CJ1 | d | 28S-CJ01a 28S-CJ01b | d | amy-CJ09a amy-CJ09b | d | ATPS-CJ02a ATPS-CJ02b | | |
| | | | | CJ10 | d | COI-CJ1 | d | 28S-CJ01a 28S-CJ01b | d | amy-CJ05a amy-CJ10b | d | ATPS-CJ02a | | |
| | | | | CJ11 | d | COI-CJ1 | d | 28S-CJ01a 28S-CJ01b | d | amy-CJ01a amy-CJ07a | d | ATPS-CJ02a ATPS-CJ02b | | |
| | | | | CJ12 | d | COI-CJ12 | d | 28S-CJ01a 28S-CJ01b | d | amy-CJ12a amy-CJ01a | d | ATPS-CJ03b ATPS-CJ12b | | |
| | | | | CJ13 | | | d | 28S-CJ01a 28S-CJ01b | | | d | ATPS-CJ02a | | |
| | | | | CJ14 | | | d | 28S-CJ01a 28S-CJ01b | | | d | ATPS-CJ12b ATPS-CJ14b | | |
| | | | | CJ15 | d | COI-Jp3 | d | 28S-CJ01a 28S-CJ01b | d | amy-CJ04b amy-CJ03b | d | ATPS-CJ03b ATPS-CJ02b | | |
| | | | | CJ16 | | | d | 28S-CJ01a 28S-CJ01b | d | amy-CJ04b amy-CJ09a | | | | |
| | | | | CJ17 | | | d | 28S-CJ17a 28S-CJ01b | | | | | | |
| | | | | CJ18 | d | COI-Jp3 | d | 28S-CJ01b | d | amy-CJ04b amy-CJ03b | d | ATPS-CJ18a ATPS-CJ18b | | |
| | | | | CJ19 | d | COI-CJ19 | d | 28S-CJ01a 28S-CJ01b | d | amy-CJ01b | d | ATPS-CJ02a | | |
| | | | | CJ20 | | | d | 28S-CJ01a 28S-CJ01b | | | d | ATPS-CJ03b | | |
| | | | | CJ21 | d | COI-Jp3 | | | d | amy-CJ04b amy-CJ03b | | | | |
| | | | | CJ22 | | | d | 28S-CJ01b | | | d | ATPS-CJ02a ATPS-CJ03b | | |
| South-East_Asia | *C.* sp. (*C. fluminea* ) | Korea | UMMZ | 266690* | | | | | | | c | ATPS-AA12c | Androgenetic | Lack of genetic diversity (2) |
| | | Taiwan | ZMB | 170096* | d | COI-170096 | | | c | amy-IA2 | | | Androgenetic | Unreduced sperm (2) |
| | | Kalasin, Lampao Dam, Thailand | UMMZ | 266691* | | | | | | | c | ATPS-AA12c **ATPS-CFG01b** | Androgenetic | Lineage identification (2) |
| | | Philippines | ZMB | 103026* | | | | | | | c | ATPS-AA12c | Androgenetic | Lineage identification (2) |
| Japan | *C.* sp. (*C. fluminea* /*leana* morphs) | Ehime Shigenobu Takubo, Japan | EHM | EHM101 | | | | | | | d | ATPS-CFG14a ATPS-Db22a | Androgenetic | Sperm morphology (1) |
| | | | | EHM102 | d | COI-FW1 | d | 28S-AB11a | d | amy-EHM102a amy-AB12d | d | ATPS-CFG14a ATPS-Db22a | | |
| | | | | EHM103 | d | COI-FW1 | d+c | 28S-AB11a | d+c | amy-EHM102a amy-AB12d | d+c | ATPS-CFG14a ATPS-Db22a | | |
| | | | | EHM104 | d | COI-FW1 | d | 28S-AB11a | d | amy-EHM102a amy-AB12d | d | ATPS-CFG14a ATPS-Db22a | | |
| | | | | EHM105 | d | COI-FW1 | d | 28S-AB11a | d | amy-EHM102a amy-AB12d | d | ATPS-CFG14a ATPS-Db22a | | |
| | | | | EHM106 | d | COI-FW1 | d | 28S-AB11a | d | amy-EHM102a amy-AB12d | d | ATPS-CFG14a | | |
| | | | | EHM107 | d | COI-FW1 | d | 28S-AB11a | d | amy-EHM102a amy-AB12d | d | ATPS-CFG14a ATPS-Db22a | | |
| | | | | EHM108 | d | COI-FW1 | d | 28S-AB11a | d | amy-EHM102a amy-AB12d | d | ATPS-CFG14a ATPS-Db22a | | |
| | | | | EHM109 | d | COI-FW1 | d+c | 28S-AB11a | d+c | amy-EHM102a amy-AB12d | d+c | ATPS-CFG14a ATPS-Db22a | | |
| | | | | EHM110 | d | COI-FW1 | d | 28S-AB11a | d | amy-EHM102a amy-AB12d | d | ATPS-CFG14a ATPS-Db22a | | |
| | | | | EHM111 | d | COI-FW1 | d | 28S-AB11a | d | amy-EHM102a amy-AB12d | d+c | ATPS-CFG14a ATPS-Db22a | | |
| | | | | EHM112 | d | COI-FW1 | d | 28S-AB11a | d | amy-EHM102a amy-AB12d | d | ATPS-CFG14a ATPS-Db22a | | |
| | | | | EHM113 | d | COI-FW1 | d | 28S-AB11a | d | amy-EHM102a amy-AB12d | d | ATPS-CFG14a ATPS-Db22a | | |
| | | | | EHM114 | | | | | | | d | ATPS-CFG14a ATPS-Db22a | | |
| | | | | EHM115 | | | | | | | d | ATPS-CFG14a | | |
| | | | | EHM116 | | | | | | | d | ATPS-CFG14a ATPS-Db22a | | |
| | | | | EHM119 | d | COI-FW1 | d | 28S-AB11a | d | amy-EHM102a amy-AB12d | d | ATPS-CFG14a ATPS-Db22a | | |
| | | | | EHM120 | d | COI-FW1 | d | 28S-AB11a | d | amy-EHM102a amy-AB12d | d | ATPS-CFG14a ATPS-Db22a | | |
| | | | | EHM121 | d | COI-FW1 | d | 28S-AB11a | d | amy-EHM102a amy-AB12d | d | ATPS-CFG14a ATPS-Db22a | | |
| | | | | EHM122 | d | COI-FW1 | d | 28S-AB11a | d | amy-EHM102a amy-AB12d | d+c | ATPS-CFG14a ATPS-Db22a | | |
| | | | | EHM123 | d | COI-FW1 | d | 28S-AB11a | d | amy-EHM102a amy-AB12d | | | | |
| | | | | EHM124 | d | COI-FW1 | d | 28S-AB11a | d | amy-EHM102a amy-AB12d | | | | |
| | | Miyazaki Gongenbaru Canal, Japan | KMT | KMT201 | | | d | 28S-AB11a **28S-KMT205a** 28S-AB11b | | | d | ATPS-Db22b **ATPS-KMT205b** | Androgenetic | Lineage identification |
| | | | | KMT202 | | | d | 28S-AB11a **28S-KMT205a** 28S-AB11b | | | d | ATPS-Db22b **ATPS-KMT205b** | | |
| | | | | KMT203 | | | d | 28S-AB11a **28S-KMT205a** 28S-AB11b | | | d | ATPS-Db22b **ATPS-KMT205b** | | |
| | | | | KMT204 | | | d | 28S-AB11a **28S-KMT205a** 28S-AB11b | | | d | ATPS-Db22b **ATPS-KMT205b** | | |
| | | | | KMT205 | d | COI-FW5 | d+c | 28S-AB11a **28S-KMT205a** 28S-AB11b | d | amy-KMT205 | d+c | ATPS-Db22b **ATPS-KMT205b** | | |
| | | | | KMT206 | | | d | 28S-AB11a **28S-KMT205a** 28S-AB11b | | | d | ATPS-Db22b **ATPS-KMT205b** | | |
| | | | | KMT207 | | | d | 28S-AB11a **28S-KMT205a** 28S-AB11b | | | d | ATPS-Db22b **ATPS-KMT205b** | | |
| | | | | KMT208 | d | COI-FW5 | d | 28S-AB11a **28S-KMT205a** 28S-AB11b | d | amy-KMT205 | d | ATPS-Db22b **ATPS-KMT205b** | | |
| | | | | KMT209 | | | d | 28S-AB11a **28S-KMT205a** 28S-AB11b | | | d | ATPS-Db22b **ATPS-KMT205b** | | |
| | | | | KMT210 | | | d | 28S-AB11a **28S-KMT205a** 28S-AB11b | | | d | ATPS-Db22b **ATPS-KMT205b** | | |
| | | | | KMT211 | | | d | 28S-AB11a 28S-yy12c | d+c | amy-IA2 amy-Db22a | d | **ATPS-KMT205** **ATPS-CFG01b** ATPS-AB11a | | |
| | | | | KMT212 | | | d | 28S-AB11a 28S-yy12c | d | amy-IA2 amy-Db22a | d | **ATPS-KMT205** **ATPS-CFG01b** ATPS-AB11a | | |
| | | | | KMT213 | d | COI-FW5 | d | 28S-AB11a 28S-yy12c | d | amy-IA2 amy-Db22a | d | **ATPS-KMT205** **ATPS-CFG01b** ATPS-AB11a | | |
| | | | | KMT214 | d | COI-FW5 | d | 28S-AB11a 28S-yy12c | d | amy-IA2 amy-Db22a | d | **ATPS-KMT205** **ATPS-CFG01b** ATPS-AB11a | | |
| | | | | KMT215 | d | COI-FW5 | d | 28S-AB11a 28S-yy12c | d | amy-IA2 amy-Db22a | d | **ATPS-KMT205** **ATPS-CFG01b** ATPS-AB11a | | |
| | | | | KMT216 | d | COI-FW5 | d | 28S-AB11a 28S-yy12c | d | amy-IA2 amy-Db22a | d | **ATPS-KMT205** **ATPS-CFG01b** ATPS-AB11a | | |
| | | | | KMT217 | d | COI-FW5 | d+c | 28S-AB11a 28S-yy12c | d+c | amy-IA2 amy-Db22a | d+c | **ATPS-KMT205** **ATPS-CFG01b** ATPS-AB11a | | |
| | | | | KMT218 | d | COI-FW5 | d | 28S-AB11a 28S-yy12c | d | amy-IA2 amy-Db22a | d | **ATPS-KMT205** **ATPS-CFG01b** ATPS-AB11a | | |
| | | | | KMT219 | d | COI-FW5 | d | 28S-AB11a 28S-yy12c | d | amy-IA2 amy-Db22a | d | **ATPS-KMT205** **ATPS-CFG01b** ATPS-AB11a | | |
| | | | | KMT220 | | | | | | | d | **ATPS-KMT205** **ATPS-CFG01b** ATPS-AB11a | | |
| | | | | KMT221 | d | COI-FW5 | d | 28S-AB11a 28S-yy12c | d | amy-IA2 amy-Db22a | d | **ATPS-KMT205** **ATPS-CFG01b** ATPS-AB11a | | |
| | | | | KMT222 | d | COI-FW5 | d | 28S-AB11a 28S-yy12c | d | amy-IA2 amy-Db22a | d | **ATPS-KMT205** **ATPS-CFG01b** ATPS-AB11a | | |
| | | | | KMT223 | | | | | | | d | **ATPS-KMT205** **ATPS-CFG01b** ATPS-AB11a | | |
| | | | | KMT224 | d | COI-FW5 | d | 28S-AB11a 28S-yy12c | d | amy-IA2 amy-Db22a | d | **ATPS-KMT205** **ATPS-CFG01b** ATPS-AB11a | | |
| | | Ehime, Japan | CI | CI201 | | | d+c | 28S-CI201a 28S-IA1 28S-CI201c | d+c | **amy-CFG14a** | | | Androgenetic | Lineage identification |
| | | | | CI202 | | | d | 28S-CI201a 28S-IA1 28S-CI201c | | | | | | |
| | | | | CI204 | d | COI-CI202 | d | 28S-IA3 28S-CI204b **28S-CI204c** | d | **amy-CFG14a** amy-Db22a amy-CI204c | d | ATPS-AB11a | | |
| | | | | CI205 | d | COI-CI202 | d | 28S-IA3 28S-CI204b **28S-CI204c** | d | **amy-CFG14a** amy-Db22a amy-CI204c | d | ATPS-AB11a | | |
| | | | | CI206 | d | COI-CI202 | d | 28S-IA3 28S-CI204b **28S-CI204c** | d | **amy-CFG14a** amy-Db22a amy-CI204c | d | ATPS-AB11a | | |
| | | | | CI207 | d | COI-CI202 | d | 28S-IA3 28S-CI204b **28S-CI204c** | d | **amy-CFG14a** amy-Db22a amy-CI204c | d | ATPS-AB11a | | |
| | | | | CI208 | d | COI-CI202 | d | 28S-IA3 28S-CI204b **28S-CI204c** | d | **amy-CFG14a** amy-Db22a amy-CI204c | d | ATPS-AB11a | | |
| | | | | CI209 | | | | | | | d | ATPS-AB11a | | |
| | | | | CI210 | d | COI-CI202 | d | 28S-IA3 28S-CI204b **28S-CI204c** | d | **amy-CFG14a** amy-Db22a amy-CI204c | d | ATPS-AB11a | | |
| | | | | CI211 | d | COI-CI202 | d | 28S-IA3 28S-CI204b **28S-CI204c** | d | **amy-CFG14a** amy-Db22a amy-CI204c | d | ATPS-AB11a | | |
| | | | | CI212 | d | COI-CI202 | d | 28S-IA3 28S-CI204b **28S-CI204c** | c | **amy-CFG14a** amy-CI204c | d | ATPS-AB11a | | |
| | | | | CI213 | d | COI-CI202 | d | 28S-IA3 28S-CI204b **28S-CI204c** | d | **amy-CFG14a** amy-Db22a amy-CI204c | d | ATPS-AB11a | | |
| | | | | CI214 | d | COI-CI202 | d | 28S-IA3 28S-CI204b **28S-CI204c** | d | **amy-CFG14a** amy-Db22a amy-CI204c | d | ATPS-AB11a | | |
| | | | | CI215 | d | COI-CI202 | d | 28S-IA3 28S-CI204b **28S-CI204c** | d | **amy-CFG14a** amy-Db22a amy-CI204c | d | ATPS-AB11a | | |
| | | | | CI216 | d | COI-CI202 | d | 28S-IA3 28S-CI204b **28S-CI204c** | d | **amy-CFG14a** amy-Db22a amy-CI204c | d | ATPS-AB11a | | |
| | | | | CI217 | | | | | | | d | ATPS-AB11a | | |
| | | | | CI218 | d | COI-CI202 | d | 28S-IA3 28S-CI204b **28S-CI204c** | d | **amy-CFG14a** amy-Db22a amy-CI204c | d | ATPS-AB11a | | |
| | | | | CI220 | d | COI-CI202 | d | 28S-IA3 28S-CI204b **28S-CI204c** | d | **amy-CFG14a** amy-Db22a amy-CI204c | d | ATPS-AB11a | | |
| | | | | CI221 | d | COI-CI202 | d | 28S-IA3 28S-CI204b **28S-CI204c** | d | **amy-CFG14a** amy-Db22a amy-CI204c | d | ATPS-AB11a | | |
| | | | | CI222 | d | COI-CI202 | d | 28S-IA3 28S-CI204b **28S-CI204c** | d | **amy-CFG14a** amy-Db22a amy-CI204c | d | ATPS-AB11a | | |
| | | | | CI223 | d | COI-CI202 | d | 28S-IA3 28S-CI204b **28S-CI204c** | d | **amy-CFG14a** amy-Db22a amy-CI204c | d | ATPS-AB11a | | |
| | | | | CI224 | d | COI-CI202 | d | 28S-IA3 28S-CI204b **28S-CI204c** | d | **amy-CFG14a** amy-Db22a amy-CI204c | d | ATPS-AB11a | | |
| | *C.* sp. (*C. fluminea* yellow morph) | Sakanai River, Matsusaka, Japan | CFY | CFY01 | | | d+c | 28S-IA2 28S-IA1 | d+c | amy-CFY01a amy-Db22a | d+c | ATPS-AB11a | Androgenetic | Lineage identification |
| | | | | CFY02 | | | d | 28S-IA2 28S-IA1 | d | amy-CFY01a amy-Db22a | d | ATPS-AB11a | | |
| | | | | CFY03 | d | COI-FW5 | d | 28S-IA2 28S-IA1 | d | amy-CFY01a amy-Db22a | d | ATPS-AB11a | | |
| | | | | CFY04 | d | COI-FW5 | d | 28S-IA2 28S-IA1 | d | amy-CFY01a amy-Db22a | d | ATPS-AB11a | | |
| | | | | CFY05 | d | COI-FW5 | d | 28S-IA2 28S-IA1 | d | amy-CFY01a amy-Db22a | d | ATPS-AB11a | | |
| | | | | CFY06 | | | d+c | 28S-CI204b 28S-ggg2a 28S-AB11b | d+c | amy-IA1 amy-Db22a | d+c | ATPS-Db22a **ATPS-CFG01b** | | |
| | | | | CFY07 | d | COI-CI202 | d | 28S-IA2 28S-IA1 | d | amy-CFY01a amy-Db22a | d | ATPS-AB11a | | |
| | | | | CFY08 | | | d | 28S-IA2 28S-IA1 | d | amy-CFY01a amy-Db22a | d | ATPS-AB11a | | |
| | | | | CFY09 | d | COI-FW5 | d | 28S-IA2 28S-IA1 | d | amy-CFY01a amy-Db22a | d | ATPS-AB11a | | |
| | | | | CFY10 | d | COI-FW5 | d | 28S-IA2 28S-IA1 | d | amy-CFY01a amy-Db22a | d | ATPS-AB11a | | |
| | | | | CFY11 | d | COI-FW5 | d | 28S-IA2 28S-IA1 | d | amy-CFY01a amy-Db22a | d | ATPS-AB11a | | |
| | | | | CFY12 | d | COI-FW5 | d | 28S-IA2 28S-IA1 | d | amy-CFY01a amy-Db22a | d | ATPS-AB11a | | |
| | *C.* sp. (*C. fluminea* green morph) | Tokida River, Watarai, Japan | CFG | CFG01 | d | COI-FW5 | d+c | 28S-CI204b 28S-ggg2a | d+c | amy-IA1 amy-Db22a | d+c | ATPS-Db22a **ATPS-CFG01b** | Androgenetic | Lineage identification |
| | | | | CFG02 | d | COI-FW5 | d | 28S-CI204b 28S-ggg2a | d | amy-IA1 amy-Db22a | d | ATPS-Db22a **ATPS-CFG01b** | | |
| | | | | CFG03 | d | COI-FW5 | d | 28S-CI204b 28S-ggg2a | d | amy-IA1 amy-Db22a | d | ATPS-Db22a **ATPS-CFG01b** | | |
| | | | | CFG04 | d | COI-FW5 | d | 28S-CI204b 28S-ggg2a | d | amy-IA1 amy-Db22a | d | ATPS-Db22a **ATPS-CFG01b** | | |
| | | | | CFG05 | d | COI-FW5 | d | 28S-CI204b 28S-ggg2a | d | amy-IA1 amy-Db22a | d | ATPS-Db22a **ATPS-CFG01b** | | |
| | | | | CFG06 | d | COI-FW5 | d | 28S-CI204b 28S-ggg2a | d | amy-IA1 amy-Db22a | d | ATPS-Db22a **ATPS-CFG01b** | | |
| | | | | CFG07 | d | COI-FW5 | d | 28S-CI204b 28S-ggg2a | d | amy-IA1 amy-Db22a | d | ATPS-Db22a **ATPS-CFG01b** | | |
| | | | | CFG08 | d | COI-FW5 | d | 28S-CI204b 28S-ggg2a | d | amy-IA1 amy-Db22a | d | ATPS-Db22a **ATPS-CFG01b** | | |
| | | | | CFG09 | d | COI-FW5 | d | 28S-CI204b 28S-ggg2a | d | amy-IA1 amy-Db22a | d | ATPS-Db22a **ATPS-CFG01b** | | |
| | | | | CFG10 | d | COI-FW5 | d | 28S-CI204b 28S-ggg2a | d | amy-IA1 amy-Db22a | d | ATPS-Db22a **ATPS-CFG01b** | | |
| | | | | CFG11 | d | COI-FW5 | d | 28S-CI204b 28S-ggg2a | d | amy-IA1 amy-Db22a | d | ATPS-Db22a **ATPS-CFG01b** | | |
| | | | | CFG12 | d | COI-FW5 | d | 28S-CI204b 28S-ggg2a | d | amy-IA1 amy-Db22a | d | ATPS-Db22a **ATPS-CFG01b** | | |
| | | | | CFG13 | d | COI-FW5 | d | 28S-CI204b 28S-ggg2a | d | amy-IA1 amy-Db22a | d | ATPS-Db22a **ATPS-CFG01b** | | |
| | | Kushida River, Matsusaka, Japan | CFG | CFG14 | | | d | 28S-AB11a | d | **amy-CFG14a** amy-Db22a | d | ATPS-CFG14a **ATPS-CFG14b** | Androgenetic | Lineage identification |
| | | | | CFG15 | d | COI-FW5 | d | 28S-CI204b 28S-ggg2a | d | amy-IA1 amy-Db22a | d | ATPS-Db22a **ATPS-CFG01b** | | |
| | | | | CFG16 | d | COI-FW1 | d+c | 28S-AB11a 28S-IA2 | d+c | **amy-CFG14a** amy-Db22a | d+c | ATPS-CFG14a **ATPS-CFG14b** | | |
| | | | | CFG17 | d | COI-FW1 | d | 28S-AB11a | d | **amy-CFG14a** amy-Db22a | d | ATPS-CFG14a **ATPS-CFG14b** | | |
| | | | | CFG18 | d | COI-FW1 | d | 28S-AB11a | d | **amy-CFG14a** amy-Db22a | d | ATPS-CFG14a **ATPS-CFG14b** | | |
| | | | | CFG19 | d | COI-FW1 | d | 28S-AB11a 28S-IA2 | d | **amy-CFG14a** amy-Db22a | d | ATPS-CFG14a **ATPS-CFG14b** | | |
| | | | | CFG20 | d | COI-FW1 | d | 28S-AB11a | d | **amy-CFG14a** amy-Db22a | d | ATPS-CFG14a **ATPS-CFG14b** | | |
| | | | | CFG21 | d | COI-FW1 | d | 28S-AB11a | d | **amy-CFG14a** | d | ATPS-CFG14a **ATPS-CFG14b** | | |
| | | | | CFG22 | d | COI-FW1 | d | 28S-AB11a 28S-IA2 | d | **amy-CFG14a** amy-Db22a | d | ATPS-CFG14a **ATPS-CFG14b** | | |
| | | | | CFG23 | d | COI-FW1 | d | 28S-AB11a 28S-IA2 | d | **amy-CFG14a** amy-Db22a | d | ATPS-CFG14a **ATPS-CFG14b** | | |
| Lake_Biwa | *C.* spp. (*C. fluminea* / *leana* / *sandai* morphs) | Seta River-Lake Biwa, Japan | LB | LB01 | d | COI-LB01 | | | d | amy-LB01a amy-LB01b | | | Sexual/Androgenetic | Lineage identification |
| | | | | LB02 | d | COI-FW4 | | | d | amy-IA1 amy-CS14b | | | | |
| | | | | LB04 | d | COI-FW1 | | | d | amy-LB04a amy-LB04b | d | ATPS-LB04a ATPS-LB04b | | |
| | | | | LB05 | d | COI-LB05 | | | d | **amy-CFG14a** | | | | |
| | | | | LB06 | d | | | | d | amy-IA1 amy-LB06b | | | | |
| | | | | LB08 | d | COI-FW4 | | | | | d | **ATPS-CFG01b** | | |
| | | | | LB11 | d | COI-LB11 | | | d+c | **amy-CFG14a** amy-IA2 | | | | |
| | | | | LB12 | d | COI-LB11 | | | d | amy-LB01a **amy-CFG14a** | d | ATPS-LB12 | | |
| | | | | LB13 | d | COI-FW5 | | | d | amy-IA1 amy-CS04a | d | **ATPS-CFG01b** | | |
| | | | | LB14 | d | COI-LB11 | | | d | amy-LB14 | | | | |
| | | | | LB16 | d | COI-FW4 | | | d | amy-IA1 amy-CS04a | d | **ATPS-CFG01b** | | |
| | | | | LB17 | d | COI-FW5 | | | d+c | **amy-CFG14a** amy-LB17b amy-IA1 | | | | |
| | | | | LB18 | d | COI-FW1 | | | d | **amy-CFG14a** amy-LB14 | | | | |
| | | | | LB19 | d | COI-FW5 | | | d | amy-IA1 amy-LB19b | d | **ATPS-CFG01b** | | |
| | | | | LB20 | d | COI-LB11 | | | d+c | **amy-CFG14a** amy-LB20b | d | ATPS-LB04b | | |
| | | | | LB22 | d | COI-LB22 | | | d+c | **amy-CFG14a** amy-LB14 | d | ATPS-LB22a ATPS-LB04a | | |
| | | | | LB31 | d | COI-FW5 | | | d | amy-IA1 | d | **ATPS-CFG01b** | | |
| | | | | LB36 | d | COI-CS04 | | | d | **amy-CFG14a** | d | **ATPS-CFG14b** | | |
| | | | | LB40 | d | COI-FW5 | | | d | amy-IA1 amy-CS04a | d | **ATPS-CFG01b** | | |
| | | | | LB42 | d | COI-LB01 | | | d | **amy-CFG14a** amy-LB14 | | | | |
| | | | | LB43 | d | COI-FW5 | | | d | amy-IA1 amy-CS14b | d | **ATPS-CFG01b** | | |
| | | | | LB44 | d | COI-FW4 | | | d | amy-IA1 amy-LB44b | d | **ATPS-CFG01b** | | |
| | | | | LB45 | d | COI-FW1 | | | | | | | | |
| | | | | LB47 | d | COI-FW4 | | | | | d | **ATPS-CFG01b** | | |
| | | | | LB48 | d | COI-FW1 | | | | | d | ATPS-LB04b | | |
| | | | | LB49 | d | COI-FW1 | | | d | **amy-CFG14a** | | | | |
| | | | | LB58 | d | COI-LB11 | | | d | **amy-CFG14a** | d | ATPS-LB04a | | |
| | | | | LB59 | d | COI-LB59 | | | d+c | amy-LB04a amy-LB04b | d | ATPS-LB04b | | |
| | | | | LB62 | d | COI-LB05 | | | d | amy-IA1 amy-LB62b | d | ATPS-LB12 | | |
| | | | | LB64 | d | COI-LB64 | | | d+c | amy-LB04a amy-IA2 amy-LB04b | d | ATPS-LB04b | | |
| | | | | LB66 | d | COI-FW5 | | | d | amy-IA1 amy-LB66b | d | **ATPS-CFG01b** | | |
| | | | | LB67 | d | COI-LB11 | | | d | **amy-CFG14a** | d | ATPS-LB04b | | |
| | | | | LB68 | d | COI-FW5 | | | d | amy-CS06b | d | **ATPS-CFG01b** | | |
| | | | | LB69 | d | COI-FW5 | | | d | amy-IA1 amy-LB69b | d | **ATPS-KMT201b** | | |
| | | | | LB71 | d | COI-LB11 | | | d | amy-IA1 amy-LB71b | | | | |
| | | | | LB72 | d | COI-FW5 | | | d+c | **amy-CFG14a** amy-CS06b | d | **ATPS-KMT201b** | | |
| | | | | LB74 | d | COI-LB22 | | | | | d | ATPS-LB22a ATPS-LB74b | | |
| | | | | LB75 | d | COI-CS04 | | | | | | | | |
| | | | | LB76 | d | COI-FW5 | | | d | amy-IA1 amy-CS04a | d | **ATPS-CFG01b** | | |
| | | | | LB77 | d | COI-LB77 | | | | | d | **ATPS-CFG14b** | | |
| | | | | LB80 | d | COI-FW5 | | | | | | | | |
| | | | | LB81 | d | COI-LB81 | | | d | **amy-CFG14a** | | | | |
| | | | | LB91 | d | COI-FW5 | | | | | d | **ATPS-CFG01b** | | |
| | | | | LB92 | d | COI-LB11 | | | | | | | | |
| | | | | LB98 | d | COI-LB11 | | | d | amy-LB04a amy-LB04b amy-LB98c | | | | |
| | *C. sandai* | Lake Biwa, Japan | CS | CS04 | d | COI-CS04 | d | 28S-CS04a 28S-CS06a 28S-IA3 28S-CS04d | d | amy-CS04a amy-CS04b | | | Sexual | Lineage identification, sperm morphology (1) |
| | | | | CS06 | d | COI-CS06 | d | 28S-CS06a 28S-CS06b 28S-IA3 28S-CS04d | d | amy-CS06a amy-CS06b | | | | |
| | | | | CS09 | d | COI-CS06 | d | 28S-CS04a 28S-CS04d 28S-IA3 | d | amy-CS09a amy-CS06b | | | | |
| | | | | CS10 | d | COI-CS10 | d | 28S-IA3 28S-CS06b 28S-CS06a | d | amy-CS09a amy-CS06b | | | | |
| | | | | CS11 | d | | d | 28S-CS06a | | | | | | |
| | | | | CS12 | d | COI-CS04 | d | 28S-CS06a 28S-CS06b 28S-IA3 28S-CS04d | d | amy-CS09a amy-CS06b | | | | |
| | | | | CS13 | d | COI-CS04 | d | 28S-CS06a 28S-CS06b 28S-IA3 28S-CS04d | d | amy-CS13a amy-CS04a | | | | |
| | | | | CS14 | d | COI-CS04 | d | 28S-CS06a 28S-CS06b 28S-IA3 28S-CS04d | d | amy-CS04a amy-CS14b | | | | |
| | | | | CS15 | d | COI-CS04 | d+c | 28S-CS06a 28S-CS06b 28S-IA3 28S-CS04d | d | amy-CS04a amy-CS14b | | | | |
| | | | | CS18 | d | | d | 28S-CS06a | | | | | | |
| | | | | CS19 | d | COI-CS04 | d | 28S-CS06a 28S-CS06b 28S-IA3 28S-CS04d | d | amy-IA1 | | | | |
| | | | | CS21 | d | COI-CS06 | d | 28S-CS06a 28S-CS06b 28S-IA3 28S-CS04d | d | amy-CS09a amy-CS06b | | | | |
| | | | | CS22 | d | COI-CS04 | d+c | 28S-CS06a 28S-CS06b 28S-IA3 | d | amy-IA1 amy-CS04a | | | | |
| | | | | CS23 | d | COI-CS23 | d | 28S-CS23a 28S-IA3 28S-CS23c | d | amy-CS04a amy-CS14b | | | | |
| | | | | CS24 | d | COI-CS06 | d | 28S-CS06a 28S-CS04d | d | amy-CS09a amy-CS06b | | | | |
| | | | | CS25 | d | COI-CS04 | d | 28S-CS23a 28S-IA3 28S-CS23c | d | amy-CS25a amy-CS06b | | | | |
| | | | | CS26 | d | COI-CS05 | d | 28S-CS04a 28S-CS06a 28S-IA3 | d | amy-IA1 amy-CS04a | | | | |
| | | | | CS27 | d | COI-CS27 | d | 28S-CS06a | d | amy-CS09a amy-CS06b | | | | |
| | | | | CS31 | d | COI-CS04 | d | 28S-CS06a 28S-CS04d | d | amy-CS14b amy-CS04a | | | | |
| | | | | CS32 | d | COI-CS04 | d | 28S-CS06a | d | amy-CS04a | | | | |
| Vietnam | *C.* sp. | Cantho, Vietnam | Vt | Vt01 | d | COI-Vt01 | d | 28S-Vt01a 28S-Vt01b | d | **amy-Vt01a** amy-Vt01b | d | ATPS-Vt01a ATPS-Vt01b | Androgenetic | Sperm morphology (1) |
| | | | | Vt02 | d | COI-Vt02 | d | 28S-Vt02a 28S-Vt01a 28S-Vt02c | d | amy-Vt02a amy-Vt02b | d | ATPS-Vt01a | | |
| | | | | Vt03 | d | COI-Vt01 | d | 28S-Vt03a 28S-Vt02c | d+c | amy-Vt03a amy-C1b | d | ATPS-Vt01a ATPS-Vt01b | | |
| | | | | Vt05 | | | d | 28S-Vt05a 28S-Vt02c | | | | | | |
| | | | | Vt06 | | | d+c | 28S-Vt06a 28S-Vt02c 28S-Vt06c 28S-Vt6d | | | | | | |
| | | | | Vt07 | d | COI-Vt02 | | | | | d | ATPS-Vt01b | | |
| | | | | Vt08 | | | d | 28S-Vt03a 28S-Vt02c | | | | | | |
| | | | | Vt09 | | | d | 28S-Vt06a | | | | | | |
| | | | | Vt10 | d | COI-Vt10 | d | 28S-Vt02c 28S-Vt10b | d | amy-Vt10a amy-Vt10b | d | ATPS-Vt10a ATPS-Vt01a | | |
| | | | | Vt11 | d | COI-Vt02 | d | 28S-Vt11a 28S-Vt02c | d | amy-Vt11a amy-Vt10b | d | ATPS-Vt01b | | |
| | | | | Vt12 | d | COI-Vt01 | d | 28S-Vt01a | d | **amy-Vt01a** amy-Vt12b | d | ATPS-Vt01a | | |
| | | | | Vt13 | d | COI-Vt13 | d | 28S-Vt01a 28S-Vt02c | d | amy-Vt02a amy-Vt13b | d | ATPS-Vt01a ATPS-CR15b | | |
| | | | | Vt14 | | | d | 28S-Vt01a 28S-Vt02c | | | d | ATPS-Vt01a ATPS-Vt01b | | |
| | | | | Vt15 | d | COI-Vt02 | d | 28S-Vt03a 28S-Vt02c | d | amy-Vt15a amy-Vt15b | d | ATPS-Vt01a ATPS-Vt15b | | |
| | | | | Vt16 | d | COI-Vt02 | d | 28S-Vt06c 28S-Vt02c | d | amy-Vt10a amy-Vt10b | d | ATPS-Vt01a ATPS-Vt01b | | |
| | | | | Vt17 | d | COI-Vt02 | d | 28S-Vt17a 28S-Vt02c | d+c | amy-Vt02a amy-Vt17b | d | ATPS-Vt10a ATPS-CR15b | | |
| | | | | Vt19 | d | COI-Vt02 | d | 28S-Vt06a 28S-Vt02c | d | amy-Vt02a **amy-Vt01a** | d | ATPS-Vt01a | | |
| | | | | Vt20 | d | COI-Vt13 | d | 28S-Vt02a 28S-Vt06a | d | amy-Vt12b amy-Vt02a | | | | |
| | *C.* sp. (*C. fluminea* ) | Vietnam | CR | CR01 | d | COI-CR01 | d | 28S-Vt02a 28S-Vt02c | d+c | amy-CR1a **amy-Vt01a** | d | ATPS-Vt01a ATPS-Vt01b | Androgenetic | Lineage identification |
| | | | | CR02 | d | COI-CR02 | d | 28S-Vt02a 28S-Vt02c 28S-CR02c | d+c | amy-CR02a amy-CR02b | d | ATPS-Vt01b | | |
| | | | | CR03 | d | COI-Vt02 | d | 28S-Vt02c | d | **amy-Vt01a** amy-Vt02a | d | ATPS-Vt01b | | |
| | | | | CR04 | d | COI-CR04 | d | 28S-Vt01a 28S-Vt02c | d | amy-CR04a amy-C1b | d | ATPS-Vt01b ATPS-Vt01a | | |
| | | | | CR05 | d | COI-CR01 | d | 28S-CR05a 28S-Vt02c | d | amy-Vt13b amy-Vt10b | d | ATPS-Vt01b ATPS-Vt01a | | |
| | | | | CR06 | | | d | 28S-Vt01a | | | | | | |
| | | | | CR09 | | | d | 28S-Vt06a 28S-Vt02c | | | d | ATPS-Vt01b ATPS-Vt01a | | |
| | | | | CR12 | | | d | 28S-Vt01a 28S-Vt02c | | | | | | |
| | | | | CR13 | d | COI-Vt02 | d | 28S-CR13a 28S-Vt02c | d | amy-Vt02a amy-Vt10b | d | ATPS-Vt01a | | |
| | | | | CR14 | | | | | | | d | ATPS-Vt01a ATPS-Vt01b | | |
| | | | | CR15 | d | COI-Vt02 | d | 28S-Vt06a 28S-CR15b | d+c | amy-CR15a **amy-Vt01a** | d | ATPS-Vt01b ATPS-CR15b | | |
| | | | | CR16 | | | d | 28S-Vt01a 28S-Vt02c | | | | | | |
| | | | | CR17 | d | COI-Vt02 | d | 28S-Vt06a 28S-Vt02c | d | **amy-Vt01a** amy-Vt02a | d | ATPS-Vt01a ATPS-Vt01b | | |
| | | | | CR18 | | | d | 28S-Vt01a 28S-Vt02c | | | d | ATPS-Vt01a ATPS-Vt01b | | |
| | | | | CR19 | | | d | 28S-CR19a 28S-Vt02c | | | | | | |
| | | | | CR20 | | | d | 28S-CR20a | | | d | ATPS-Vt01b | | |
| | | | | CR21 | | | | | | | d | ATPS-Vt01b ATPS-Vt01a | | |
| | | | | CR22 | d | COI-Vt13 | d | 28S-CR22a 28S-Vt02c | d | amy-Vt02a amy-Vt13b | | | | |
| | | | | CR23 | | | d | 28S-CR23a 28S-Vt02c | | | d | ATPS-Vt01b ATPS-Vt01a | | |
| | | | | CR24 | d | COI-Vt02 | d | 28S-CR24a 28S-Vt01a 28S-Vt06a | d | amy-Vt12b amy-CR24b | d | ATPS-Vt01a ATPS-Vt01b | | |
| | | | | CR25 | | | d | 28S-Vt03a | | | d | ATPS-Vt01b | | |
| | | | | CR26 | d | COI-CR26 | d | 28S-Vt06a 28S-Vt02c | | | d+c | ATPS-Vt01b ATPS-CR26b | | |
| | | | CRB | CRB01 | d | COI-CR02 | d | 28S-Vt06a 28S-Vt02c | | | d | ATPS-Vt01a ATPS-CRB01b | | |
| | | | | CRB02 | | | d | 28S-Vt03a | | | | | | |
| | | | | CRB03 | | | d | 28S-Vt03a 28S-Vt02c | | | | | | |
| | | | | CRB04 | d | COI-CRB4 | d | 28S-Vt06a 28S-CRB04b | d | amy-CRB04a amy-CRB04b | d | ATPS-Vt01a ATPS-CRB4b | | |
| | | | | CRB05 | d | COI-Vt02 | d | 28S-Vt01a 28S-Vt02c | d | amy-Vt02a amy-CRB05b | d | ATPS-CR26b ATPS-Vt01b | | |
| | | | | CRB06 | | | d | 28S-CRB06a 28S-Vt02c | | | d | ATPS-Vt01a ATPS-Vt01b | | |
| | | | | CRB07 | | | d | 28S-Vt01a 28S-Vt02c | | | | | | |
| | | | | CRB09 | d | COI-CRB09 | d | 28S-Vt03a 28S-Vt02c | | **amy-Vt01a** amy-Vt02a | | | | |
| | | | | CRB10 | | | d | 28S-Vt03a 28S-Vt02c | | | d | ATPS-Vt01a ATPS-Vt01b | | |
| | | | | CRB11 | | | d | 28S-Vt6d 28S-CRB11b | | | | | | |
| | | | | CRB12 | d | COI-Vt02 | d | 28S-Vt06a 28S-Vt02c | d | amy-Vt02a amy-CR24b | d+c | ATPS-CR26b ATPS-Vt01a | | |
| | | | | CRB13 | | | d | 28S-CR05a 28S-Vt02c | | | d | ATPS-Vt01b ATPS-CRB4b | | |
| | | | | CRB14 | d | COI-Vt02 | d | 28S-Vt02a | d | amy-Vt10b | d | ATPS-Vt01a | | |
| | | | | CRB15 | d | COI-Vt02 | d | 28S-Vt02c | | | | | | |
| | | | | CRB16 | d | COI-Vt02 | | | | | d | ATPS-Vt01b ATPS-CR15b | | |
| | | | | CRB17 | | | | | d | **amy-Vt01a** | | | | |
| | | | | CRB18 | | | d | 28S-Vt06a 28S-Vt02c | | | d | ATPS-Vt01a ATPS-CRB4b | | |
| | | | | CRB19 | | | | | | | d | ATPS-CR26b ATPS-Vt01b | | |
| | | | | CRB20 | | | d | 28S-CR19a 28S-Vt02c | | | d | ATPS-Vt01a | | |
| | | | | CRB21 | | | d | 28S-Vt06c 28S-Vt02c | | | | | | |
| Indonesia | *C. loehensis* | Lake Mahalona, Sulawesi, Indonesia | ZMB | 190852* | d | COI-190768 | | | | | | | Sexual | Reproductive morphology (2) |
| | | Lake Lontoa, Sulawesi, Indonesia | ZMB | 190768* | d | COI-190768 | | | | | | | Sexual | Reproductive morphology (2) |
| | *C. matannensis* | Lake Towuti, Sulawesi, Indonesia | ZMB | 191042* | d | COI-191042 | | | | | | | Sexual | Reproductive morphology (2) |
| | *C. tobae* | Indonesia | ZMB | 103027* | | | | | c | **amy-Vt01a** | | | ? | |
| South_Africa | *C. fluminalis africana* | Mooi River, Potchefstroom, South Africa | ZA | ZA01 | d | COI-ZA01 | d+c | 28S-ZA01a 28S-ZA01b 28S-ZA01c | d | amy-C1b | d | ATPS-C1a | ? | |
| | | | | ZA02 | | | | | | | d | ATPS-C1a | | |
| | | | | ZA03 | d | COI-ZA03 | d | 28S-ZA01a 28S-ZA01c | | | | | | |
| | | | | ZA04 | | | d | 28S-ZA01a 28S-ZA01c | | | d | ATPS-C1a | | |
| | | | | ZA05 | d | COI-ZA03 | d | 28S-ZA01a 28S-ZA01c | d | amy-C1b | d | ATPS-C1a | | |
| | | | | ZA06 | d | COI-ZA03 | d | 28S-ZA01a 28S-ZA01c | d | amy-C1b | d | ATPS-C1a | | |
| | | | | ZA07 | d | COI-ZA03 | d | 28S-ZA01a 28S-ZA01c | d+c | amy-C1b | | | | |
| | | | | ZA08 | | | d | 28S-ZA01a 28S-ZA01c | | | d | ATPS-C1a | | |
| | | | | ZA09 | d | COI-ZA03 | d | 28S-ZA01a 28S-ZA01c | d | amy-C1b | d | ATPS-C1a | | |
| | | | | ZA10 | d | COI-ZA03 | d | 28S-ZA01a 28S-ZA01c | d | amy-C1b | d | ATPS-C1a | | |
| | | | | ZA11 | d | COI-ZA03 | d | 28S-ZA01a 28S-ZA01c | d | amy-C1b | d | ATPS-C1a | | |
| | | | | ZA12 | d | COI-ZA03 | d+c | 28S-ZA01a 28S-ZA01c | d+c | amy-C1b | d | ATPS-C1a | | |
| | | | | ZA13 | d | COI-ZA03 | d | 28S-ZA01a 28S-ZA01c | d | amy-C1b | d | ATPS-C1a | | |
| | | | | ZA14 | | | d | 28S-ZA01a 28S-ZA01c | | | d | ATPS-C1a | | |
| | | | | ZA15 | d | COI-ZA03 | d | 28S-ZA01a 28S-ZA01c | d | amy-C1b | d | ATPS-C1a | | |
| | | | | ZA16 | d | COI-ZA03 | d | 28S-ZA01a 28S-ZA01c | d | amy-C1b | d | ATPS-C1a | | |
| | | | | ZA17 | d | COI-ZA03 | d | 28S-ZA01a 28S-ZA01c | d | amy-C1b | d | ATPS-C1a | | |
| | | | | ZA18 | d | COI-ZA03 | d | 28S-ZA01a 28S-ZA01c | d | amy-C1b | d | ATPS-C1a | | |
| | | | | ZA19 | d | COI-ZA03 | d | 28S-ZA01a 28S-ZA01c | d | amy-C1b | d | ATPS-C1a | | |
| | | | | ZA20 | | | | | | | d | ATPS-C1a | | |
| Madagascar | *C. madagascariensis* | Madagascar | Mad | Mad2 | d | COI-Mad2 | d | **28S-CI204c** **28S-Hw4b** 28S-IA3 | | | | | ? | |
| | | | | Mad3 | | | d | **28S-CI204c** **28S-Hw4b** 28S-IA3 | | | | | | |
| | | | | Mad6 | | | d+c | **28S-CI204c** **28S-Hw4b** 28S-IA3 | | | | | | |
| **Total** | | | | | | **N=251; N haplotypes=44** | | **N individuals=266; N heterozygous=224; N haplotypes=56** | | **N individuals=237; N heterozygous=201; N haplotypes=74** | | **N individuals=248; N heterozygous=148; N haplotypes=40** | | |


(1) Pigneur L-M, Etoundi E, Aldridge D, Marescaux J, Yasuda N, Van Doninck K (2014) Genetic uniformity and long-distance clonal dispersal in the invasive androgenetic *Corbicula* clams. *Molecular Ecology* 23, 5102-16

(2) Hedtke SM, Glaubrecht M, Hillis DM (2011) Rare gene capture in predominantly androgenetic species. *Proceedings of the National Academy of Sciences, USA*. **108**, 9520-9524

(3) Etoundi E, unpublished data

(4) Byrne M, Phelps H, Church T, Adair V, Selvakumaraswamy P, Potts J (2000) Reproduction and development of the freshwater clam *Corbicula australis* in southeast Australia. *Hydrobiologia*. **418**, 185-197
