## Supplementary material for "Substantial genetic mixing among sexual and androgenetic lineages within the clam genus *Corbicula*": Table S3

Table S3: list of the 141 analyzed individuals for which all 4 markers are available. The color code corresponds to the four invasive lineages: alleles retrieved from *C.* sp. form A/R, B, C/S and Rlc are respectively in red, blue, orange and green. Invasive alleles (IA), defined here as alleles found in at least two different invasive lineages, are shown in purple. Alleles shared between idiotypes (referring to Figure 4) are in bold. For additional details, see Table S2.

| Taxon | Location | Population code | Collection number | COI haplotypes | 28S haplotypes | | | α-amylase haplotypes | | | | ATPS-α haplotypes | | |
|---|---|---|---|---|---|---|---|---|---|---|---|---|---|---|
| *C.* sp. form A/R (*C. fluminea* , white form) | San Gabriel River, USA | AA | AA11 | COI-FW5 | 28S-IA1 | 28S-IA2 | 28S-IA3 | amy-IA1 | amy-IA2 | amy-AA10c | | ATPS-AA11a | ATPS-IA1 | |
| | | | AA12 | COI-FW5 | 28S-IA1 | 28S-IA2 | 28S-IA3 | amy-IA1 | amy-IA2 | amy-AA10c | | ATPS-AA11a | ATPS-IA1 | ATPS-AA12c |
| | | | AA13 | COI-FW5 | 28S-IA1 | 28S-IA2 | 28S-IA3 | amy-IA1 | amy-IA2 | amy-AA10c | | ATPS-AA11a | ATPS-IA1 | ATPS-AA12c |
| *C.* sp. form B (purple form) | San Gabriel River, USA | AB | AB11 | COI-FW1 | 28S-AB11a | 28S-AB11b | | amy-IA1 | amy-IA2 | amy-IA3 | amy-AB12d | ATPS-AB11a | ATPS-AB11b | |
| | | | AB13 | COI-FW1 | 28S-AB11a | 28S-AB11b | | amy-IA1 | amy-IA2 | amy-IA3 | amy-AB12d | ATPS-AB11a | ATPS-AB11b | |
| | | | AB14 | COI-FW1 | 28S-AB11a | 28S-AB11b | | amy-IA1 | amy-IA2 | amy-IA3 | amy-AB12d | ATPS-AB11a | ATPS-AB11b | |
| | | | AB15 | COI-FW1 | 28S-AB11a | 28S-AB11b | | amy-IA1 | amy-IA2 | amy-IA3 | amy-AB12d | ATPS-AB11a | ATPS-AB11b | |
| | USA | yy | yy12 | COI-FW1 | 28S-AB11a | 28S-AB11b | 28S-yy12c | amy-IA3 | amy-AB12d | | | ATPS-AB11a | | |
| *C.* sp. form C/S (*C. fluminalis, C. largilierti* ) | Argentina | C | C1 | COI-FW17 | 28S-C1a | 28S-IA1 | 28S-C1c | amy-C1a | amy-C1b | | | ATPS-C1a | ATPS-IA1 | |
| | | | C2 | COI-FW17 | 28S-C1a | 28S-IA1 | 28S-C1c | amy-C1a | amy-C1b | | | ATPS-C1a | ATPS-IA1 | |
| | | | C3 | COI-FW17 | 28S-C1a | 28S-IA1 | 28S-C1c | amy-C1a | amy-C1b | | | ATPS-C1a | ATPS-IA1 | |
| | | | C4 | COI-FW17 | 28S-C1a | 28S-IA1 | 28S-C1c | amy-C1a | amy-C1b | | | ATPS-C1a | ATPS-IA1 | |
| | Saône River, France | Sa | Sa22 | COI-FW17 | 28S-C1a | 28S-IA1 | 28S-C1c | amy-C1a | amy-C1b | | | ATPS-C1a | ATPS-IA1 | |
| *C.* sp. form Rlc | Doubs River, France | Rlc | Db22 | COI-FW4 | 28S-IA2 | | | amy-Db22a | amy-IA3 | | | ATPS-Db22a | ATPS-Db22b | |
| | | | Db23 | COI-FW4 | 28S-IA2 | | | amy-Db22a | amy-IA3 | | | ATPS-Db22a | ATPS-Db22b | |
| | | | Db24 | COI-FW4 | 28S-IA2 | 28S-IA3 | | amy-Db22a | amy-IA3 | | | ATPS-Db22a | ATPS-Db22b | |
| | | | Db25 | COI-FW5 | 28S-IA2 | | | amy-Db22a | amy-IA3 | | | ATPS-Db22a | ATPS-Db22b | |
| | | | Db26 | COI-FW4 | 28S-IA2 | | | amy-Db22a | amy-IA3 | | | ATPS-Db22a | ATPS-Db22b | |
| *C. japonica* | Japan | CJ | CJ03 | COI-CJ03 | 28S-CJ03 | | | amy-CJ03a | amy-CJ03b | | | ATPS-CJ03a | ATPS-CJ03b | |
| | | | CJ04 | COI-CJ03 | 28S-CJ03 | | | amy-CJ04a | amy-CJ04b | | | ATPS-CJ03a | ATPS-CJ04b | |
| | | | CJ07 | COI-CJ07 | 28S-CJ07a | 28S-CJ03 | | amy-CJ07a | amy-CJ07b | | | ATPS-CJ03b | ATPS-CJ07b | |
| | | | CJ09 | COI-CJ09 | 28S-CJ07a | 28S-CJ03 | | amy-CJ09a | amy-CJ09b | | | ATPS-CJ03a | ATPS-CJ04b | |
| | | | CJ10 | COI-CJ09 | 28S-CJ07a | 28S-CJ03 | | amy-CJ10a | amy-CJ10b | | | ATPS-CJ03a | | |
| | | | CJ11 | COI-CJ09 | 28S-CJ07a | 28S-CJ03 | | amy-CJ04a | amy-CJ07a | | | ATPS-CJ03a | ATPS-CJ04b | |
| | | | CJ12 | COI-CJ12 | 28S-CJ07a | 28S-CJ03 | | amy-CJ12a | amy-CJ04a | | | ATPS-CJ03b | ATPS-CJ12b | |
| | | | CJ15 | COI-CJ03 | 28S-CJ07a | 28S-CJ03 | | amy-CJ04b | amy-CJ03b | | | ATPS-CJ03b | ATPS-CJ04b | |
| | | | CJ18 | COI-CJ18 | 28S-CJ07a | | | amy-CJ04b | amy-CJ03b | | | ATPS-CJ18a | ATPS-CJ18b | |
| | | | CJ19 | COI-CJ19 | 28S-CJ07a | 28S-CJ03 | | amy-CJ19 | | | | ATPS-CJ03a | | |
| *C* . sp. (*C. fluminea /leana* morphs) | Ehime Shigenobu Takubo, Japan | EHM | EHM102 | COI-FW1 | 28S-AB11a | | | amy-EHM102a | amy-AB12d | | | ATPS-CFG14a | ATPS-Db22a | |
| | | | EHM103 | COI-FW1 | 28S-AB11a | | | amy-EHM102a | amy-AB12d | | | ATPS-CFG14a | ATPS-Db22a | |
| | | | EHM104 | COI-FW1 | 28S-AB11a | | | amy-EHM102a | amy-AB12d | | | ATPS-CFG14a | ATPS-Db22a | |
| | | | EHM105 | COI-FW1 | 28S-AB11a | | | amy-EHM102a | amy-AB12d | | | ATPS-CFG14a | ATPS-Db22a | |
| | | | EHM106 | COI-FW1 | 28S-AB11a | | | amy-EHM102a | amy-AB12d | | | ATPS-CFG14a | | |
| | | | EHM107 | COI-FW1 | 28S-AB11a | | | amy-EHM102a | amy-AB12d | | | ATPS-CFG14a | ATPS-Db22a | |
| | | | EHM108 | COI-FW1 | 28S-AB11a | | | amy-EHM102a | amy-AB12d | | | ATPS-CFG14a | ATPS-Db22a | |
| | | | EHM109 | COI-FW1 | 28S-AB11a | | | amy-EHM102a | amy-AB12d | | | ATPS-CFG14a | ATPS-Db22a | |
| | | | EHM110 | COI-FW1 | 28S-AB11a | | | amy-EHM102a | amy-AB12d | | | ATPS-CFG14a | ATPS-Db22a | |
| | | | EHM111 | COI-FW1 | 28S-AB11a | | | amy-EHM102a | amy-AB12d | | | ATPS-CFG14a | ATPS-Db22a | |
| | | | EHM112 | COI-FW1 | 28S-AB11a | | | amy-EHM102a | amy-AB12d | | | ATPS-CFG14a | ATPS-Db22a | |
| | | | EHM113 | COI-FW1 | 28S-AB11a | | | amy-EHM102a | amy-AB12d | | | ATPS-CFG14a | ATPS-Db22a | |
| | | | EHM119 | COI-FW1 | 28S-AB11a | | | amy-EHM102a | amy-AB12d | | | ATPS-CFG14a | ATPS-Db22a | |
| | | | EHM120 | COI-FW1 | 28S-AB11a | | | amy-EHM102a | amy-AB12d | | | ATPS-CFG14a | ATPS-Db22a | |
| | | | EHM121 | COI-FW1 | 28S-AB11a | | | amy-EHM102a | amy-AB12d | | | ATPS-CFG14a | ATPS-Db22a | |
| | | | EHM122 | COI-FW1 | 28S-AB11a | | | amy-EHM102a | amy-AB12d | | | ATPS-CFG14a | ATPS-Db22a | |
| | Miyazaki Gongenbaru Canal, Japan | KMT | KMT205 | COI-FW5 | 28S-AB11a | **28S-KMT205a** | 28S-AB11b | amy-KMT205 | | | | ATPS-Db22b | **ATPS-KMT205b** | |
| | | | KMT208 | COI-FW5 | 28S-AB11a | **28S-KMT205a** | 28S-AB11b | amy-KMT205 | | | | ATPS-Db22b | **ATPS-KMT205b** | |
| | | | KMT213 | COI-FW5 | 28S-AB11a | 28S-yy12c | | amy-IA2 | amy-Db22a | | | **ATPS-KMT205l** | **ATPS-CFG01b** | ATPS-AB11a |
| | | | KMT214 | COI-FW5 | 28S-AB11a | 28S-yy12c | | amy-IA2 | amy-Db22a | | | **ATPS-KMT205l** | **ATPS-CFG01b** | ATPS-AB11a |
| | | | KMT215 | COI-FW5 | 28S-AB11a | 28S-yy12c | | amy-IA2 | amy-Db22a | | | **ATPS-KMT205l** | **ATPS-CFG01b** | ATPS-AB11a |
| | | | KMT216 | COI-FW5 | 28S-AB11a | 28S-yy12c | | amy-IA2 | amy-Db22a | | | **ATPS-KMT205l** | **ATPS-CFG01b** | ATPS-AB11a |
| | | | KMT217 | COI-FW5 | 28S-AB11a | 28S-yy12c | | amy-IA2 | amy-Db22a | | | **ATPS-KMT205l** | **ATPS-CFG01b** | ATPS-AB11a |
| | | | KMT218 | COI-FW5 | 28S-AB11a | 28S-yy12c | | amy-IA2 | amy-Db22a | | | **ATPS-KMT205l** | **ATPS-CFG01b** | ATPS-AB11a |
| | | | KMT219 | COI-FW5 | 28S-AB11a | 28S-yy12c | | amy-IA2 | amy-Db22a | | | **ATPS-KMT205l** | **ATPS-CFG01b** | ATPS-AB11a |
| | | | KMT221 | COI-FW5 | 28S-AB11a | 28S-yy12c | | amy-IA2 | amy-Db22a | | | **ATPS-KMT205l** | **ATPS-CFG01b** | ATPS-AB11a |
| | | | KMT222 | COI-FW5 | 28S-AB11a | 28S-yy12c | | amy-IA2 | amy-Db22a | | | **ATPS-KMT205l** | **ATPS-CFG01b** | ATPS-AB11a |
| | | | KMT224 | COI-FW5 | 28S-AB11a | 28S-yy12c | | amy-IA2 | amy-Db22a | | | **ATPS-KMT205l** | **ATPS-CFG01b** | ATPS-AB11a |
| | Ehime, Japan | Cl | Cl204 | COI-Cl204 | 28S-IA3 | 28S-Cl204b | **28S-Cl204c** | **amy-CFG14a** | amy-Db22a | amy-Cl204c | | ATPS-AB11a | | |
| | | | Cl205 | COI-Cl204 | 28S-IA3 | 28S-Cl204b | **28S-Cl204c** | **amy-CFG14a** | amy-Db22a | amy-Cl204c | | ATPS-AB11a | | |
| | | | Cl206 | COI-Cl204 | 28S-IA3 | 28S-Cl204b | **28S-Cl204c** | **amy-CFG14a** | amy-Db22a | amy-Cl204c | | ATPS-AB11a | | |
| | | | Cl207 | COI-Cl204 | 28S-IA3 | 28S-Cl204b | **28S-Cl204c** | **amy-CFG14a** | amy-Db22a | amy-Cl204c | | ATPS-AB11a | | |
| | | | Cl208 | COI-Cl204 | 28S-IA3 | 28S-Cl204b | **28S-Cl204c** | **amy-CFG14a** | amy-Db22a | amy-Cl204c | | ATPS-AB11a | | |
| | | | Cl210 | COI-Cl204 | 28S-IA3 | 28S-Cl204b | **28S-Cl204c** | **amy-CFG14a** | amy-Db22a | amy-Cl204c | | ATPS-AB11a | | |
| | | | Cl211 | COI-Cl204 | 28S-IA3 | 28S-Cl204b | **28S-Cl204c** | **amy-CFG14a** | amy-Db22a | amy-Cl204c | | ATPS-AB11a | | |
| | | | Cl212 | COI-Cl204 | 28S-IA3 | 28S-Cl204b | **28S-Cl204c** | **amy-CFG14a** | amy-Cl204c | | | ATPS-AB11a | | |
| | | | Cl213 | COI-Cl204 | 28S-IA3 | 28S-Cl204b | **28S-Cl204c** | **amy-CFG14a** | amy-Db22a | amy-Cl204c | | ATPS-AB11a | | |
| | | | Cl214 | COI-Cl204 | 28S-IA3 | 28S-Cl204b | **28S-Cl204c** | **amy-CFG14a** | amy-Db22a | amy-Cl204c | | ATPS-AB11a | | |
| | | | Cl215 | COI-Cl204 | 28S-IA3 | 28S-Cl204b | **28S-Cl204c** | **amy-CFG14a** | amy-Db22a | amy-Cl204c | | ATPS-AB11a | | |
| | | | Cl216 | COI-Cl204 | 28S-IA3 | 28S-Cl204b | **28S-Cl204c** | **amy-CFG14a** | amy-Db22a | amy-Cl204c | | ATPS-AB11a | | |
| | | | Cl218 | COI-Cl204 | 28S-IA3 | 28S-Cl204b | **28S-Cl204c** | **amy-CFG14a** | amy-Db22a | amy-Cl204c | | ATPS-AB11a | | |
| | | | Cl220 | COI-Cl204 | 28S-IA3 | 28S-Cl204b | **28S-Cl204c** | **amy-CFG14a** | amy-Db22a | amy-Cl204c | | ATPS-AB11a | | |
| | | | Cl221 | COI-Cl204 | 28S-IA3 | 28S-Cl204b | **28S-Cl204c** | **amy-CFG14a** | amy-Db22a | amy-Cl204c | | ATPS-AB11a | | |
| | | | Cl222 | COI-Cl204 | 28S-IA3 | 28S-Cl204b | **28S-Cl204c** | **amy-CFG14a** | amy-Db22a | amy-Cl204c | | ATPS-AB11a | | |
| | | | Cl223 | COI-Cl204 | 28S-IA3 | 28S-Cl204b | **28S-Cl204c** | **amy-CFG14a** | amy-Db22a | amy-Cl204c | | ATPS-AB11a | | |
| | | | Cl224 | COI-Cl204 | 28S-IA3 | 28S-Cl204b | **28S-Cl204c** | **amy-CFG14a** | amy-Db22a | amy-Cl204c | | ATPS-AB11a | | |
| *C.* sp. (*C. fluminea* yellow morph) | Sakanai River, Matsusaka, Japan | CFY | CFY03 | COI-FW5 | 28S-IA2 | 28S-IA1 | | amy-CFY01a | amy-Db22a | | | ATPS-AB11a | | |
| | | | CFY04 | COI-FW5 | 28S-IA2 | 28S-IA1 | | amy-CFY01a | amy-Db22a | | | ATPS-AB11a | | |
| | | | CFY05 | COI-FW5 | 28S-IA2 | 28S-IA1 | | amy-CFY01a | amy-Db22a | | | ATPS-AB11a | | |
| | | | CFY07 | COI-Cl204 | 28S-IA2 | 28S-IA1 | | amy-CFY01a | amy-Db22a | | | ATPS-AB11a | | |
| | | | CFY09 | COI-FW5 | 28S-IA2 | 28S-IA1 | | amy-CFY01a | amy-Db22a | | | ATPS-AB11a | | |
| | | | CFY10 | COI-FW5 | 28S-IA2 | 28S-IA1 | | amy-CFY01a | amy-Db22a | | | ATPS-AB11a | | |
| | | | CFY11 | COI-FW5 | 28S-IA2 | 28S-IA1 | | amy-CFY01a | amy-Db22a | | | ATPS-AB11a | | |
| | | | CFY12 | COI-FW5 | 28S-IA2 | 28S-IA1 | | amy-CFY01a | amy-Db22a | | | ATPS-AB11a | | |
| *C.* sp. (*C. fluminea* green morph) | Tokida River, Watarai, Japan | CFG | CFG01 | COI-FW5 | 28S-Cl204b | 28S-ggg2a | | amy-IA1 | amy-Db22a | | | ATPS-Db22a | **ATPS-CFG01b** | |
| | | | CFG02 | COI-FW5 | 28S-Cl204b | 28S-ggg2a | | amy-IA1 | amy-Db22a | | | ATPS-Db22a | **ATPS-CFG01b** | |
| | | | CFG03 | COI-FW5 | 28S-Cl204b | 28S-ggg2a | | amy-IA1 | amy-Db22a | | | ATPS-Db22a | **ATPS-CFG01b** | |
| | | | CFG04 | COI-FW5 | 28S-Cl204b | 28S-ggg2a | | amy-IA1 | amy-Db22a | | | ATPS-Db22a | **ATPS-CFG01b** | |
| | | | CFG05 | COI-FW5 | 28S-Cl204b | 28S-ggg2a | | amy-IA1 | amy-Db22a | | | ATPS-Db22a | **ATPS-CFG01b** | |
| | | | CFG06 | COI-FW5 | 28S-Cl204b | 28S-ggg2a | | amy-IA1 | amy-Db22a | | | ATPS-Db22a | **ATPS-CFG01b** | |
| | | | CFG07 | COI-FW5 | 28S-Cl204b | 28S-ggg2a | | amy-IA1 | amy-Db22a | | | ATPS-Db22a | **ATPS-CFG01b** | |
| | | | CFG08 | COI-FW5 | 28S-Cl204b | 28S-ggg2a | | amy-IA1 | amy-Db22a | | | ATPS-Db22a | **ATPS-CFG01b** | |
| | | | CFG09 | COI-FW5 | 28S-Cl204b | 28S-ggg2a | | amy-IA1 | amy-Db22a | | | ATPS-Db22a | **ATPS-CFG01b** | |
| | | | CFG10 | COI-FW5 | 28S-Cl204b | 28S-ggg2a | | amy-IA1 | amy-Db22a | | | ATPS-Db22a | **ATPS-CFG01b** | |
| | | | CFG11 | COI-FW5 | 28S-Cl204b | 28S-ggg2a | | amy-IA1 | amy-Db22a | | | ATPS-Db22a | **ATPS-CFG01b** | |
| | | | CFG12 | COI-FW5 | 28S-Cl204b | 28S-ggg2a | | amy-IA1 | amy-Db22a | | | ATPS-Db22a | **ATPS-CFG01b** | |
| | | | CFG13 | COI-FW5 | 28S-Cl204b | 28S-ggg2a | | amy-IA1 | amy-Db22a | | | ATPS-Db22a | **ATPS-CFG01b** | |
| | Kushida River, Matsusaka, Japan | CFG | CFG15 | COI-FW5 | 28S-Cl204b | 28S-ggg2a | | amy-IA1 | amy-Db22a | | | ATPS-Db22a | **ATPS-CFG01b** | |
| | | | CFG16 | COI-FW1 | 28S-AB11a | 28S-IA2 | | **amy-CFG14a** | amy-Db22a | | | ATPS-CFG14a | **ATPS-CFG14b** | |
| | | | CFG17 | COI-FW1 | 28S-AB11a | | | **amy-CFG14a** | amy-Db22a | | | ATPS-CFG14a | **ATPS-CFG14b** | |
| | | | CFG18 | COI-FW1 | 28S-AB11a | | | **amy-CFG14a** | amy-Db22a | | | ATPS-CFG14a | **ATPS-CFG14b** | |
| | | | CFG19 | COI-FW1 | 28S-AB11a | 28S-IA2 | | **amy-CFG14a** | amy-Db22a | | | ATPS-CFG14a | **ATPS-CFG14b** | |
| | | | CFG20 | COI-FW1 | 28S-AB11a | | | **amy-CFG14a** | amy-Db22a | | | ATPS-CFG14a | **ATPS-CFG14b** | |
| | | | CFG21 | COI-FW1 | 28S-AB11a | | | **amy-CFG14a** | | | | ATPS-CFG14a | **ATPS-CFG14b** | |
| | | | CFG22 | COI-FW1 | 28S-AB11a | 28S-IA2 | | **amy-CFG14a** | amy-Db22a | | | ATPS-CFG14a | **ATPS-CFG14b** | |
| | | | CFG23 | COI-FW1 | 28S-AB11a | 28S-IA2 | | **amy-CFG14a** | amy-Db22a | | | ATPS-CFG14a | **ATPS-CFG14b** | |
| *C.* sp. | Cantho, Vietnam | Vt | Vt01 | COI-Vt01 | 28S-Vt01a | 28S-Vt01b | | amy-Vt01a | amy-Vt01b | | | ATPS-Vt01a | ATPS-Vt01b | |
| | | | Vt02 | COI-Vt02 | 28S-Vt02a | 28S-Vt01a | 28S-Vt02c | amy-Vt02a | amy-Vt02b | | | ATPS-Vt01a | | |
| | | | Vt03 | COI-Vt01 | 28S-Vt03a | 28S-Vt02c | | amy-Vt03a | amy-C1b | | | ATPS-Vt01a | ATPS-Vt01b | |
| | | | Vt10 | COI-Vt10 | 28S-Vt02c | 28S-Vt10b | | amy-Vt10a | amy-Vt10b | | | ATPS-Vt10a | ATPS-Vt01a | |
| | | | Vt11 | COI-Vt02 | 28S-Vt11a | 28S-Vt02c | | amy-Vt11a | amy-Vt10b | | | ATPS-Vt01b | | |
| | | | Vt12 | COI-Vt01 | 28S-Vt01a | | | amy-Vt01a | amy-Vt12b | | | ATPS-Vt01a | | |
| | | | Vt13 | COI-Vt13 | 28S-Vt01a | 28S-Vt02c | | amy-Vt02a | amy-Vt13b | | | ATPS-Vt01a | ATPS-CR15b | |
| | | | Vt15 | COI-Vt02 | 28S-Vt03a | 28S-Vt02c | | amy-Vt15a | amy-Vt15b | | | ATPS-Vt01a | ATPS-Vt15b | |
| | | | Vt16 | COI-Vt02 | 28S-Vt06c | 28S-Vt02c | | amy-Vt10a | amy-Vt10b | | | ATPS-Vt01a | ATPS-Vt01b | |
| | | | Vt17 | COI-Vt02 | 28S-Vt17a | 28S-Vt02c | | amy-Vt02a | amy-Vt17b | | | ATPS-Vt10a | ATPS-CR15b | |
| | | | Vt19 | COI-Vt02 | 28S-Vt06a | 28S-Vt02c | | amy-Vt02a | amy-Vt01a | | | ATPS-Vt01a | | |
| *C.* sp. (*C. fluminea* ) | Vietnam | CR | CR01 | COI-CR01 | 28S-Vt02a | 28S-Vt02c | | amy-CR01a | amy-Vt01a | | | ATPS-Vt01a | ATPS-Vt01b | |
| | | | CR02 | COI-CR02 | 28S-Vt02a | 28S-Vt02c | 28S-CR02c | amy-CR02a | amy-CR02b | | | ATPS-Vt01b | | |
| | | | CR03 | COI-Vt02 | 28S-Vt02c | | | amy-Vt01a | amy-Vt02a | | | ATPS-Vt01b | | |
| | | | CR04 | COI-CR04 | 28S-Vt01a | 28S-Vt02c | | amy-CR04a | amy-C1b | | | ATPS-Vt01b | ATPS-Vt01a | |
| | | | CR05 | COI-CR01 | 28S-CR05a | 28S-Vt02c | | amy-Vt13b | amy-Vt10b | | | ATPS-Vt01b | ATPS-Vt01a | |
| | | | CR13 | COI-Vt02 | 28S-CR13a | 28S-Vt02c | | amy-Vt02a | amy-Vt10b | | | ATPS-Vt01a | | |
| | | | CR15 | COI-Vt02 | 28S-Vt06a | 28S-CR15b | | amy-CR15a | amy-Vt01a | | | ATPS-Vt01b | ATPS-CR15b | |
| | | | CR17 | COI-Vt02 | 28S-Vt06a | 28S-Vt02c | | amy-Vt01a | amy-Vt02a | | | ATPS-Vt01a | ATPS-Vt01b | |
| | | | CR24 | COI-Vt02 | 28S-CR24a | 28S-Vt01a | 28S-Vt06a | amy-Vt12b | amy-CR24b | | | ATPS-Vt01a | ATPS-Vt01b | |
| | | CRB | CRB04 | COI-CRB04 | 28S-Vt06a | 28S-CRB04b | | amy-CRB04a | amy-CRB04b | | | ATPS-Vt01a | ATPS-CRB4b | |
| | | | CRB05 | COI-Vt02 | 28S-Vt01a | 28S-Vt02c | | amy-Vt02a | amy-CRB05b | | | ATPS-CRB05a | ATPS-Vt01b | |
| | | | CRB12 | COI-Vt02 | 28S-Vt06a | 28S-Vt02c | | amy-Vt02a | amy-CR24b | | | ATPS-CRB05a | ATPS-Vt01a | |
| | | | CRB14 | COI-Vt02 | 28S-Vt02a | | | amy-Vt10b | | | | ATPS-Vt01a | | |
| *C. fluminalis africana* | Mooi River, Potchefstroom, South Africa | ZA | ZA01 | COI-ZA01 | 28S-ZA01a | 28S-ZA01b | 28S-ZA01c | amy-C1b | | | | ATPS-C1a | | |
| | | | ZA05 | COI-ZA03 | 28S-ZA01a | 28S-ZA01c | | amy-C1b | | | | ATPS-C1a | | |
| | | | ZA06 | COI-ZA03 | 28S-ZA01a | 28S-ZA01c | | amy-C1b | | | | ATPS-C1a | | |
| | | | ZA09 | COI-ZA03 | 28S-ZA01a | 28S-ZA01c | | amy-C1b | | | | ATPS-C1a | | |
| | | | ZA10 | COI-ZA03 | 28S-ZA01a | 28S-ZA01c | | amy-C1b | | | | ATPS-C1a | | |
| | | | ZA11 | COI-ZA03 | 28S-ZA01a | 28S-ZA01c | | amy-C1b | | | | ATPS-C1a | | |
| | | | ZA12 | COI-ZA03 | 28S-ZA01a | 28S-ZA01c | | amy-C1b | | | | ATPS-C1a | | |
| | | | ZA13 | COI-ZA03 | 28S-ZA01a | 28S-ZA01c | | amy-C1b | | | | ATPS-C1a | | |
| | | | ZA15 | COI-ZA03 | 28S-ZA01a | 28S-ZA01c | | amy-C1b | | | | ATPS-C1a | | |
| | | | ZA16 | COI-ZA03 | 28S-ZA01a | 28S-ZA01c | | amy-C1b | | | | ATPS-C1a | | |
| | | | ZA17 | COI-ZA03 | 28S-ZA01a | 28S-ZA01c | | amy-C1b | | | | ATPS-C1a | | |
| | | | ZA18 | COI-ZA03 | 28S-ZA01a | 28S-ZA01c | | amy-C1b | | | | ATPS-C1a | | |
| | | | ZA19 | COI-ZA03 | 28S-ZA01a | 28S-ZA01c | | amy-C1b | | | | ATPS-C1a | | |
