## Supplementary material for "Substantial genetic mixing among sexual and androgenetic lineages within the clam genus *Corbicula*": Table S4

Table S4: KoT results on COI marker for the complete dataset (n=359). A group number is assigned to each individual as in KoT output (n=251). This clusterisation was inferred for individual with no COI available but belonging to the same populations. In case of ambiguity, the COI was not inferred, these individuals are labelled in black. The 12 corresponding colors were applied on haplowebs (Figure S2).

| Individuals | Groups | Statut | Color names on haplowebs | RGB |
|---|---|---|---|---|
| Mad2 | 1 | Output | olive | 128,128,0 |
| Mad3 | 1 | Inferred | olive | 128,128,0 |
| Mad6 | 1 | Inferred | olive | 128,128,0 |
| AA10 | 2 | Output | red | 255,0,0 |
| AA11 | 2 | Output | red | 255,0,0 |
| AA12 | 2 | Output | red | 255,0,0 |
| AA13 | 2 | Output | red | 255,0,0 |
| AA14 | 2 | Output | red | 255,0,0 |
| ccc7 | 2 | Inferred | red | 255,0,0 |
| CFG01 | 2 | Output | red | 255,0,0 |
| CFG02 | 2 | Output | red | 255,0,0 |
| CFG03 | 2 | Output | red | 255,0,0 |
| CFG04 | 2 | Output | red | 255,0,0 |
| CFG05 | 2 | Output | red | 255,0,0 |
| CFG06 | 2 | Output | red | 255,0,0 |
| CFG07 | 2 | Output | red | 255,0,0 |
| CFG08 | 2 | Output | red | 255,0,0 |
| CFG09 | 2 | Output | red | 255,0,0 |
| CFG10 | 2 | Output | red | 255,0,0 |
| CFG11 | 2 | Output | red | 255,0,0 |
| CFG12 | 2 | Output | red | 255,0,0 |
| CFG13 | 2 | Output | red | 255,0,0 |
| CFG15 | 2 | Output | red | 255,0,0 |
| CFY01 | 2 | Inferred | red | 255,0,0 |
| CFY02 | 2 | Inferred | red | 255,0,0 |
| CFY03 | 2 | Output | red | 255,0,0 |
| CFY04 | 2 | Output | red | 255,0,0 |
| CFY05 | 2 | Output | red | 255,0,0 |
| CFY06 | 2 | Inferred | red | 255,0,0 |
| CFY08 | 2 | Inferred | red | 255,0,0 |
| CFY09 | 2 | Output | red | 255,0,0 |
| CFY10 | 2 | Output | red | 255,0,0 |
| CFY11 | 2 | Output | red | 255,0,0 |
| CFY12 | 2 | Output | red | 255,0,0 |
| Db25 | 2 | Output | red | 255,0,0 |
| ggg2 | 2 | Inferred | red | 255,0,0 |
| Go1 | 2 | Inferred | red | 255,0,0 |
| KMT201 | 2 | Inferred | red | 255,0,0 |
| KMT202 | 2 | Inferred | red | 255,0,0 |
| KMT203 | 2 | Inferred | red | 255,0,0 |
| KMT204 | 2 | Inferred | red | 255,0,0 |
| KMT205 | 2 | Output | red | 255,0,0 |
| KMT206 | 2 | Inferred | red | 255,0,0 |
| KMT207 | 2 | Inferred | red | 255,0,0 |
| KMT208 | 2 | Output | red | 255,0,0 |

| | | | | |
|---|---|---|---|---|
| KMT209 | 2 | Inferred | red | 255,0,0 |
| KMT210 | 2 | Inferred | red | 255,0,0 |
| KMT211 | 2 | Inferred | red | 255,0,0 |
| KMT212 | 2 | Inferred | red | 255,0,0 |
| KMT213 | 2 | Output | red | 255,0,0 |
| KMT214 | 2 | Output | red | 255,0,0 |
| KMT215 | 2 | Output | red | 255,0,0 |
| KMT216 | 2 | Output | red | 255,0,0 |
| KMT217 | 2 | Output | red | 255,0,0 |
| KMT218 | 2 | Output | red | 255,0,0 |
| KMT219 | 2 | Output | red | 255,0,0 |
| KMT220 | 2 | Inferred | red | 255,0,0 |
| KMT221 | 2 | Output | red | 255,0,0 |
| KMT222 | 2 | Output | red | 255,0,0 |
| KMT223 | 2 | Inferred | red | 255,0,0 |
| KMT224 | 2 | Output | red | 255,0,0 |
| LB13 | 2 | Output | red | 255,0,0 |
| LB17 | 2 | Output | red | 255,0,0 |
| LB19 | 2 | Output | red | 255,0,0 |
| LB31 | 2 | Output | red | 255,0,0 |
| LB40 | 2 | Output | red | 255,0,0 |
| LB43 | 2 | Output | red | 255,0,0 |
| LB66 | 2 | Output | red | 255,0,0 |
| LB68 | 2 | Output | red | 255,0,0 |
| LB69 | 2 | Output | red | 255,0,0 |
| LB72 | 2 | Output | red | 255,0,0 |
| LB76 | 2 | Output | red | 255,0,0 |
| LB80 | 2 | Output | red | 255,0,0 |
| LB91 | 2 | Output | red | 255,0,0 |
| qq1 | 2 | Inferred | red | 255,0,0 |
| Su1 | 2 | Output | red | 255,0,0 |
| Su2 | 2 | Output | red | 255,0,0 |
| Su3 | 2 | Output | red | 255,0,0 |
| Su4 | 2 | Output | red | 255,0,0 |
| Su5 | 2 | Output | red | 255,0,0 |
| xx11 | 2 | Inferred | red | 255,0,0 |
| 170096 | 2 | Output | red | 255,0,0 |
| BA1 | 3 | Output | blue | 0,0,255 |
| BA2 | 3 | Inferred | blue | 0,0,255 |
| BA3 | 3 | Inferred | blue | 0,0,255 |
| BA4 | 3 | Inferred | blue | 0,0,255 |
| BA5 | 3 | Inferred | blue | 0,0,255 |
| BA6 | 3 | Output | blue | 0,0,255 |
| BA7 | 3 | Inferred | blue | 0,0,255 |
| BA8 | 3 | Inferred | blue | 0,0,255 |
| FU1 | 3 | Output | blue | 0,0,255 |
| FU4 | 3 | Inferred | blue | 0,0,255 |
| FU5 | 3 | Inferred | blue | 0,0,255 |
| FU6 | 3 | Inferred | blue | 0,0,255 |
| Muj02 | 3 | Inferred | blue | 0,0,255 |
| Muj08 | 3 | Inferred | blue | 0,0,255 |

| | | | | |
|---|---|---|---|---|
| Muj10 | 3 | Inferred | blue | 0,0,255 |
| 190768 | 4 | Output | violet | 238,130,238 |
| 190852 | 4 | Output | violet | 238,130,238 |
| 191042 | 5 | Output | purple | 128,0,128 |
| CS04 | 6 | Output | green | 0,128,0 |
| CS06 | 6 | Output | green | 0,128,0 |
| CS09 | 6 | Output | green | 0,128,0 |
| CS10 | 6 | Output | green | 0,128,0 |
| CS11 | 6 | Inferred | green | 0,128,0 |
| CS12 | 6 | Output | green | 0,128,0 |
| CS13 | 6 | Output | green | 0,128,0 |
| CS14 | 6 | Output | green | 0,128,0 |
| CS15 | 6 | Output | green | 0,128,0 |
| CS18 | 6 | Inferred | green | 0,128,0 |
| CS19 | 6 | Output | green | 0,128,0 |
| CS21 | 6 | Output | green | 0,128,0 |
| CS22 | 6 | Output | green | 0,128,0 |
| CS23 | 6 | Output | green | 0,128,0 |
| CS24 | 6 | Output | green | 0,128,0 |
| CS25 | 6 | Output | green | 0,128,0 |
| CS26 | 6 | Output | green | 0,128,0 |
| CS27 | 6 | Output | green | 0,128,0 |
| CS31 | 6 | Output | green | 0,128,0 |
| CS32 | 6 | Output | green | 0,128,0 |
| LB36 | 6 | Output | green | 0,128,0 |
| LB75 | 6 | Output | green | 0,128,0 |
| LB77 | 6 | Output | green | 0,128,0 |
| ZA02 | 7 | Inferred | brown | 165,42,42 |
| ZA03 | 7 | Output | brown | 165,42,42 |
| ZA04 | 7 | Inferred | brown | 165,42,42 |
| ZA05 | 7 | Output | brown | 165,42,42 |
| ZA06 | 7 | Output | brown | 165,42,42 |
| ZA07 | 7 | Output | brown | 165,42,42 |
| ZA08 | 7 | Inferred | brown | 165,42,42 |
| ZA09 | 7 | Output | brown | 165,42,42 |
| ZA10 | 7 | Output | brown | 165,42,42 |
| ZA11 | 7 | Output | brown | 165,42,42 |
| ZA12 | 7 | Output | brown | 165,42,42 |
| ZA13 | 7 | Output | brown | 165,42,42 |
| ZA14 | 7 | Inferred | brown | 165,42,42 |
| ZA15 | 7 | Output | brown | 165,42,42 |
| ZA16 | 7 | Output | brown | 165,42,42 |
| ZA17 | 7 | Output | brown | 165,42,42 |
| ZA18 | 7 | Output | brown | 165,42,42 |
| ZA19 | 7 | Output | brown | 165,42,42 |
| ZA20 | 7 | Inferred | brown | 165,42,42 |
| ZA01 | 7 | Output | brown | 165,42,42 |
| C1 | 8 | Output | orange | 255,165,0 |
| C2 | 8 | Output | orange | 255,165,0 |
| C3 | 8 | Output | orange | 255,165,0 |
| C4 | 8 | Output | orange | 255,165,0 |

| | | | |
|---|---|---|---|
| fff2 | 8 Inferred | orange | 255,165,0 |
| Sa22 | 8 Output | orange | 255,165,0 |
| Sa23 | 8 Output | orange | 255,165,0 |
| Sa24 | 8 Output | orange | 255,165,0 |
| Sa25 | 8 Output | orange | 255,165,0 |
| Sa26 | 8 Output | orange | 255,165,0 |
| Sa72 | 8 Output | orange | 255,165,0 |
| Sa73 | 8 Output | orange | 255,165,0 |
| Sa74 | 8 Output | orange | 255,165,0 |
| Sa75 | 8 Output | orange | 255,165,0 |
| U3 | 8 Inferred | orange | 255,165,0 |
| U4 | 8 Inferred | orange | 255,165,0 |
| AB11 | 9 Output | cyan | 0,255,255 |
| AB12 | 9 Output | cyan | 0,255,255 |
| AB13 | 9 Output | cyan | 0,255,255 |
| AB14 | 9 Output | cyan | 0,255,255 |
| AB15 | 9 Output | cyan | 0,255,255 |
| AB16 | 9 Output | cyan | 0,255,255 |
| CFG14 | 9 Inferred | cyan | 0,255,255 |
| CFG16 | 9 Output | cyan | 0,255,255 |
| CFG17 | 9 Output | cyan | 0,255,255 |
| CFG18 | 9 Output | cyan | 0,255,255 |
| CFG19 | 9 Output | cyan | 0,255,255 |
| CFG20 | 9 Output | cyan | 0,255,255 |
| CFG21 | 9 Output | cyan | 0,255,255 |
| CFG22 | 9 Output | cyan | 0,255,255 |
| CFG23 | 9 Output | cyan | 0,255,255 |
| CFY07 | 9 Output | cyan | 0,255,255 |
| Cl201 | 9 Inferred | cyan | 0,255,255 |
| Cl202 | 9 Inferred | cyan | 0,255,255 |
| Cl204 | 9 Output | cyan | 0,255,255 |
| Cl205 | 9 Output | cyan | 0,255,255 |
| Cl206 | 9 Output | cyan | 0,255,255 |
| Cl207 | 9 Output | cyan | 0,255,255 |
| Cl208 | 9 Output | cyan | 0,255,255 |
| Cl209 | 9 Inferred | cyan | 0,255,255 |
| Cl210 | 9 Output | cyan | 0,255,255 |
| Cl211 | 9 Output | cyan | 0,255,255 |
| Cl212 | 9 Output | cyan | 0,255,255 |
| Cl213 | 9 Output | cyan | 0,255,255 |
| Cl214 | 9 Output | cyan | 0,255,255 |
| Cl215 | 9 Output | cyan | 0,255,255 |
| Cl216 | 9 Output | cyan | 0,255,255 |
| Cl217 | 9 Inferred | cyan | 0,255,255 |
| Cl218 | 9 Output | cyan | 0,255,255 |
| Cl220 | 9 Output | cyan | 0,255,255 |
| Cl221 | 9 Output | cyan | 0,255,255 |
| Cl222 | 9 Output | cyan | 0,255,255 |
| Cl223 | 9 Output | cyan | 0,255,255 |
| Cl224 | 9 Output | cyan | 0,255,255 |
| Db22 | 9 Output | cyan | 0,255,255 |

| | | | | |
|---|---|---|---|---|
| Db23 | 9 | Output | cyan | 0,255,255 |
| Db24 | 9 | Output | cyan | 0,255,255 |
| Db26 | 9 | Output | cyan | 0,255,255 |
| EHM101 | 9 | Inferred | cyan | 0,255,255 |
| EHM102 | 9 | Output | cyan | 0,255,255 |
| EHM103 | 9 | Output | cyan | 0,255,255 |
| EHM104 | 9 | Output | cyan | 0,255,255 |
| EHM105 | 9 | Output | cyan | 0,255,255 |
| EHM106 | 9 | Output | cyan | 0,255,255 |
| EHM107 | 9 | Output | cyan | 0,255,255 |
| EHM108 | 9 | Output | cyan | 0,255,255 |
| EHM109 | 9 | Output | cyan | 0,255,255 |
| EHM110 | 9 | Output | cyan | 0,255,255 |
| EHM111 | 9 | Output | cyan | 0,255,255 |
| EHM112 | 9 | Output | cyan | 0,255,255 |
| EHM113 | 9 | Output | cyan | 0,255,255 |
| EHM114 | 9 | Inferred | cyan | 0,255,255 |
| EHM115 | 9 | Inferred | cyan | 0,255,255 |
| EHM116 | 9 | Inferred | cyan | 0,255,255 |
| EHM119 | 9 | Output | cyan | 0,255,255 |
| EHM120 | 9 | Output | cyan | 0,255,255 |
| EHM121 | 9 | Output | cyan | 0,255,255 |
| EHM122 | 9 | Output | cyan | 0,255,255 |
| EHM123 | 9 | Output | cyan | 0,255,255 |
| EHM124 | 9 | Output | cyan | 0,255,255 |
| LB01 | 9 | Output | cyan | 0,255,255 |
| LB02 | 9 | Output | cyan | 0,255,255 |
| LB04 | 9 | Output | cyan | 0,255,255 |
| LB05 | 9 | Output | cyan | 0,255,255 |
| LB08 | 9 | Output | cyan | 0,255,255 |
| LB11 | 9 | Output | cyan | 0,255,255 |
| LB12 | 9 | Output | cyan | 0,255,255 |
| LB14 | 9 | Output | cyan | 0,255,255 |
| LB16 | 9 | Output | cyan | 0,255,255 |
| LB18 | 9 | Output | cyan | 0,255,255 |
| LB20 | 9 | Output | cyan | 0,255,255 |
| LB22 | 9 | Output | cyan | 0,255,255 |
| LB42 | 9 | Output | cyan | 0,255,255 |
| LB44 | 9 | Output | cyan | 0,255,255 |
| LB45 | 9 | Output | cyan | 0,255,255 |
| LB47 | 9 | Output | cyan | 0,255,255 |
| LB48 | 9 | Output | cyan | 0,255,255 |
| LB49 | 9 | Output | cyan | 0,255,255 |
| LB58 | 9 | Output | cyan | 0,255,255 |
| LB59 | 9 | Output | cyan | 0,255,255 |
| LB62 | 9 | Output | cyan | 0,255,255 |
| LB64 | 9 | Output | cyan | 0,255,255 |
| LB67 | 9 | Output | cyan | 0,255,255 |
| LB71 | 9 | Output | cyan | 0,255,255 |
| LB74 | 9 | Output | cyan | 0,255,255 |
| LB81 | 9 | Output | cyan | 0,255,255 |

| | | | | |
|---|---|---|---|---|
| LB92 | 9 | Output | cyan | 0,255,255 |
| LB98 | 9 | Output | cyan | 0,255,255 |
| rr1 | 9 | Inferred | cyan | 0,255,255 |
| yy11 | 9 | Inferred | cyan | 0,255,255 |
| yy12 | 9 | Output | cyan | 0,255,255 |
| CR01 | 10 | Output | magenta | 255,0,255 |
| CR02 | 10 | Output | magenta | 255,0,255 |
| CR03 | 10 | Output | magenta | 255,0,255 |
| CR04 | 10 | Output | magenta | 255,0,255 |
| CR05 | 10 | Output | magenta | 255,0,255 |
| CR06 | 10 | Inferred | magenta | 255,0,255 |
| CR09 | 10 | Inferred | magenta | 255,0,255 |
| CR12 | 10 | Inferred | magenta | 255,0,255 |
| CR13 | 10 | Output | magenta | 255,0,255 |
| CR14 | 10 | Inferred | magenta | 255,0,255 |
| CR15 | 10 | Output | magenta | 255,0,255 |
| CR16 | 10 | Inferred | magenta | 255,0,255 |
| CR17 | 10 | Output | magenta | 255,0,255 |
| CR18 | 10 | Inferred | magenta | 255,0,255 |
| CR19 | 10 | Inferred | magenta | 255,0,255 |
| CR20 | 10 | Inferred | magenta | 255,0,255 |
| CR21 | 10 | Inferred | magenta | 255,0,255 |
| CR22 | 10 | Output | magenta | 255,0,255 |
| CR23 | 10 | Inferred | magenta | 255,0,255 |
| CR24 | 10 | Output | magenta | 255,0,255 |
| CR25 | 10 | Inferred | magenta | 255,0,255 |
| CR26 | 10 | Output | magenta | 255,0,255 |
| CRB01 | 10 | Output | magenta | 255,0,255 |
| CRB02 | 10 | Inferred | magenta | 255,0,255 |
| CRB03 | 10 | Inferred | magenta | 255,0,255 |
| CRB04 | 10 | Output | magenta | 255,0,255 |
| CRB05 | 10 | Output | magenta | 255,0,255 |
| CRB06 | 10 | Inferred | magenta | 255,0,255 |
| CRB07 | 10 | Inferred | magenta | 255,0,255 |
| CRB09 | 10 | Output | magenta | 255,0,255 |
| CRB10 | 10 | Output | magenta | 255,0,255 |
| CRB11 | 10 | Output | magenta | 255,0,255 |
| CRB12 | 10 | Output | magenta | 255,0,255 |
| CRB13 | 10 | Output | magenta | 255,0,255 |
| CRB14 | 10 | Output | magenta | 255,0,255 |
| CRB15 | 10 | Output | magenta | 255,0,255 |
| CRB16 | 10 | Output | magenta | 255,0,255 |
| CRB17 | 10 | Inferred | magenta | 255,0,255 |
| CRB18 | 10 | Inferred | magenta | 255,0,255 |
| CRB19 | 10 | Inferred | magenta | 255,0,255 |
| CRB20 | 10 | Inferred | magenta | 255,0,255 |
| CRB21 | 10 | Inferred | magenta | 255,0,255 |
| Vt01 | 10 | Output | magenta | 255,0,255 |
| Vt02 | 10 | Output | magenta | 255,0,255 |
| Vt03 | 10 | Output | magenta | 255,0,255 |
| Vt05 | 10 | Inferred | magenta | 255,0,255 |

| | | | | |
|---|---|---|---|---|
| Vt06 | 10 | Inferred | magenta | 255,0,255 |
| Vt07 | 10 | Output | magenta | 255,0,255 |
| Vt08 | 10 | Inferred | magenta | 255,0,255 |
| Vt09 | 10 | Inferred | magenta | 255,0,255 |
| Vt10 | 10 | Output | magenta | 255,0,255 |
| Vt11 | 10 | Output | magenta | 255,0,255 |
| Vt12 | 10 | Output | magenta | 255,0,255 |
| Vt13 | 10 | Output | magenta | 255,0,255 |
| Vt14 | 10 | Inferred | magenta | 255,0,255 |
| Vt15 | 10 | Output | magenta | 255,0,255 |
| Vt16 | 10 | Output | magenta | 255,0,255 |
| Vt17 | 10 | Output | magenta | 255,0,255 |
| Vt19 | 10 | Output | magenta | 255,0,255 |
| Vt20 | 10 | Output | magenta | 255,0,255 |
| CJ01 | 11 | Output | lightslategrey | 119,136,153 |
| CJ02 | 11 | Inferred | lightslategrey | 119,136,153 |
| CJ03 | 11 | Output | lightslategrey | 119,136,153 |
| CJ04 | 11 | Output | lightslategrey | 119,136,153 |
| CJ05 | 11 | Output | lightslategrey | 119,136,153 |
| CJ06 | 11 | Inferred | lightslategrey | 119,136,153 |
| CJ07 | 11 | Output | lightslategrey | 119,136,153 |
| CJ08 | 11 | Inferred | lightslategrey | 119,136,153 |
| CJ09 | 11 | Output | lightslategrey | 119,136,153 |
| CJ10 | 11 | Output | lightslategrey | 119,136,153 |
| CJ11 | 11 | Output | lightslategrey | 119,136,153 |
| CJ12 | 11 | Output | lightslategrey | 119,136,153 |
| CJ13 | 11 | Inferred | lightslategrey | 119,136,153 |
| CJ14 | 11 | Inferred | lightslategrey | 119,136,153 |
| CJ15 | 11 | Output | lightslategrey | 119,136,153 |
| CJ16 | 11 | Inferred | lightslategrey | 119,136,153 |
| CJ17 | 11 | Inferred | lightslategrey | 119,136,153 |
| CJ18 | 11 | Output | lightslategrey | 119,136,153 |
| CJ19 | 11 | Output | lightslategrey | 119,136,153 |
| CJ20 | 11 | Inferred | lightslategrey | 119,136,153 |
| CJ21 | 11 | Output | lightslategrey | 119,136,153 |
| CJ22 | 11 | Inferred | lightslategrey | 119,136,153 |
| Jp01 | 11 | Output | lightslategrey | 119,136,153 |
| Jp02 | 11 | Output | lightslategrey | 119,136,153 |
| Jp03 | 11 | Output | lightslategrey | 119,136,153 |
| Jp04 | 11 | Inferred | lightslategrey | 119,136,153 |
| Jp06 | 11 | Inferred | lightslategrey | 119,136,153 |
| Jp08 | 11 | Inferred | lightslategrey | 119,136,153 |
| Jp10 | 11 | Output | lightslategrey | 119,136,153 |
| Ka1 | 11 | Output | lightslategrey | 119,136,153 |
| Ka2 | 11 | Output | lightslategrey | 119,136,153 |
| Ka3 | 11 | Output | lightslategrey | 119,136,153 |
| Ka5 | 11 | Output | lightslategrey | 119,136,153 |
| 103026 | NA | Not-inferred | black | 0,0,0 |
| 103027 | NA | Not-inferred | black | 0,0,0 |
| 266690 | NA | Not-inferred | black | 0,0,0 |
| 266691 | NA | Not-inferred | black | 0,0,0 |

| | | | | |
|---|---|---|---|---|
| Hw2 | NA | Not-inferred | black | 0,0,0 |
| Hw3 | NA | Not-inferred | black | 0,0,0 |
| Hw4 | NA | Not-inferred | black | 0,0,0 |
| Hw5 | NA | Not-inferred | black | 0,0,0 |
| Hw6 | NA | Not-inferred | black | 0,0,0 |
| LB06 | NA | Not-inferred | black | 0,0,0 |
| Wa1 | NA | Not-inferred | black | 0,0,0 |
| Wa2 | NA | Not-inferred | black | 0,0,0 |
