## Supplementary material for "Substantial genetic mixing among sexual and androgenetic lineages within the clam genus *Corbicula*": Table S5

Table S5: KoT results on COI marker for the filtered dataset (n=141). A group number is assigned to each individual as in KoT output. The 8 corresponding colors were applied on haplowebs (Figure 2).

| Individuals | Groups | Color names on haplowebs | RGB |
|---|---|---|---|
| AA11 | 1 | red | 255,0,0 |
| AA12 | 1 | red | 255,0,0 |
| AA13 | 1 | red | 255,0,0 |
| CFG01 | 1 | red | 255,0,0 |
| CFG02 | 1 | red | 255,0,0 |
| CFG03 | 1 | red | 255,0,0 |
| CFG04 | 1 | red | 255,0,0 |
| CFG05 | 1 | red | 255,0,0 |
| CFG06 | 1 | red | 255,0,0 |
| CFG07 | 1 | red | 255,0,0 |
| CFG08 | 1 | red | 255,0,0 |
| CFG09 | 1 | red | 255,0,0 |
| CFG10 | 1 | red | 255,0,0 |
| CFG11 | 1 | red | 255,0,0 |
| CFG12 | 1 | red | 255,0,0 |
| CFG13 | 1 | red | 255,0,0 |
| CFG15 | 1 | red | 255,0,0 |
| CFY03 | 1 | red | 255,0,0 |
| CFY04 | 1 | red | 255,0,0 |
| CFY05 | 1 | red | 255,0,0 |
| CFY09 | 1 | red | 255,0,0 |
| CFY10 | 1 | red | 255,0,0 |
| CFY11 | 1 | red | 255,0,0 |
| CFY12 | 1 | red | 255,0,0 |
| Db25 | 1 | red | 255,0,0 |
| KMT205 | 1 | red | 255,0,0 |
| KMT208 | 1 | red | 255,0,0 |
| KMT213 | 1 | red | 255,0,0 |
| KMT214 | 1 | red | 255,0,0 |
| KMT215 | 1 | red | 255,0,0 |
| KMT216 | 1 | red | 255,0,0 |
| KMT217 | 1 | red | 255,0,0 |
| KMT218 | 1 | red | 255,0,0 |
| KMT219 | 1 | red | 255,0,0 |
| KMT221 | 1 | red | 255,0,0 |
| KMT222 | 1 | red | 255,0,0 |
| KMT224 | 1 | red | 255,0,0 |
| ZA05 | 2 | brown | 165,42,42 |
| ZA06 | 2 | brown | 165,42,42 |
| ZA09 | 2 | brown | 165,42,42 |
| ZA10 | 2 | brown | 165,42,42 |
| ZA11 | 2 | brown | 165,42,42 |
| ZA12 | 2 | brown | 165,42,42 |
| ZA13 | 2 | brown | 165,42,42 |
| ZA15 | 2 | brown | 165,42,42 |
| ZA16 | 2 | brown | 165,42,42 |
| ZA17 | 2 | brown | 165,42,42 |

| | | | |
|---|---|---|---|
| ZA18 | 2 | brown | 165,42,42 |
| ZA19 | 2 | brown | 165,42,42 |
| ZA01 | 2 | brown | 165,42,42 |
| Sa22 | 3 | orange | 255,165,0 |
| C1 | 3 | orange | 255,165,0 |
| C2 | 3 | orange | 255,165,0 |
| C3 | 3 | orange | 255,165,0 |
| C4 | 3 | orange | 255,165,0 |
| Db22 | 4 | green | 146,208,80 |
| Db23 | 4 | green | 146,208,80 |
| Db24 | 4 | green | 146,208,80 |
| Db26 | 4 | green | 146,208,80 |
| AB11 | 5 | cyan | 0,255,255 |
| AB13 | 5 | cyan | 0,255,255 |
| AB14 | 5 | cyan | 0,255,255 |
| AB15 | 5 | cyan | 0,255,255 |
| CFG16 | 5 | cyan | 0,255,255 |
| CFG17 | 5 | cyan | 0,255,255 |
| CFG18 | 5 | cyan | 0,255,255 |
| CFG19 | 5 | cyan | 0,255,255 |
| CFG20 | 5 | cyan | 0,255,255 |
| CFG21 | 5 | cyan | 0,255,255 |
| CFG22 | 5 | cyan | 0,255,255 |
| CFG23 | 5 | cyan | 0,255,255 |
| EHM102 | 5 | cyan | 0,255,255 |
| EHM103 | 5 | cyan | 0,255,255 |
| EHM104 | 5 | cyan | 0,255,255 |
| EHM105 | 5 | cyan | 0,255,255 |
| EHM106 | 5 | cyan | 0,255,255 |
| EHM107 | 5 | cyan | 0,255,255 |
| EHM108 | 5 | cyan | 0,255,255 |
| EHM109 | 5 | cyan | 0,255,255 |
| EHM110 | 5 | cyan | 0,255,255 |
| EHM111 | 5 | cyan | 0,255,255 |
| EHM112 | 5 | cyan | 0,255,255 |
| EHM113 | 5 | cyan | 0,255,255 |
| EHM119 | 5 | cyan | 0,255,255 |
| EHM120 | 5 | cyan | 0,255,255 |
| EHM121 | 5 | cyan | 0,255,255 |
| EHM122 | 5 | cyan | 0,255,255 |
| yy12 | 5 | cyan | 0,255,255 |
| CFY07 | 6 | blue | 0,0,255 |
| Cl204 | 6 | blue | 0,0,255 |
| Cl205 | 6 | blue | 0,0,255 |
| Cl206 | 6 | blue | 0,0,255 |
| Cl207 | 6 | blue | 0,0,255 |
| Cl208 | 6 | blue | 0,0,255 |
| Cl210 | 6 | blue | 0,0,255 |
| Cl211 | 6 | blue | 0,0,255 |
| Cl212 | 6 | blue | 0,0,255 |
| Cl213 | 6 | blue | 0,0,255 |

| | | |
|---|---|---|
| Cl214 | 6 blue | 0,0,255 |
| Cl215 | 6 blue | 0,0,255 |
| Cl216 | 6 blue | 0,0,255 |
| Cl218 | 6 blue | 0,0,255 |
| Cl220 | 6 blue | 0,0,255 |
| Cl221 | 6 blue | 0,0,255 |
| Cl222 | 6 blue | 0,0,255 |
| Cl223 | 6 blue | 0,0,255 |
| Cl224 | 6 blue | 0,0,255 |
| Vt10 | 7 magenta | 255,0,255 |
| Vt13 | 7 magenta | 255,0,255 |
| CR02 | 7 magenta | 255,0,255 |
| CR01 | 7 magenta | 255,0,255 |
| CR05 | 7 magenta | 255,0,255 |
| CRB04 | 7 magenta | 255,0,255 |
| Vt01 | 7 magenta | 255,0,255 |
| Vt03 | 7 magenta | 255,0,255 |
| Vt12 | 7 magenta | 255,0,255 |
| CR03 | 7 magenta | 255,0,255 |
| CR13 | 7 magenta | 255,0,255 |
| CR15 | 7 magenta | 255,0,255 |
| CR17 | 7 magenta | 255,0,255 |
| CR24 | 7 magenta | 255,0,255 |
| CRB05 | 7 magenta | 255,0,255 |
| CRB12 | 7 magenta | 255,0,255 |
| CRB14 | 7 magenta | 255,0,255 |
| Vt02 | 7 magenta | 255,0,255 |
| Vt11 | 7 magenta | 255,0,255 |
| Vt15 | 7 magenta | 255,0,255 |
| Vt16 | 7 magenta | 255,0,255 |
| Vt17 | 7 magenta | 255,0,255 |
| Vt19 | 7 magenta | 255,0,255 |
| CR04 | 7 magenta | 255,0,255 |
| CJ09 | 8 lightslategrey | 119,136,153 |
| CJ10 | 8 lightslategrey | 119,136,153 |
| CJ11 | 8 lightslategrey | 119,136,153 |
| CJ07 | 8 lightslategrey | 119,136,153 |
| CJ18 | 8 lightslategrey | 119,136,153 |
| CJ19 | 8 lightslategrey | 119,136,153 |
| CJ03 | 8 lightslategrey | 119,136,153 |
| CJ04 | 8 lightslategrey | 119,136,153 |
| CJ15 | 8 lightslategrey | 119,136,153 |
| CJ12 | 8 lightslategrey | 119,136,153 |
