## Supplementary material for "Substantial genetic mixing among sexual and androgenetic lineages within the clam genus *Corbicula*": Figure S1

**Figure S1.** Schematic description of the haploweb method (Flot *et al.* 2010).

Let us imagine a hypothetical sampling of 6 diploid individuals, among which five are heterozygous and one is homozygous for a marker of interest, with each allele/haplotype sequence represented by a specific color below.

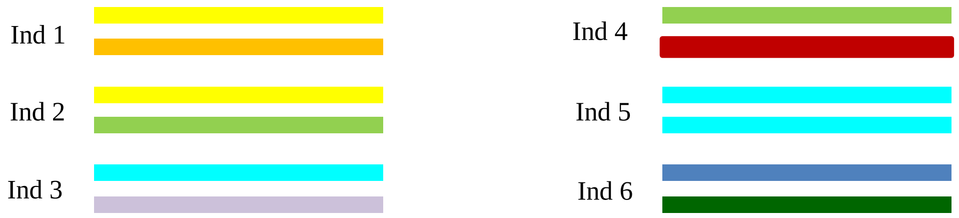


A first step is to draw a median-joining network of the alleles (*i.e.,* a distance-based network representing sequence similarities), in which each haplotype is represented by a circle of diameter proportional to the number of individuals harboring this haplotype in the dataset.

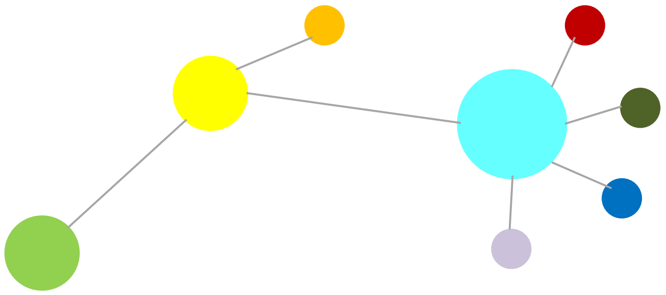

As a second step, connections are added between each pair of alleles found co-occurring in heterozygous individuals.

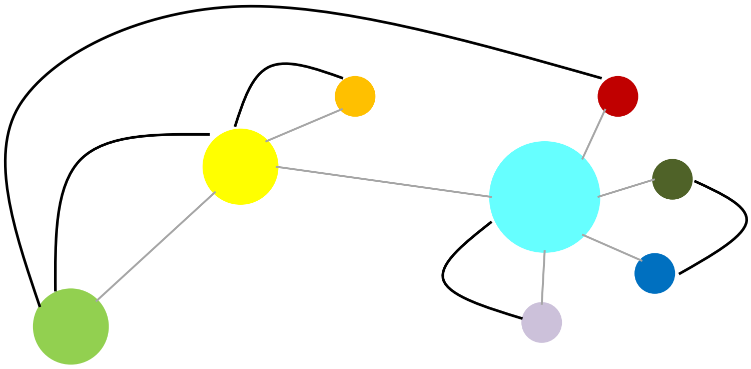

Based on allele sharing, the last step is to delimit fields for recombination (FFRs *sensu* Doyle 1995), *i.e.* groups of individuals belonging to the same allelic pool.

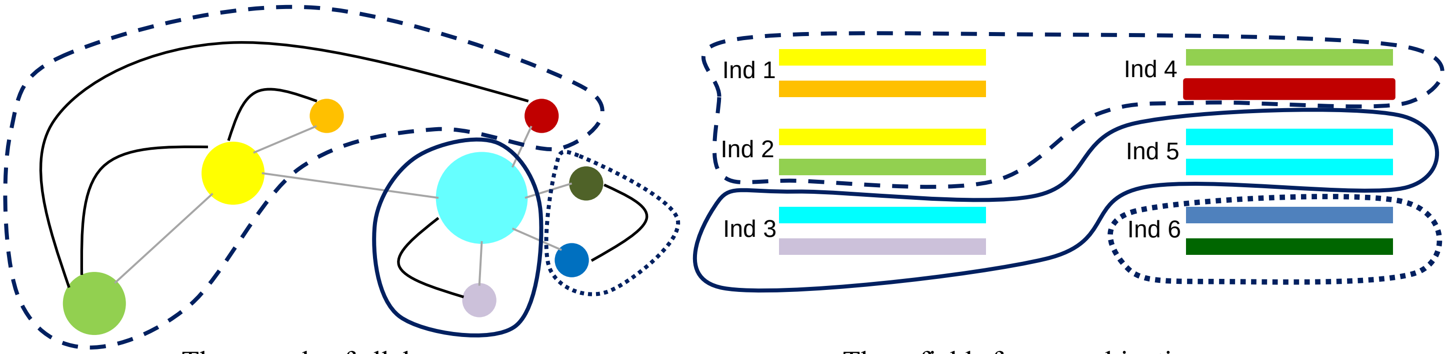


Three pools of alleles

Three fields for recombination
