## Supplementary material for "Substantial genetic mixing among sexual and androgenetic lineages within the clam genus *Corbicula*": Figure S2

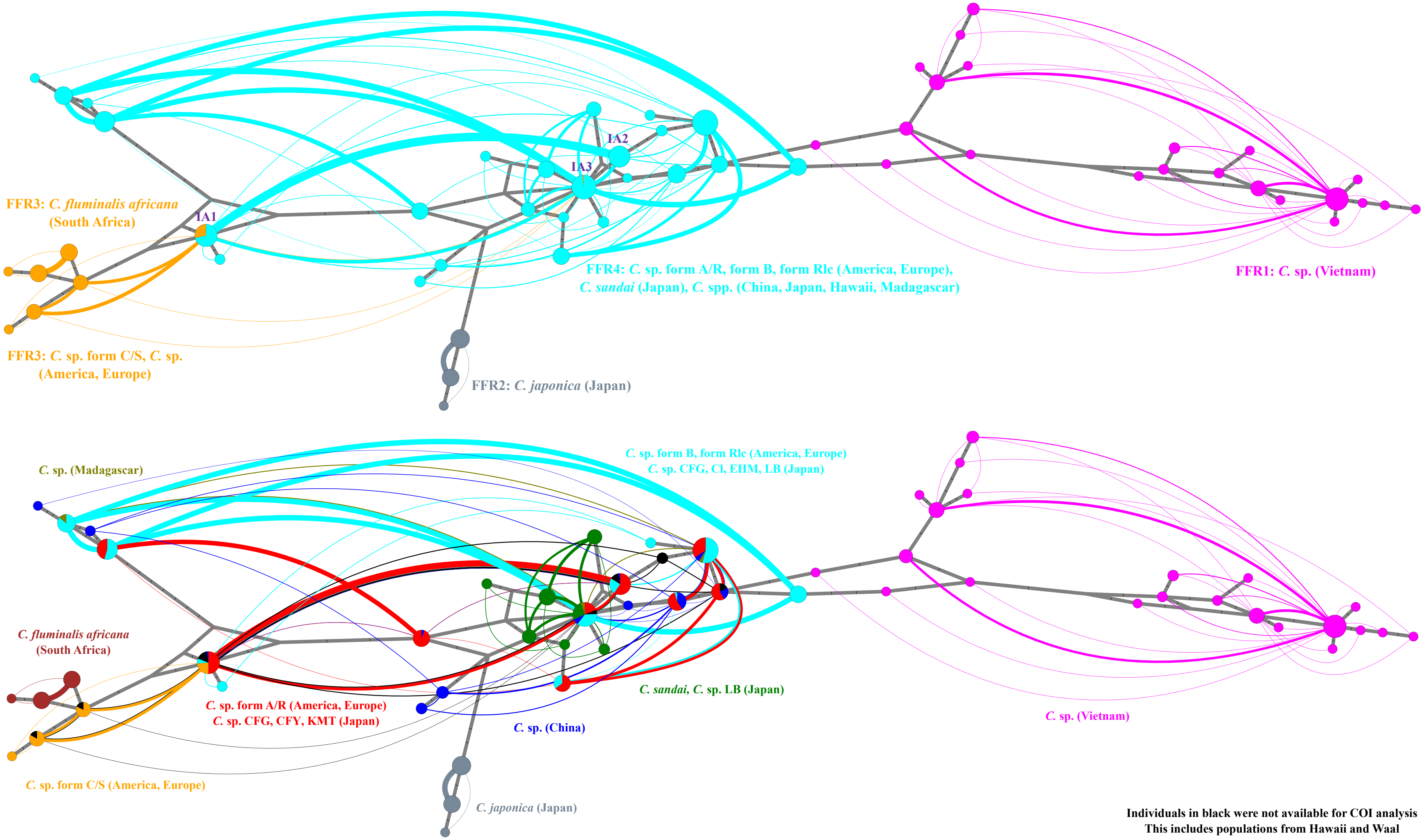


Figure S2a: 28S haploweb (n = 266).

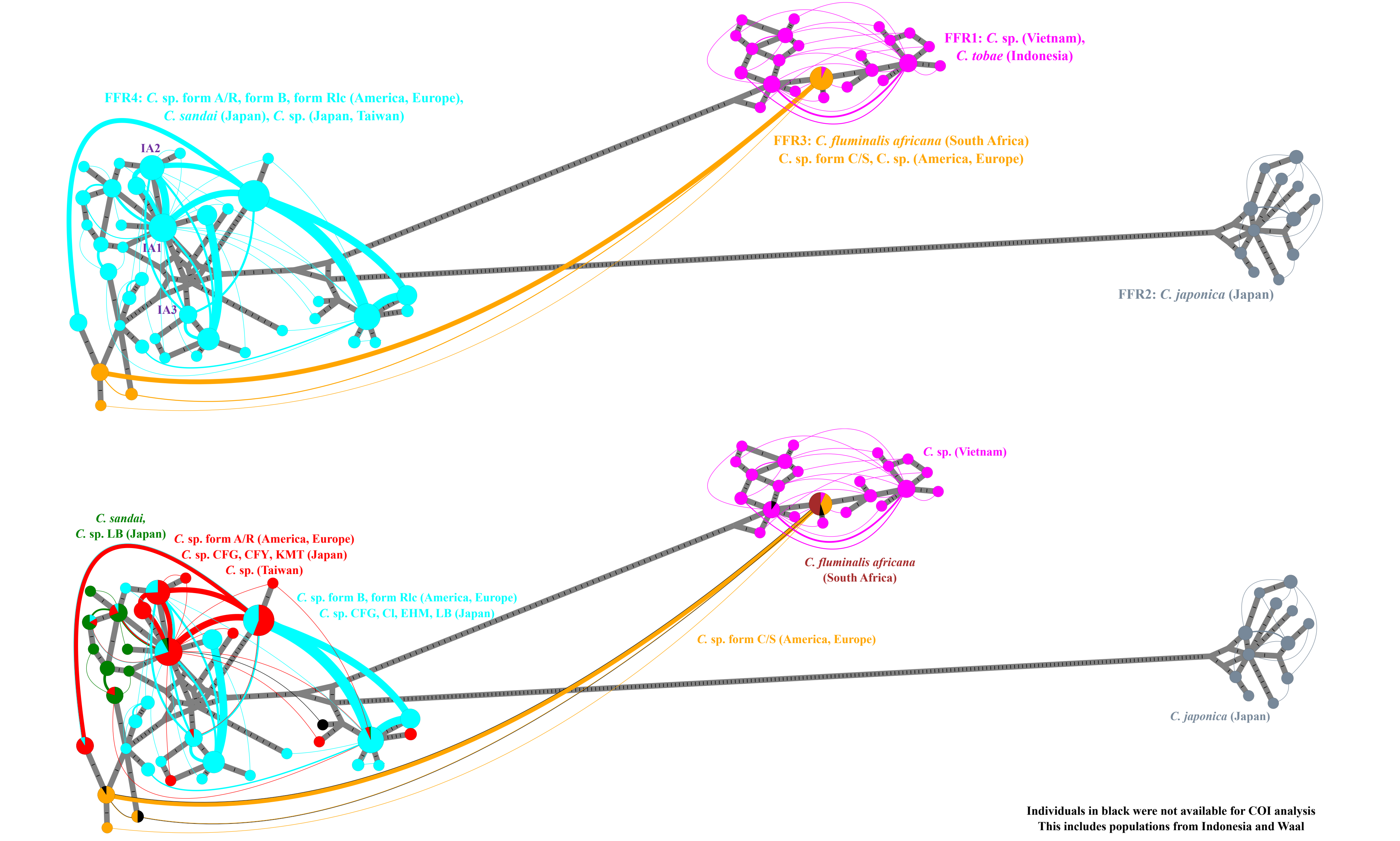


Figure S2b: alpha-amylase haploweb (n=237).

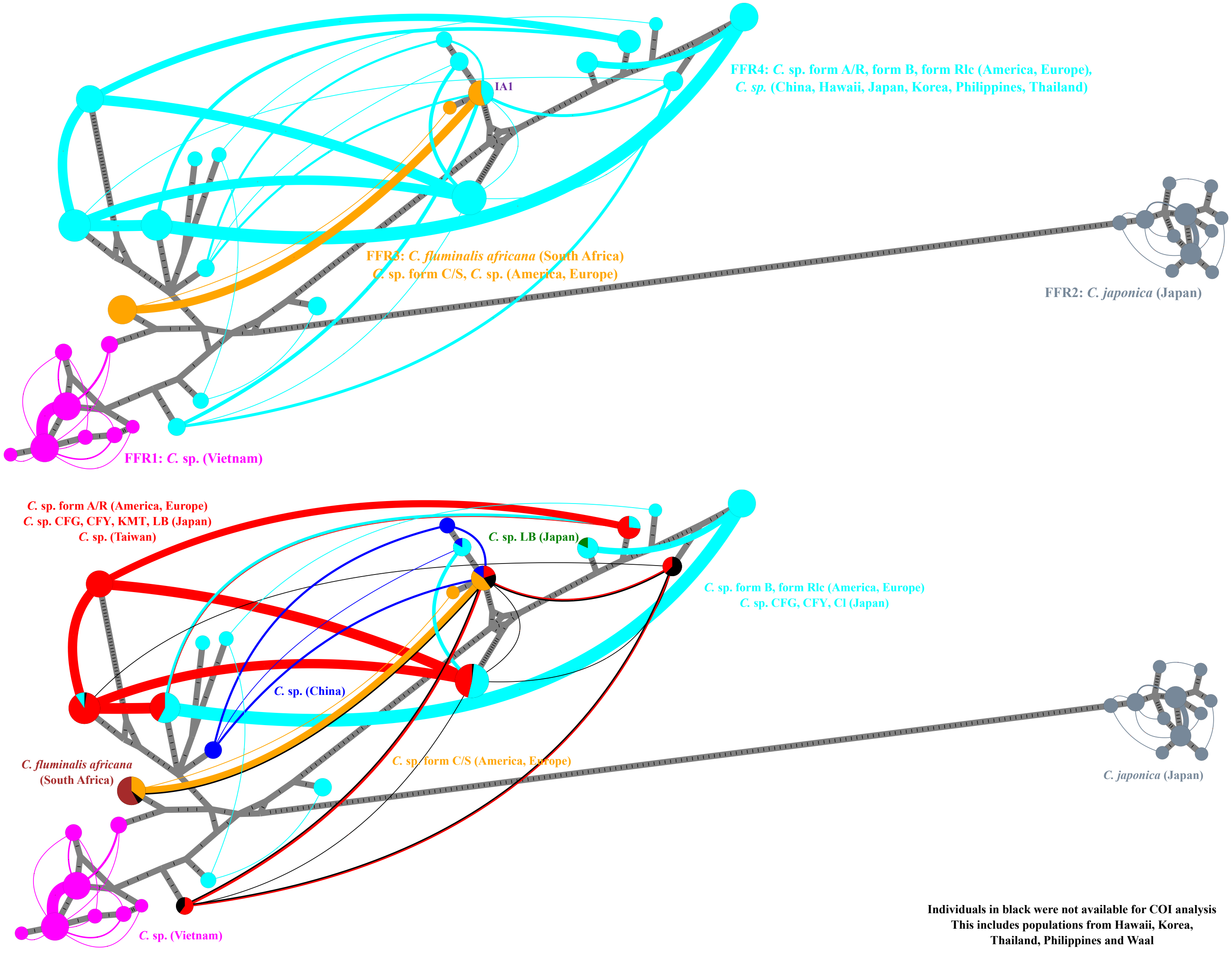


Figure S2c: ATPS haploweb (n=248).

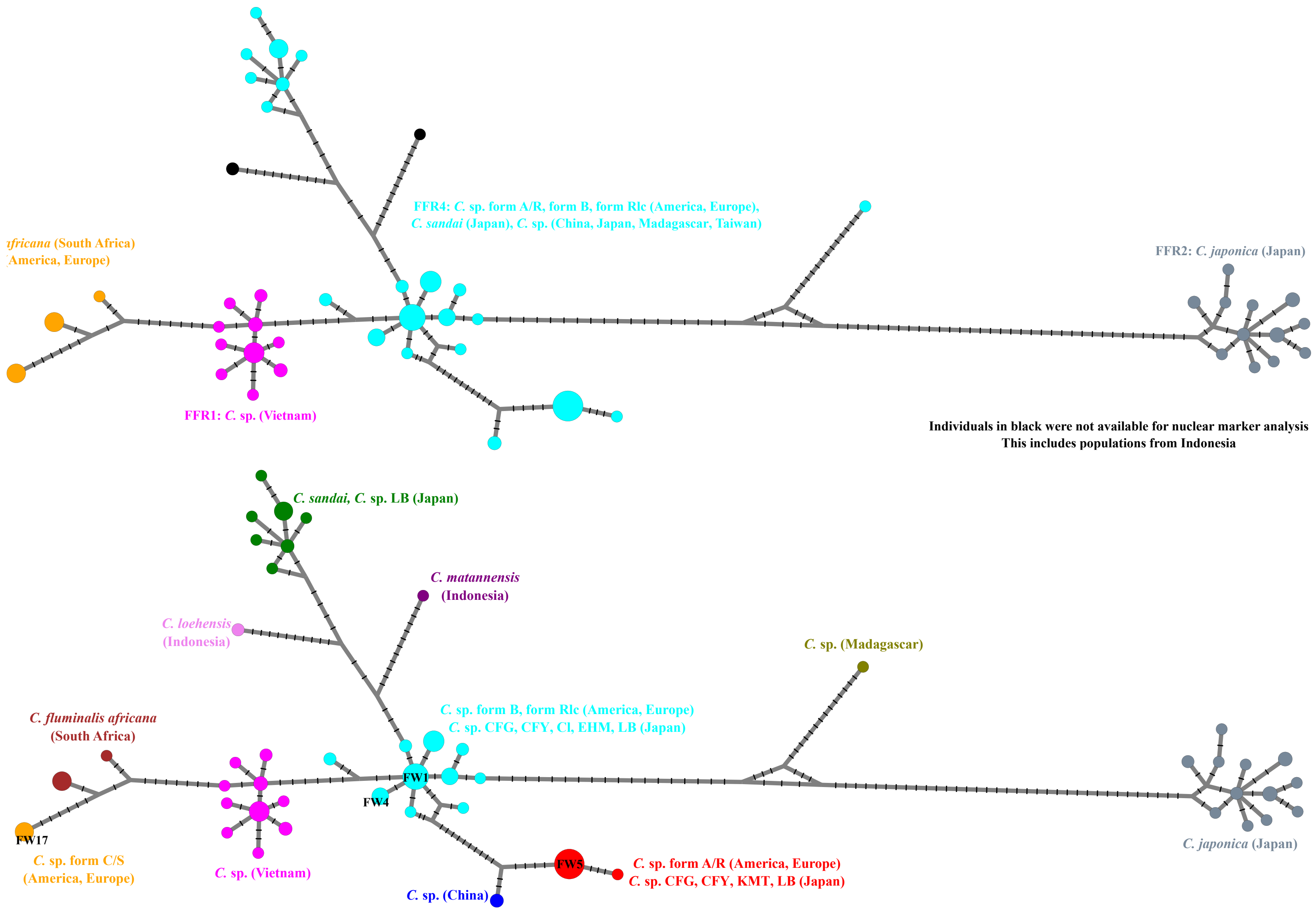


Figure S2d: COI haplonet (n = 251).
